# Evaluating the reliability of functional near-infrared spectroscopy data in the context of a reasoning paradigm

**DOI:** 10.64898/2026.01.16.699971

**Authors:** Patrick G. Kelly, Heather Bortfeld, Keanan J. Joyner, Silvia A. Bunge

**Author notes:** Corresponding author: Patrick G. Kelly (;). Joint senior authors (Keanan J. Joyner, Silvia A. Bunge).

## Abstract

Functional near-infrared spectroscopy (fNIRS) is a portable, motion-tolerant neuroimaging method particularly well suited for developmental and naturalistic research. To evaluate the utility of fNIRS for studying individual differences and longitudinal changes, we measured activation and functional connectivity during a relational reasoning task in young adults (N = 73). We sought to (1) establish whether fNIRS captures frontoparietal activation patterns consistent with prior fMRI studies using similar paradigms, (2) assess the effect of the amount of data (number of task blocks) on signal strength and precision, (3) assess the paradigm’s measurement properties in the form of intra- and interindividual stability of activation and functional connectivity within and across testing sessions, and (4) examine whether grouping channels into anatomical regions of interest (ROIs) conferred benefits to the above. We observed robust task-evoked activation across lateral prefrontal and parietal cortices, with effect sizes on par with prior fMRI studies. Generally, we observed diminishing returns in effect size and measurement precision beyond ∼7 minutes. Internal consistency and test-retest reliability varied across metrics; while they were very low for a specific task contrast, they were extremely high for functional connectivity, confirming the robustness of channel- and ROI-level connectivity as a stable marker of functional architecture. Exploratory analyses supported prior observations of lower signal quality in participants with darker skin tones and hair, underscoring the need for inclusive methodological strategies. Together, these findings highlight key design considerations for optimizing longitudinal and individual-differences research on higher-level cognition, particularly in diverse and developmentally variable populations.

**Highlights:**

- We measured within- and between-session reliability of fNIRS metrics
- Collecting more data yielded diminishing returns in effect size and precision
- There were tradeoffs to aggregating channel data into regions of interest
- General task activation was more reliable than a specific task contrast
- Functional connectivity showed extremely high test–retest reliability

## 1. Introduction

A central goal of developmental cognitive neuroscience is to understand the neural changes underpinning higher-level cognition, including reasoning. Prior work using functional magnetic resonance imaging (fMRI) has yielded insights into the neural basis of relational reasoning and its development from middle childhood onward (Dumontheil et al., 2010; Wendelken et al., 2011; 2017). However, to extend this research to younger and more diverse and representative populations and naturalistic settings, there are advantages to using alternative neuroimaging methods that are portable, versatile in terms of testing conditions, and tolerant of head motion, such as functional near-infrared spectroscopy (fNIRS).

fNIRS has gained significant attention in recent years due to its robustness to motion artifacts and environmental noise, making it particularly suitable for applications beyond traditional laboratory settings (Huppert et al., 2006; Pinti et al., 2018; Burgess et al., 2022) and for studying hard-to-test populations (Lloyd-Fox et al., 2010). Similar to fMRI, fNIRS indexes changes in blood oxygenation levels. The increased metabolic demands of highly active brain tissue results in elevated concentrations of oxygenated hemoglobin (HbO) in the blood, as well as reduced concentrations of deoxygenated hemoglobin (HbR), both of which can be measured by shining near-infrared light via a head cap at the wavelengths absorbed by HbO and HbR. fNIRS provides a cost-effective and versatile alternative to fMRI for studying functional brain activity (Ferrari & Quaresima, 2012). Although fNIRS has poorer spatial resolution than fMRI and shallower measurement depth (typically ∼3-4cm), it has a higher sampling rate, greater motion tolerance, and fewer practical testing restrictions. Accordingly, fNIRS has attracted growing attention over the last two decades, now with widespread use in developmental research – particularly in infants (Smith et al., 2017; Gu et al., 2017; Weibley et al., 2021; Fiske et al., 2022; Skau et al., 2022; Hancock et al., 2023; Amaireh et al., 2024).

To advance the use of fNIRS in developmental research, it is critical to move beyond demonstrations of feasibility and begin systematically evaluating its reliability and measurement properties across different contexts and timescales – i.e., its neurometric properties (analogous to psychometrics). With regards to fMRI data, test–retest reliability, typically measured via intraclass correlation coefficients (ICCs), is often low, but varies by metric and methodology (Noble et al., 2021; Elliott et al., 2020; Kragel et al., 2021). For example, one meta-analysis found that reliability was particularly low for task-evoked activation at the voxel and regional levels, and for resting-state connectivity at the edge level, but that it improved under certain conditions, such as shorter intervals between sessions and task-based paradigms for estimating functional connectivity (Noble et al., 2021). Relative to fMRI, however, there has been less methodological research evaluating the reliability and robustness of fNIRS, including its measurement properties (but see Holmes et al., 2024; Yücel et al., 2024). To determine its utility for developmental research, it is crucial to assess both test-retest reliability across sessions, important for tracking developmental or intervention-related changes, to assess its ability to reliably measure longitudinal changes over development or over the course of an intervention, in addition to its and internal consistency (within-sessions reliability), important for studying and understanding individual differences. Relatedly, since reliability can vary depending on measurement conditions and the paradigm employed, it is important to identify the optimal amount of data needed to achieve reliable results for a given paradigm. This complements the growing scrutiny of the variability in collection, preprocessing, and quantification of the fNIRS signals across researchers (Yucel et al., 2025)—all of which are needed to enable rigorous scientific investigations using this tool.

While fNIRS has reliably captured group-level effects during cognitive and motor tasks, its ability to produce stable individual-level activation patterns across sessions has not yet been widely examined. Recent studies have employed fNIRS to examine load-dependent activation in working memory tasks (Yeung et al., 2023; Nittel et al., 2025), motor cortex engagement during finger tapping (de Rond et al., 2023), and resting-state connectivity in clinical populations (Xu et al., 2023). Yeung et al. (2023) reported high test-retest reliability in behavioral performance but only moderate reliability in neural signals, with regional differences in signal stability. Similarly, Nittel et al. (2025) reported consistent group-level activation during inhibitory control and working memory tasks but observed substantial variability in individual-level neural reliability. In a motor task study, de Rond et al. (2023) found high short-term reliability when the fNIRS cap remained in place between sessions, but reliability declined significantly when the cap was removed and refitted 24 hours later. Xu et al. (2023) showed that resting-state fNIRS in stroke patients exhibited strong test-retest reliability for global and local efficiency metrics, particularly in low-frequency bands and with longer scan durations. Together, these findings underscore the utility of fNIRS for detecting group-level effects, while leaving open the question of how to achieve stable and reproducible individual-level measurements across sessions.

With respect to functional connectivity, even less is known. Although recent studies have used fNIRS to examine inter-regional coherence, network-level dynamics, and functional connectivity (Chenot et al., 2021; Arredonado et al., 2022; Zhilin et al., 2023; Zou et al., 2023), the stability of these metrics within and across sessions remains largely unexplored. These gaps in the literature highlight the need for further methodological advancements to enhance the reproducibility of fNIRS measures before they can be reliably applied in clinical or developmental contexts. Here, we aim to evaluate the robustness of both the within- and between-session reliability of task activation and functional connectivity, both at the level of individual channels and aggregated groups of channels in a region.

A key methodological consideration is how much data is sufficient to ensure robust and interpretable estimates of fNIRS activation and connectivity. While longer task durations can improve signal quality and statistical power by enhancing the signal-to-noise ratio, this comes with diminishing returns; indeed, prolonged data collection could compromise the ability to measure the cognitive processes of interest. Long sessions can result in attentional decline and participant fatigue, on one hand, or practice-related performance improvements on the other. Additionally, prolonged wear of the tight-fitting fNIRS cap can lead to physical discomfort, an aspect that is especially important for developmental populations. Thus, identifying the optimal data quantity needed to achieve stable, high-quality neural metrics is of great importance. This is particularly consequential for developmental and clinical populations, who are more susceptible to cognitive overload and less tolerant of lengthy procedures and long testing sessions. As such, we conducted a systematic evaluation of how activation strength, variability, and statistical significance change with increasing block count, aiming to inform fNIRS study design in sensitive populations, and in time-limited and sensitive research environments.

To address these questions, we collected fNIRS data using a paradigm previously employed in behavioral and fMRI research to investigate relational reasoning. Prior fMRI studies of relational reasoning paradigms consistently implicate two broad regions, lateral prefrontal cortex (LPFC) and posterior parietal cortex (PPC), as central hubs in the reasoning network (Christoff et al., 2001; 2003; Jung & Haier, 2007; Prado et al., 2011; Krawczyk, 2012; Bunge & Vendetti, 2014; Holyoak & Monti, 2021). Within these broader regions, rostrolateral prefrontal cortex (RLPFC) and inferior parietal lobule (IPL) have been linked to higher-level *relational thinking*—or the capacity to represent, compare, and integrate relationships among sets of stimuli—for example in the service of analogical reasoning (e.g., Christoff et al., 2001; 2003; Bunge et al., 2005; Green et al., 2006; Wendelken et al., 2011; Green et al., 2017). Moreover, reasoning performance has been linked to both lateral frontoparietal functional connectivity and white matter tract coherence (Shokri-Kojori et al., 2012; Cocchi et al., 2013; Ebisch et al., 2013; Bazargani et al., 2014; Wendelken et al., 2017).

While numerous fNIRS studies have studied the neural basis of executive functions (e.g., Cutini et al., 2008; Moriguchi & Hiraki 2013; Pinti et al., 2018), this is, to our knowledge, the first study to use fNIRS to examine relational reasoning. As such, we begin by evaluating whether this neuroimaging method can reliably detect activation and functional connectivity patterns consistent with prior findings from fMRI. To this end, we placed optodes over bilateral LPFC and PPC and measured fNIRS activation and functional connectivity on a variant of the relational matching task previously used in behavioral and fMRI studies involving both adults and/or children (Christoff et al., 2003; Dumontheil et al., 2010; Wendelken et al., 2011; Starr et al., 2022). This paradigm was designed to manipulate relational complexity (Halford et al., 1998) while minimizing other cognitive demands. In the task variant used here, participants assessed similarity among colored shapes, with two levels of relational complexity (see **Figure 2**).

1st-order trials involved direct comparisons of features (color or shape) between two paired images, whereas 2nd-order trials involved comparing two pairs of images, and determining whether they matched along the same feature (color or shape). Thus, successful completion of 2nd-order trials required integration of two relations. Prior fMRI research comparing blocks of trials of each type on relational matching tasks has shown stronger LPFC and PPC activation in response to 2nd-order than 1st-order blocks of trials, particularly in RLPFC and IPL (Christoff et al., 2003), and often more selectively in the left hemisphere (Wendelken & Helskog, 2009; Wendelken et al., 2011).

Our primary fNIRS analyses focused on HbO concentration, given its higher signal-to-noise ratio compared to HbR. We conducted analyses at both the single channel level (38 channels) and aggregated across multiple neighboring channels into 10 regions of interest (ROIs)—bilateral RLPFC, dorsolateral PFC (DLPFC),ventrolateral PFC (VLPFC), IPL, and superior parietal lobule (SPL)—to test whether average ROI metrics would be more stable than individual channels. Given its widespread use in fNIRS research, the ROI aggregation approach was employed alongside all channel-level analyses to evaluate the validity of anatomically defined ROI grouping and to assess the associated trade-offs in spatial specificity and statistical robustness.

A subset of participants completed two sessions spaced 2–4 weeks apart, allowing us to assess the between-session and within-session reliability of behavioral performance, task-evoked activation, and functional connectivity using GLM-based activation estimates and channel-level pearson and ICCs of HbO concentration. In addition to task-evoked activation, we examined functional connectivity, measured as correlations between fluctuating HbO over time, enabling us to assess whether network-level coordination of cortical regions during reasoning tasks is consistent over time.

In sum, we first sought to test whether 2nd-order relational reasoning blocks would elicit greater frontoparietal activation and functional connectivity than 1st-order blocks, consistent with increased relational integration demands. Based on prior fMRI findings, we expected the strongest activation for 2nd vs. 1st-order blocks in channels located around RLPFC and IPL, particularly in the left hemisphere. Next, we evaluated the within- and between-session reliability of task-evoked activation and functional connectivity at both the channel and ROI levels.These analyses also served to assess the validity of aggregating channels into anatomically based ROIs, helping to determine whether, or when, such aggregation may enhance signal stability or reduce spatial specificity.

We then examined how activation strength, variability, and statistical power changed as a function of the number of task blocks included in the analyses, so as to inform design considerations for time-constrained studies. Finally, to explore the inclusivity and generalizability of fNIRS for diverse populations, we conducted exploratory analyses building on recent work highlighting the effects of skin tone, hair color, and hair texture on fNIRS data quality (Holmes et al., 2024; Simmons et al., 2025; Yücel et al., 2024). The overarching goal of this study was to evaluate the validity and reliability of fNIRS to investigate relational reasoning, with the goal of enabling its broader application across a variety of populations and paradigms in future research.

## 2. Methods

### 2.1 Participants

#### 2.1.1 Full sample

A total of 84 young adults participated in this study; four participants were excluded due to poor task performance, and an additional seven participants were excluded due to low quality fNIRS data (as per criteria described below), resulting in a final sample of 73 participants (*M*_age_ = 21.03 years, SD = 2.09, range = 18-29) (see **Supplementary Table 1** for demographic information). Participants were students enrolled in a Psychology course who were offered course credit for their participation. The total duration of the study was between 60 and 90 minutes. Participants gave written informed consent before participation, and all experimental procedures were approved by the Institutional Review Board at the University of California, Berkeley. All participants had normal (or corrected to normal) vision, no pre-existing neurological or sensorimotor deficits, and no history of self-reported major psychiatric illnesses.

#### 2.1.2 Functional Connectivity Sample

For our initial functional connectivity analysis, we selected a subset of 36 participants with the highest quality data (no more than 8 bad channels out of 46) for pairwise channel-level (channel x channel) connectivity analyses (see **Supplementary Table 1** for demographic information). We additionally performed ROI-level connectivity analyses on both this subsample and the full final sample of 73 participants.

#### 2.1.3 Test-Retest sample

To evaluate the stability of task activation and functional connectivity, we recruited a subset of 26 participants who had high-quality data during session 1 and were willing to return 2-4 weeks (mean = 20.31 days) after their first session (see **Supplementary Table 1** for demographic information). All of these participants met our standard inclusion criteria, and were therefore included in the main analyses. From this test-retest subset of 26 participants, we additionally conducted follow-up functional connectivity analyses on 17 of these participants who had no more than 8 bad channels. ROI-level results for the full sample, reported in the main text, were almost identical to those obtained for the 17 participants with the highest quality data.

### 2.2 fNIRS Data Acquisition

fNIRS data were collected from a continuous-wave NIRSport2 device (NIRx Medical Technologies, LLC). The wavelengths of emitted light (LED sources) in this system were 760 nm and 850 nm. The data were collected at a sampling rate of 5.1 Hz using the NIRx acquisition software, Aurora fNIRS. The fNIRS cap contained a total of 16 sources and 15 detectors, with the 16th detector used for 8 short channel detectors, creating 38 total channels covering bilateral LPFC (22 channels) and PPC (16 channels). The optode montage (**Figure 2**) was designed using fOLD (fNIRS Optodes’ Location Decider; Morais et al., 2018) in combination with an MNI-to-EEG cap converter to optimize coverage of lateral prefrontal and parietal cortices. Guided by prior relational reasoning studies (Wendelken et al., 2008, 2012; Wendelken & Bunge, 2010; Watson & Chatterjee, 2012; Vendetti & Bunge, 2014), target locations were mapped to the international 10–20 system using the Brodmann Brain Atlas. Ten sources and seven detectors were positioned to cover bilateral dorsolateral (BA 9/46), rostrolateral (lateral BA 10), and ventrolateral (BA 45/47) PFC, while six sources and eight detectors targeted bilateral inferior (BA 39) and superior (BA 7) parietal lobules. Analyses were conducted at both the channel and ROI levels, with the 36/38 channels that met our criterion for signal quality being grouped into these 10 anatomically defined regions (see **Figure 1C** and **Supplementary Table 2**).

**Figure 1.**
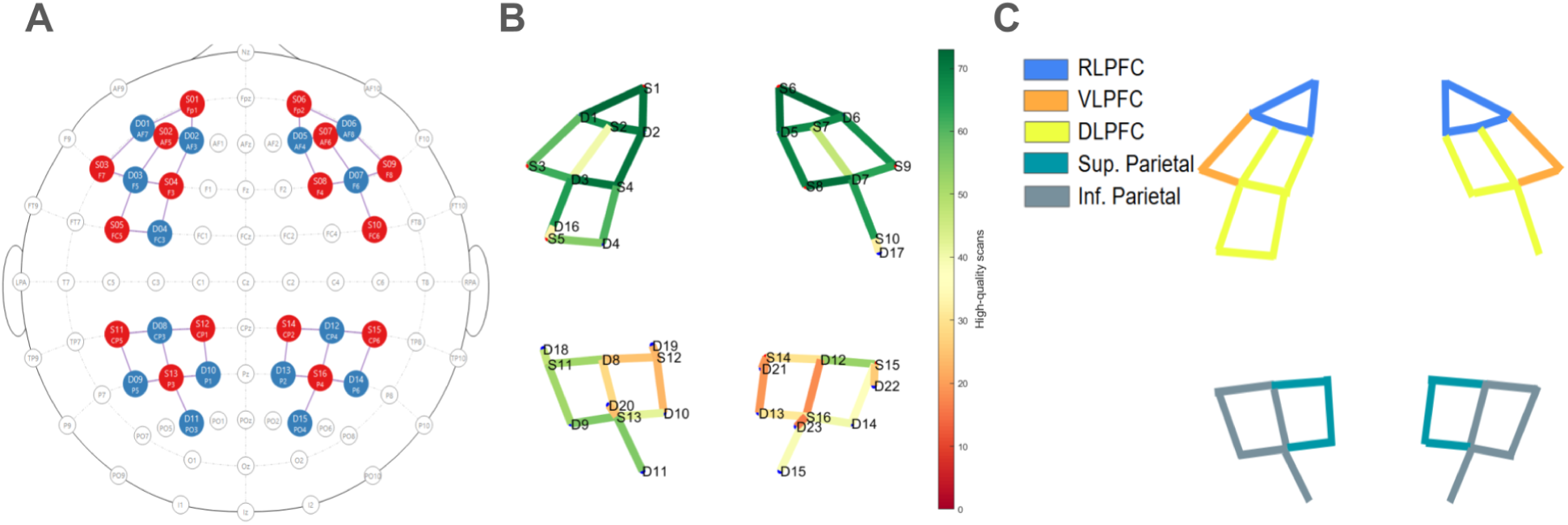
**A**: fNIRS Montage created using the NIRSite software showing the source and detectors in their respective locations that cover bilateral frontal and parietal cortex **B**. Montage figure generated by the QT NIRS package in MATLAB showing the amount of good quality data for each channel in its location within the montage indicating a higher proportion of good quality data in the frontal channels than parietal channels **C.** Colored coded montage diagram showing how the channels that were aggregated into anatomical ROIs.

### 2.3 Experimental Procedures and Task Design

Participants were prescreened online at the time they signed up for the study. Upon arrival, they completed an additional prescreening survey to ensure they had no neurological, sensory motor, or any self-reported major psychiatric disorders that would exclude them from participating in the study. They then provided informed consent and completed a demographic survey. Once all the intake and pre-screening procedures were completed, experimenters measured the participants’ head to determine cap size and placement, then began to set up the cap while participants were given task instructions. The cap was then placed on the participant’s head, moving hair as needed to provide clear access to the scalp for the sources and detectors. Cap alignment was verified based on the international 10–20 location of Cz (Klem et al., 1999). fNIRS data were then calibrated and checked for quality using the Aurora fNIRS Software before proceeding with the experiment. If any channels did not have sufficiently high quality signal (see **Section 2.4.2** for criteria), placement and hair-clearing were performed again. The task was coded and presented using the experiment software PsychoPy (Peirce et al., 2019). Visual stimuli consisted of a stimulus array of four simple colored geometric figures. Two figures were displayed in the top row and two in the bottom row, thereby facilitating the comparison of relations among objects within a row and across rows. Stimulus arrays were presented pseudorandomly and included eight distinct shapes presented in one of eight colors. These figures were combined in unique pairings on each trial to eliminate the possibility of familiarity effects.

The relational matching task consisted of three trial types: Color, Shape, and Match. Color and Shape blocks involved 1st-order judgments, and Match blocks involved 2nd-order judgments (**Figure 2**). On each trial, participants saw two pairs of objects, with a line separating the top pair from the bottom pair. In 1st-order trials, the objects varied either in color or shape. A cue word (“Color” or “Shape”) simultaneously indicated whether participants should focus on the top or bottom row and the type of judgment they should be making. On Shape and Color trials, respectively, participants determined whether the two objects in the cued row had the same shape or color. For example, if the word ‘Shape’ appeared next to the top row of objects, that indicated to the participant to make a shape judgment on the top set of objects. On 2nd-order trials, participants determined whether the relationship between the objects in the top pair was the same as the relationship between the bottom pair of objects. Here, the cue word “Match” was presented beside the line separating the two rows. Across the entire experiment, there were equal numbers of valid and invalid trials for each of the three trial types

**Figure 2.**
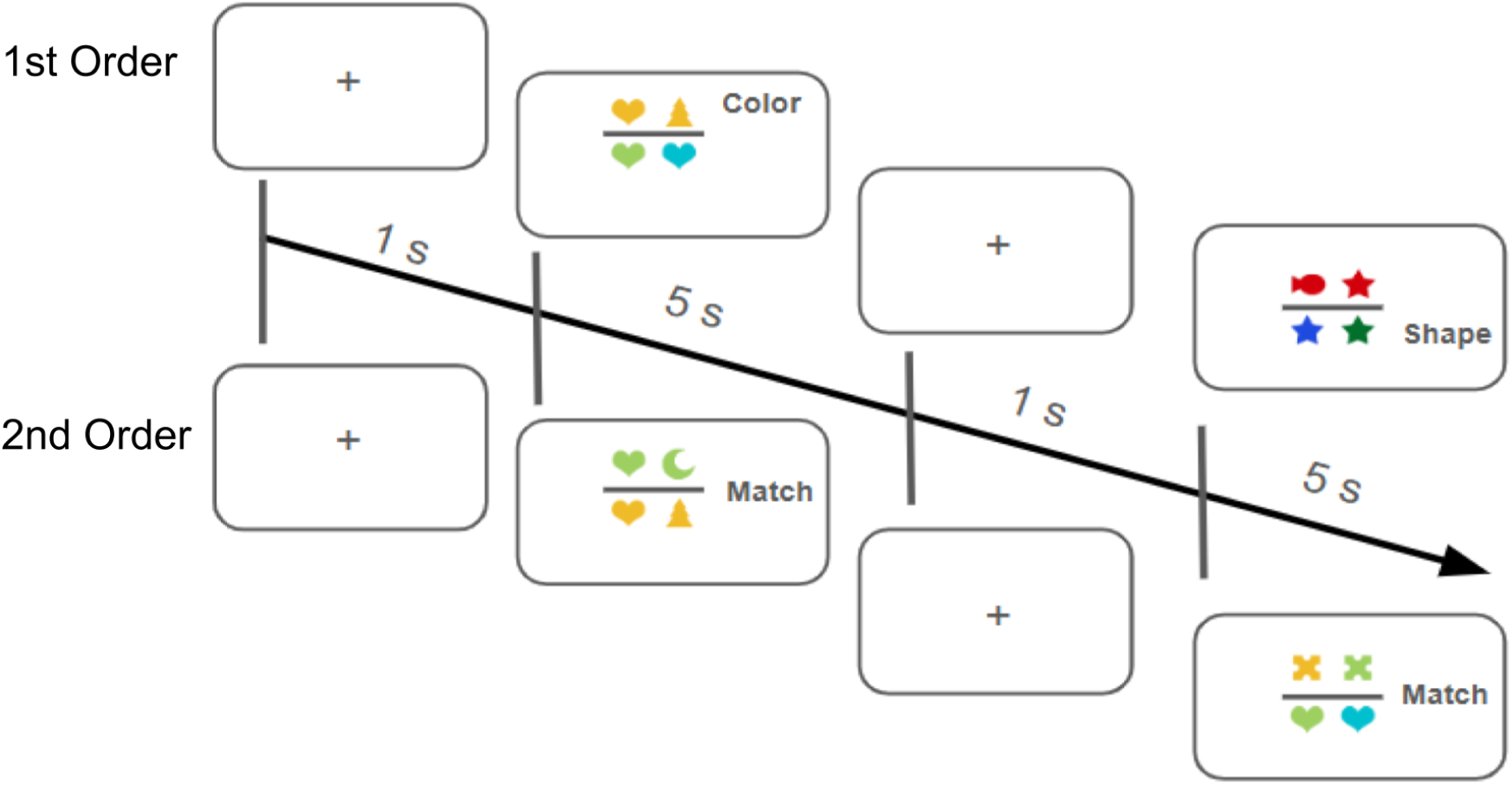
Participants completed a relational matching task consisting of 1st-order (Color and Shape) and 2nd-order (Match) blocks. Each trial displayed two object pairs, with a cue indicating whether to judge shape, color, or relational similarity. Shape and Color blocks required simple feature matching (e.g., “Are these the same shape?”), while Match blocks required comparing the relationship between the top and bottom pairs. Trials lasted 5 seconds with a 1-second inter-trial interval. Blocks consisted of six trials (36 seconds total), presented in a pseudorandomized order and interleaved with jittered rest periods (16–20 seconds). The full session included 9 1st-order blocks, 9 2nd-order blocks, and 19 rest periods (∼16.7 minutes total).

The task was presented in a blocked design, with task blocks including 6 1st-order trials (with Shape and Color trials intermixed) or 6 2nd-order trials. Individual trials were 5 seconds long (stimulus remained on the screen for the entire duration regardless of response time (RT)), with a 1-second inter-trial interval. Individual task blocks were 36 seconds long, and were interleaved with jittered rest periods of 16-20 seconds (16, 18, or 20 seconds, with equal numbers of rest periods of each duration presented pseudorandomly across the experiment). Task blocks were presented in a random order for each participant, and were always followed by a rest period. The entire experiment consisted of a total of 9 1st-order blocks, 9 2nd-order blocks, and 19 rest periods, for a total duration of ∼1000 seconds (16.7 minutes). There was a break half-way through, and participants indicated via a button press when they were ready to move on.

Prior to the start of the experiment, participants were given instructions, followed by five untimed practice trials with verbal feedback for each block type. Prior to data acquisition, they performed five timed trials for each block type without feedback, to familiarize them with the pacing of the task. During these practice trials, researchers monitored their understanding and adherence to task instructions.

## 3. Analyses

All statistical analyses were performed using the pandas and scipy libraries in Python (v. 3.11; Van Rossum & Drake, 2009). Four types of statistical analyses were used to determine the relationship between variables: Pearson’s correlations, ICCs, robust regressions, and permutation-based significance testing. The significance level was set to *p* < 0.05, and post-hoc *p*-values were corrected for multiple comparisons using the Holm-Bonferroni procedure (Abdi, 2010). Results are reported based on group-level statistical parametric maps and false discovery rate (FDR)-corrected q-values (q < 0.05).

### 3.1 Behavioral performance

Accuracy and response times (RTs) were quantified by calculating task accuracy for each participant, defined as the proportion of correct responses on each condition across all 18 blocks of trials. Participants whose accuracy was below 70% on one or both conditions (3 participants on 1st-order blocks and one additional participant on 2nd-order blocks) were excluded from further analysis, leaving a subsample of 80 participants whose data were subsequently screened for fNIRS data quality.

### 3.2 fNIRS quality check and pre-processing pipeline

Because fNIRS signals are sensitive to superficial physiological noise (Santosa et al., 2018), we identified and excluded individual channels and participants exhibiting low-quality data. Following a visual inspection of each participant’s data in the NIRS Toolbox, we conducted a channel-wise quality assessment on the raw fNIRS data using the QT-NIRS module (Montero-Hernandez & Pollonini, 2019), which integrates multiple metrics to identify low-quality measurements. QT-NIRS implements a data-driven algorithm to evaluate signal quality based on physiological and signal characteristics. The criteria outlined below were applied for each channel and participant. Any channels that failed to meet these criteria were flagged as bad channels and removed from subsequent analyses.

Several thresholds were configured: we set the qThreshold to 0.7, specifying that a channel must pass at least 70% of quality checks to be retained; the standard fCut parameter was set to [0.5-2.5] Hz, specifying the frequency range used to estimate the presence of physiological oscillations (e.g., heartbeat, respiration). This frequency content is extracted using a bandpass filter, and its power is analyzed to assess the presence of expected physiological signals, indicating functional optode-skin coupling. We selected a sciThreshold of 0.7, using the Scalp Coupling Index—a correlation-based metric that compares short- and long-separation channel signal shapes to assess optode contact quality. Channels with SCI values below this threshold were classified as poorly coupled. Additionally, the pspThreshold evaluation is based on Peak Spectral Power (PSP). PSP calculates the ratio of spectral power in the expected physiological band (typically around the cardiac frequency) to the total signal power. A low PSP indicates a lack of physiological oscillations in the signal, suggesting poor data quality due to insufficient physiological engagement or sensor detachment. By setting a threshold 0.1, channels with a peak spectral power below this level are flagged as low-quality.

We also calculated the number of participants with high-quality data per channel to assess overall signal quality and excluded channels that had insufficient high-quality data across participants from subsequent analyses. An *a priori* power analysis conducted using G*Power indicated that a minimum of 23 high-quality data points per channel was required to achieve 75% power (1 − β = 0.75) to detect a moderate effect size (Cohen’s d = 0.5) at an alpha level of 0.05 using a one-tailed test. Based on these criteria, four of the 46 channels, 2 in right SPL and IPL (S14_D13, S16_D12), and 2 short separation channels (S14_D21, S16_D23) were excluded from subsequent analyses (see **Supplementary Figure 1**). Additionally, one channel (S12_D10) and two short channels (S12_19, S13_D20) located over left SPL consistently produced signal quality values that were near or at the exclusion threshold across multiple participants, highlighting a broader issue in fNIRS research: superior parietal cortex is notoriously difficult to image reliably due to increased skull thickness (Beauchamp et al., 2011; Tachtsidis et al., 2016; Skau et al., 2022; Klein, J.H., 2024).

To identify and exclude participants with poor fNIRS data quality, we calculated the mean and standard deviation of the number of bad channels across all scans (mean = 16.12, SD = 8.63). Participants for whom the number of bad channels exceeded 1 SD above the mean (i.e., >27 bad channels) were excluded from further analysis (see **Supplementary Figure 2**). This approach downweights noisy channels in the linear model (see next section). Including these channels does not increase the likelihood of a false positive effect, but the statistical power to detect an effect in this area is reduced (Santosa et al., 2018). Based on this exclusion criterion, seven of the 80 participants with good behavioral data were identified and excluded from further analyses resulting in a final sample of 73 participants.

After excluding low-quality channels, fNIRS data were analyzed using the NIRS Toolbox in MATLAB. Raw .nirs files containing light intensity measurements were imported and converted to optical density. We applied motion correction using both temporal derivative distribution repair (TDDR) to remove any motion artifacts from the optical density data (OD), and resampled the data to 1.5 Hz. We then converted the cleaned OD data into oxygenated (HbO) and deoxygenated (HbR) hemoglobin concentrations using the modified Beer-Lambert law (Strangman et al., 2003). After each participant’s data were cleaned and converted to HbO and HbR concentrations, participant-level statistics were calculated. All subsequent analyses focused on HbO, given its higher signal-to-noise ratio compared to HbR (Pinti et al., 2019). Results for HbR are provided in **Supplementary Figure 4.**

#### 2.4.4 fNIRS Signal Processing

fNIRS data have distinct properties that are not adequately addressed by conventional fMRI-based analysis methods, which can lead to inflated type I error rates (Huppert, 2016). Compared to fMRI, fNIRS data are more strongly affected by serially correlated errors due to its higher sampling rate relative to the underlying physiological signals, as well as by heavy-tailed noise distributions resulting from motion artifacts and substantial variability in signal-to-noise ratio across channels and participants (Huppert, 2016). To address these challenges, a first-level general linear model was applied to each participant’s data using an autoregressive, iteratively reweighted least-squares (AR-IRLS) approach. The AR-IRLS model utilizes an autoregressive filter known as ‘pre-whitening’ to correct for serially correlated errors, along with robust weighted regression to iteratively down-weight motion-related outliers and physiological noise (Barker et al., 2013). This pre-whitening filter is applied to both the participant-level design matrix and the resulting data to remove correlated noise inherent in continuous fNIRS signals. This model generates participant-level regression coefficients along with their error-covariance matrices, which are used for individual statistical tests, contrast estimation, and subsequent group-level analyses.

In line with current research on the sensitivity and specificity of fNIRS basis sets depending on signal quality and task duration (Santosa et al., 2019), a canonical haemodynamic response function (HRF) basis set was chosen for this analysis. Santosa et al. (2019) found that for task blocks lasting more than 10 seconds (as in this study), the canonical HRF offers optimal performance in a sensitivity-specificity (ROC) analysis. Compared to a full deconvolution of the hemodynamic response (finite impulse response functions, or FIR model), the canonical model has greater degrees of freedom, making it better able to detect a true signal in the presence of noise while improving both sensitivity and specificity in the analysis (Santosa et al., 2019).

Next, group-level statistical models were calculated, using the full covariance matrix from the first-level models to perform a weighted least-squares regression (Santosa et al., 2018) with condition as a fixed effect and subject as a random intercept (beta ∼ -1 + cond + (1|subject)). Robust regression was also applied to the group-level model to down-weight outliers at the group-level. The results of this analysis were used for the following group-level contrasts for each individual channel: 1st-order activation compared to baseline, 2nd-order activation compared to baseline, the 2nd-order vs. 1st-order contrast, and the Task activation vs. Baseline contrast (1st- and 2nd-order combined into a single ‘task’ condition). Group activation results are reported as statistical maps using channel-level beta values (mean task-related change in hemoglobin concentration) and *t* values (beta divided by SE) with corresponding *p* values.

A slightly different approach was taken to conduct our anatomically informed, ROI-based, analysis. Channel-wise HbO signals were aggregated into ten bilateral cortical regions of interest (ROIs) based on Brodmann Areas (see **Table 1** and **Figure 1C**), spanning RLPFC, DLPFC, and VLPFC, as well as superior and inferior parietal areas. For each ROI, signals from predefined source-detector pairs were averaged to produce a single representative time course. Dummy source-detector mappings were then assigned to ensure compatibility with group-level visualization routines. General Linear Model (GLM) analysis was conducted at the subject level using autoregressive iteratively reweighted least squares (AR-IRLS), which corrects for serial correlations and outliers in the time series as described above. Group-level inference was performed using a linear mixed-effects model with condition as a fixed effect and participant as a random intercept (beta ∼ -1 + cond + (1|subject)), and dummy coding enabled for condition comparisons. A planned contrast comparing condition 2 to condition 1 was computed using a contrast vector [-1, 1, 0], and corresponding statistical maps were visualized and exported. All participant- and group-level results were exported for further statistical analysis.

Benjamini-Hochberg false-discovery rate-corrected *p*-values (e.g., *q*-values; Benjamini and Hochberg, 1995) were applied to all group-level analyses, including 36 channels (10 for ROI aggregation analysis), both oxy- and deoxy-hemoglobin signals, and three experimental conditions, making the correction highly conservative across all tests.

### 3.3 Effects of Number of Task Blocks on Task Activation

To determine the amount of fNIRS data required to obtain stable and interpretable neural activation estimates, we examined t-values and channel-level variance based on incremental block inclusion at both the group and subject levels using activation estimates from the 2nd vs. 1st-order Contrast and the Task vs. Baseline contrast. Specifically, we reanalyzed the data in intervals of 3 blocks (i.e., using the first 3, 6, 9, 12, 15, and all 18 blocks) to evaluate how the quantity of task data influenced the reliability and strength of activation patterns. For each increment of block count, group-level general linear models (GLMs) were computed and beta and t-values as well as standard error (SE) were extracted to assess how activation evolved as a function of increasing amounts of data. For each block count, we calculated summary statistics including the mean of beta values and SE across channels, as well as the proportion of significant channels. Beta values and SE were first averaged across channels within each participant before being aggregated across participants for each block count.

This two-tiered fNIRS analysis allowed us to quantify data stability as a function of the number of blocks. Importantly, this is not necessarily a case of “more is better”: while increasing the amount of data can enhance statistical power and reduce noise, it may also introduce trade-offs, such as participant fatigue (e.g., attentional drift) and practice-related effects (e.g., increased automaticity or shifts in task strategy). To interpret fNIRS effects across the session in the context of behavioral performance, we also measured accuracy and RT for each of the 18 task blocks. This approach can guide the design of future studies involving young children or clinical populations with limited tolerance for extended scanning, as well as tasks prone to pronounced practice effects, by empirically determining the number of blocks needed to obtain reliable estimates of task-evoked neural activity.

### 3.4 Functional Connectivity Analyses

Functional connectivity analyses were also performed using the NIRS Toolbox in MATLAB (Santosa et. al, 2018). First, the preprocessed time series for HbO concentration (see above) across the entire ∼1000 s collected for each participant was segmented by condition (1st-order and 2nd-order), resulting in a total of 1,500 samples (1.5 Hz × 1000 s) for each participant and condition. Functional connectivity was calculated on HbO only, given its higher signal-to-noise ratio than HbR (Pinti et al., 2019). Functional connectivity was computed using robust pairwise correlations between all channels within each condition block, employing an autoregressive (AR) innovations approach to remove autocorrelation artifacts and isolate inter-regional synchrony. Connectivity estimates were computed separately for each condition (1st-order, 2nd-order, and Task vs. Baseline), with all data from each block included in the connectivity estimates.

To evaluate condition-level differences in connectivity at the group level, a linear mixed-effects model was fit to the subject-level correlation coefficients (R) using the formula R ∼ -1 + cond + (1|subject), treating condition as a fixed effect and subject as a random intercept. This model enabled estimation of condition-specific connectivity patterns while accounting for within-subject dependence. Results were exported in .csv format for further statistical and visualization analysis in Python. By computing Pearson’s correlations between all channels, we obtained a coefficient for each pair – i.e., a pairwise correlation matrix of *r* values – for each participant for each condition. We then computed the average *r* coefficient of each element of the matrix across participants, resulting in a global matrix, showing the channel-level connectivity at the group level for channel level (resulting in a 36 × 36 correlation matrix) and the ROI level (resulting in a 10 × 10 correlation matrix.) For ROI analyses, we were able to include the full sample (N = 73), excluding channels marked as bad for each participant and averaged over the remaining valid channels within each ROI. For channel-level analyses, we included only a subsample of 36 participants with the highest signal quality, ensuring consistent coverage across source-detector pairs.

For ROI-level analyses, channel-wise time series were aggregated into Brodmann Area (BA)-based regions (see **Figure 1C** and **Table 1**) following the same preprocessing pipeline as the Task vs. Baseline analysis using the NIRS Toolbox in MATLAB. Functional connectivity matrices were estimated using pairwise robust correlations with an autoregressive innovation model (whitening = 4 × sampling rate), and group-level effects were modeled using a linear mixed-effects model with condition as a fixed effect and subject as a random intercept (*r* ∼ -1 + cond + (1|subject)).

To ensure data quality in the full sample (N = 73), QT-NIRS–based quality control procedures were applied to exclude low-quality channels on a participant-by-participant basis. Crucially, ROI aggregation enabled full anatomical coverage even in the presence of channel-level signal loss, as each participant contributed usable data to every ROI.

Subsequent functional connectivity analyses were performed using the pandas and scipy libraries in Python (v. 3.11; Van Rossum & Drake, 2009). Three types of statistical analyses were used to determine the relationship between variables: Pearson’s correlations, robust regressions, and permutation significance testing. The significance level was set to *p* < 0.05, and post-hoc *p*-values were corrected for multiple comparisons using the Holm-Bonferroni procedure (Abdi, 2010).

### 3.5 Within-Session Stability Analyses

To evaluate within-session spatial stability (individual-level) and internal consistency reliability (group-level) of task-evoked activation and functional connectivity, we separated stimulus blocks into interleaved (odd- and even-numbered) blocks and applied the full activation pipeline independently to each subset. For each half, we estimated condition specific task activation using participant-level general linear models (GLMs) and session-level linear mixed effects models (LMMs). Analyses centered on changes in oxyhemoglobin (HbO) signals, due to HbO’s greater sensitivity to task-evoked neural dynamics.

#### 3.5.1 Within Session Task Activation

To calculate internal consistency at the individual level, beta values for the Task vs. Baseline and 2-order vs. 1st-order contrasts were extracted for each channel for each participant. ICCs for absolute agreement (ICC(2,1)) were used to quantify internal consistency reliability. Cross-classified mixed effect models where halves of the data are nested 1) within-subjects, 2) within-channels, and 3) the interaction between subject and channel (beta ∼ 1 = (1|subject) + (1|channel) + (1|subject:channel)) to provide an estimate of internal consistency reliability of 1) the fNIRS signal on average at the individual-level, 2) stable channel-level variation in activation, and 3) variation across channels in the degree of reliability at the individual-level. This variation in reliability across channels was visualized by calculating the ICC value within each channel separately, and plotted in **Figure 9**. Again accounting for the reduced amount of data in each split-half, Spearman-Brown prophecy correction was applied to these ICC estimates.

For task activation at the group level, t-values for the Task vs. Baseline and 2nd-order vs. 1st-order contrasts were computed for each channel and ROI then extracted into vectors at the group level. At the group level, Pearson correlations were computed between odd and even halves to quantify the spatial consistency of activation patterns (roughly equal numbers of 1st-and 2nd-order blocks in each half). To account for the reduced amount of data in each split half, we applied the Spearman-Brown prophecy correction formula to the resulting reliability coefficients, yielding adjusted estimates of internal consistency for the full-length data (Spearman, 1910; Brown, 1910). In addition to raw t-value correlations, we assessed the stability of variance estimates (t²), correlating these across split halves.

#### 3.5.2 Within Session Functional Connectivity

For functional connectivity analyses, connectivity values (Pearson’s r) for both contrasts were extracted and matched between odd and even block-derived matrices for each channel or ROI pair. To assess the stability of group-level connectivity patterns, a Pearson correlation was calculated between the vectorized connectivity values from odd and even blocks, yielding a condition-specific reliability coefficient. At the individual level, we assessed the internal consistency reliability of individual functional connectivity profiles by correlating connectivity matrices derived from odd and even blocks for each subject.

This within-session correlation approach for interleaved blocks offers a robust index of the internal consistency of activation and functional connectivity estimates, capturing the reliability of network dynamics within a single session. In addition to this interleaved split-half analysis, we also examined change in fNIRS signal from the 1st half to the 2nd half of the first session to evaluate differences in activation across a testing session (see **Supplementary Section 9**).

### 3.6 Between-Session Stability Analysis

We assessed the stability of behavioral and fNIRS data across Sessions 1 and 2 in the 26 participants who were tested twice, with a 2-4 week interval between sessions. We examined the stability of accuracy and response times as well as fNIRS-derived task activation, functional connectivity, and variance at both the group and participant levels.

#### 3.6.1 Between Session Task Activation

To calculate test-retest reliability at the individual level, ICCs for absolute agreement were used that mirrored the split-half calculations, this time with the full data for each session as the two data points per subject. Cross-classified mixed effect models were specified where halves of the data were nested 1) within-subjects, 2) within-channels, and 3) the interaction between subject and channel (beta ∼ 1 + (1|subject) + (1|channel) + (1|subject:channel)), and because data are not being split in half, no Spearman-Brown prophecy correction is applied to estimate test-retest reliability. Variation in test-retest reliability across channels was visualized by calculating the ICC value within each channel separately, and plotted in **Figure 11**.

To assess test–retest reliability at the group level, task-related activation was estimated using first-level general linear models (GLMs) for each participant and session, followed by second-level linear mixed effects models (LMMs) for each session. Analyses focused on oxyhemoglobin (HbO) signals, with conditions modeled separately to capture task-specific activation. For each model, t-values were computed per channel and condition, aggregated into group-level vectors, and correlated across sessions to assess the spatial consistency of activation for each condition (1st-order, 2nd-order, 2nd > 1st-order, and Task vs. Baseline). To evaluate variance stability, t-values were squared (t²) to index signal strength irrespective of direction and correlated across sessions for each channel. This approach quantified the reliability of spatial activation patterns and the reproducibility of signal magnitude across sessions at both the group and individual levels.

#### 3.6.2 Between Session Functional Connectivity

We also evaluated the test–retest reliability of group-level functional connectivity by quantifying the consistency of Channel x Channel and ROI x ROI connectivity estimates across experimental sessions. Functional connectivity matrices derived from session 1 and session 2 were first restricted to channel pairs involving oxygenated hemoglobin (HbO), given its greater sensitivity to task-evoked neural responses. For each task condition represented in both sessions, channel-level connectivity values (Pearson’s r) were extracted and paired with identical channel and ROI pairs. To assess stability, a Pearson correlation was computed between the connectivity estimates from session 1 and session 2 for each condition, yielding a condition-specific reliability coefficient. Statistically, this procedure quantifies the linear relationship between corresponding channels and ROI-to-ROI edges across time points, providing a global metric of reproducibility in functional network structure. Higher correlation values indicate greater consistency of task-evoked connectivity at the group level, while lower values suggest increased session-to-session variability in network recruitment.

At the individual level, this analysis assessed the within-subject reliability of task-related functional connectivity patterns across sessions using a pairwise correlation approach. Channel x Channel and ROI x ROI functional connectivity matrices were derived independently for session 1 and session 2 and filtered to include only oxygenated hemoglobin (HbO) channel pairs, which are more sensitive to task-evoked hemodynamic changes. For each participant, connectivity values from corresponding ROI pairs (defined by source-detector combinations) were aligned across sessions. A Pearson correlation coefficient (r) was then computed between the connectivity strengths observed in session 1 and session 2, providing a metric for evaluating the stability of task-induced brain network organization at the individual level.

## 4. Results

### 4.1 Behavioral Results

To confirm participant engagement and assess task performance across conditions, we calculated accuracy and response times (RTs) for 1st- and 2nd-order trials. Four participants failed to meet the pre-established performance threshold of 70% accuracy for each condition, and were therefore not included in the final analysis. Participants in the final sample (N = 73) performed with accuracy above 90% in both conditions (Mean = 91.2, SD = .023 and Mean = 94.27, SD = .031 for 1st- and 2nd-order trials, respectively. Unexpectedly, accuracy was 3 percentage points higher for 2nd-order trials (*t*(72) = −7.77, *p* < .0001); see **Supplementary Figure 3**). Conversely, participants responded slightly but significantly more quickly on 1st-order trials (M = 1.56 s, SD = 0.31) than 2nd-order trials (M = 2.15 s, SD = 0.50), *t*(72) = −15.55, *p* < .0001, as predicted, suggestive of a small speed-accuracy tradeoff.

The unexpected finding of slightly higher accuracy in the 2nd-order condition, which had higher relational complexity, was small but significant. We hypothesize that it is attributable to higher task-switching demands on the 1st-order trials. Unlike previous studies using similar paradigms, the two 1st-order trial types – shape and color matches – were intermixed within blocks, as opposed to being presented in separate blocks. Moreover, in the current study design, participants were cued, in an unpredictable manner, to respond to one of two rows on every 1st-order trial. In sum, we posit that the increased relational complexity on 2nd-order trials was fairly well balanced by other executive function demands on 1st-order trials, rendering the 2nd-vs. 1st-order comparison a cleaner – albeit more subtle – task manipulation with which to identify activation associated with relational thinking than in prior studies.

### 4.2 Task-Based Activation

Results are reported based on group-level statistical parametric maps, with significant channel-level and ROI-level results defined as those having false discovery rate (FDR)-corrected q-values < .05).

#### 4.2.1 General Task Activation

To identify cortical regions engaged across both conditions relative to baseline, we performed a contrast comparing neural activation across all 18 blocks (1st and 2nd-order combined). This analysis revealed 20 channels across bilateral prefrontal and parietal cortices that exhibited significantly greater activation for task-related activity (**Figure 3** and **Table 2**). Robust activation was observed in the left and right RLPFC and DLPFC channels, with additional activation in a left VLPFC channel, reflecting distributed prefrontal engagement. Significant parietal activation was identified in left SPL and IPL channels, indicating broad posterior involvement. At the ROI level, significant activation increases relative to baseline were observed across nearly all prefrontal and parietal regions (**Table 2**) reflecting large-scale cortical recruitment aligned with task demands. While the channel-level analysis provided fine-grained spatial resolution, the ROI-level approach produced stronger and more widespread effects for this general contrast (see **Table 2**).

**Figure 3.**
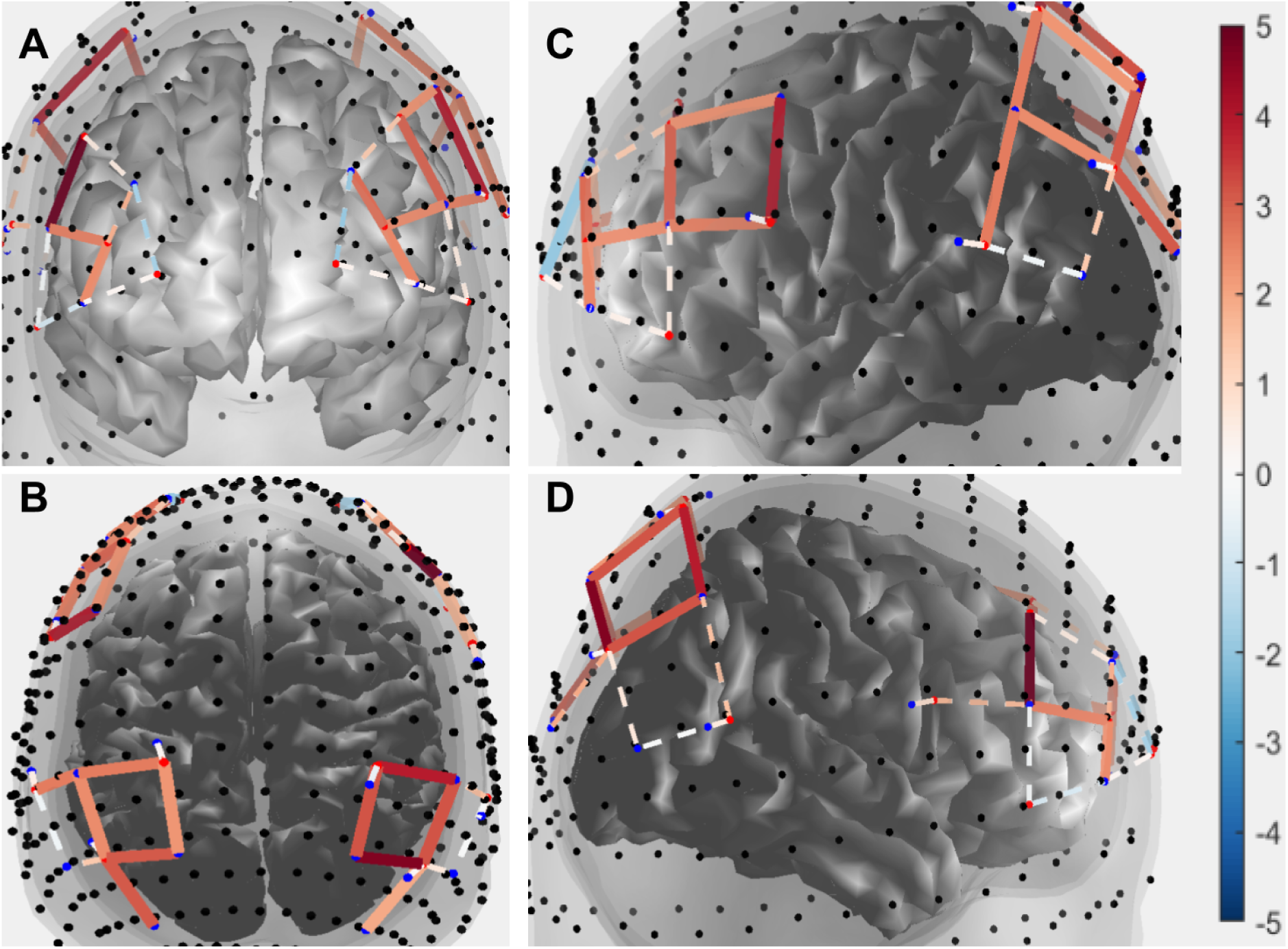
Channel level Task vs. Baseline contrast for task activation across participants. Solids lines indicate channels that were significant after correcting for multiple comparisons across 36 channels; and dashed lines indicate those that did not survive correction. The color bar indicates t-statistic ranging from 5 to -5 **A.** Anterior view of brain showing significant activation in channels over bilateral RLPFC, bilateral DLPFC **B**. Dorsal view of brain showing significant activation in channels over bilateral inferior and superior parietal cortex **C.** Lateral view of the left hemisphere showing significant activation in channels over left RLPFC and DLPFC, as well as left IPL and SPL. **D**. Lateral view of the right hemisphere showing significant activation in channels over right RLPFC, right DLPFC, and right IPL and SPL.

**Table 1.**
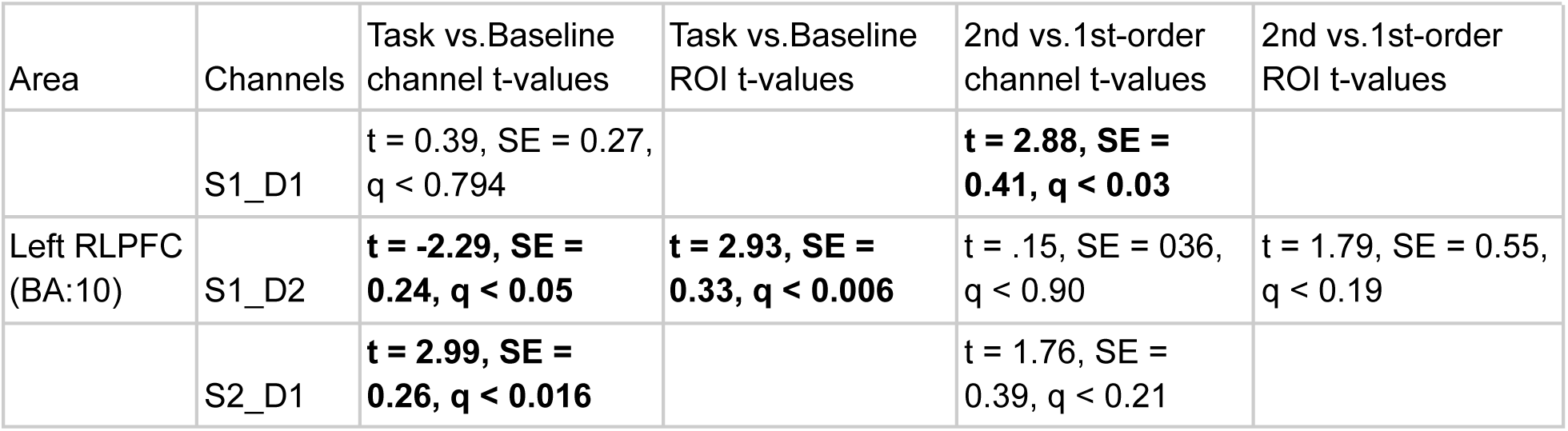
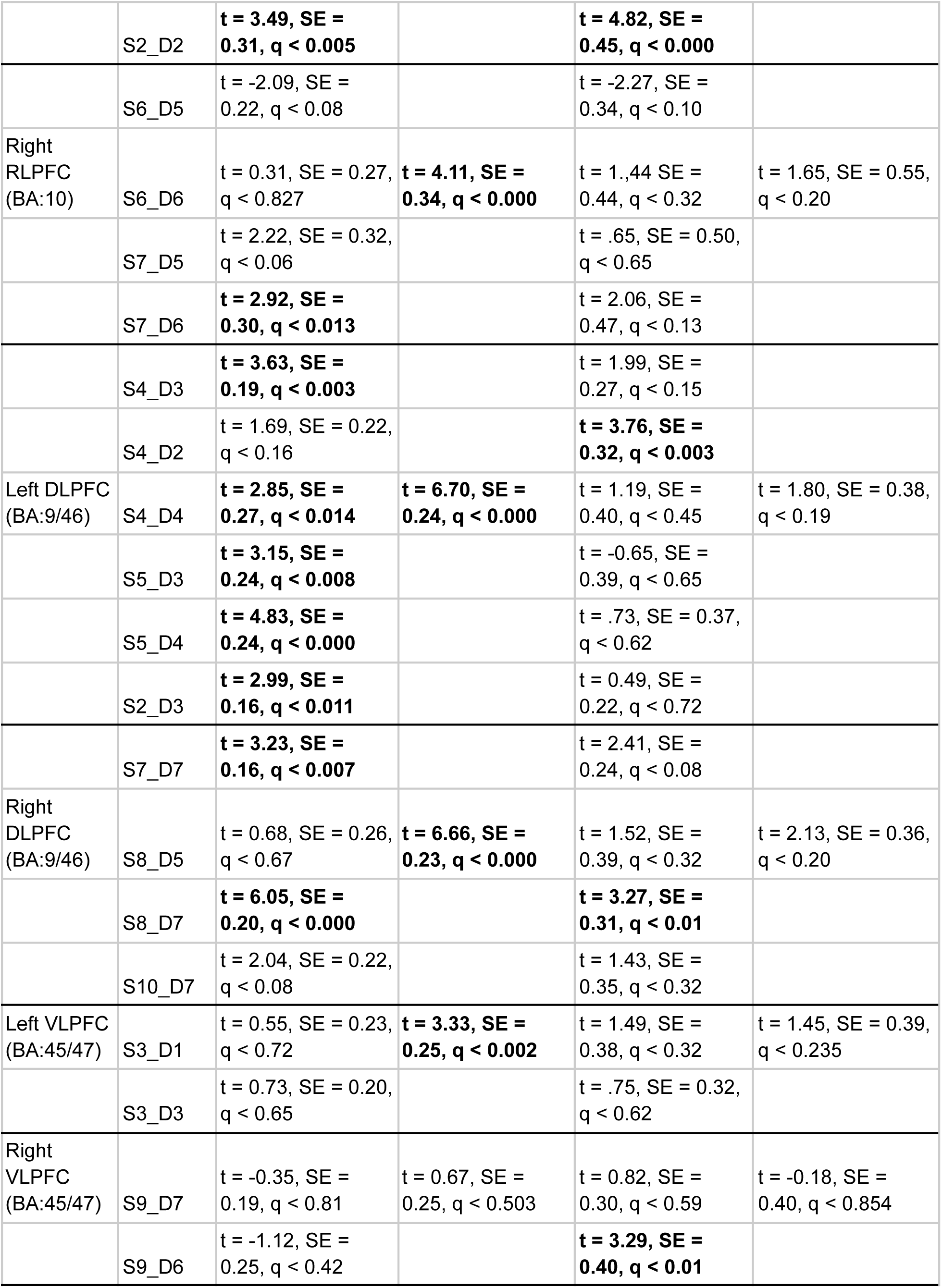
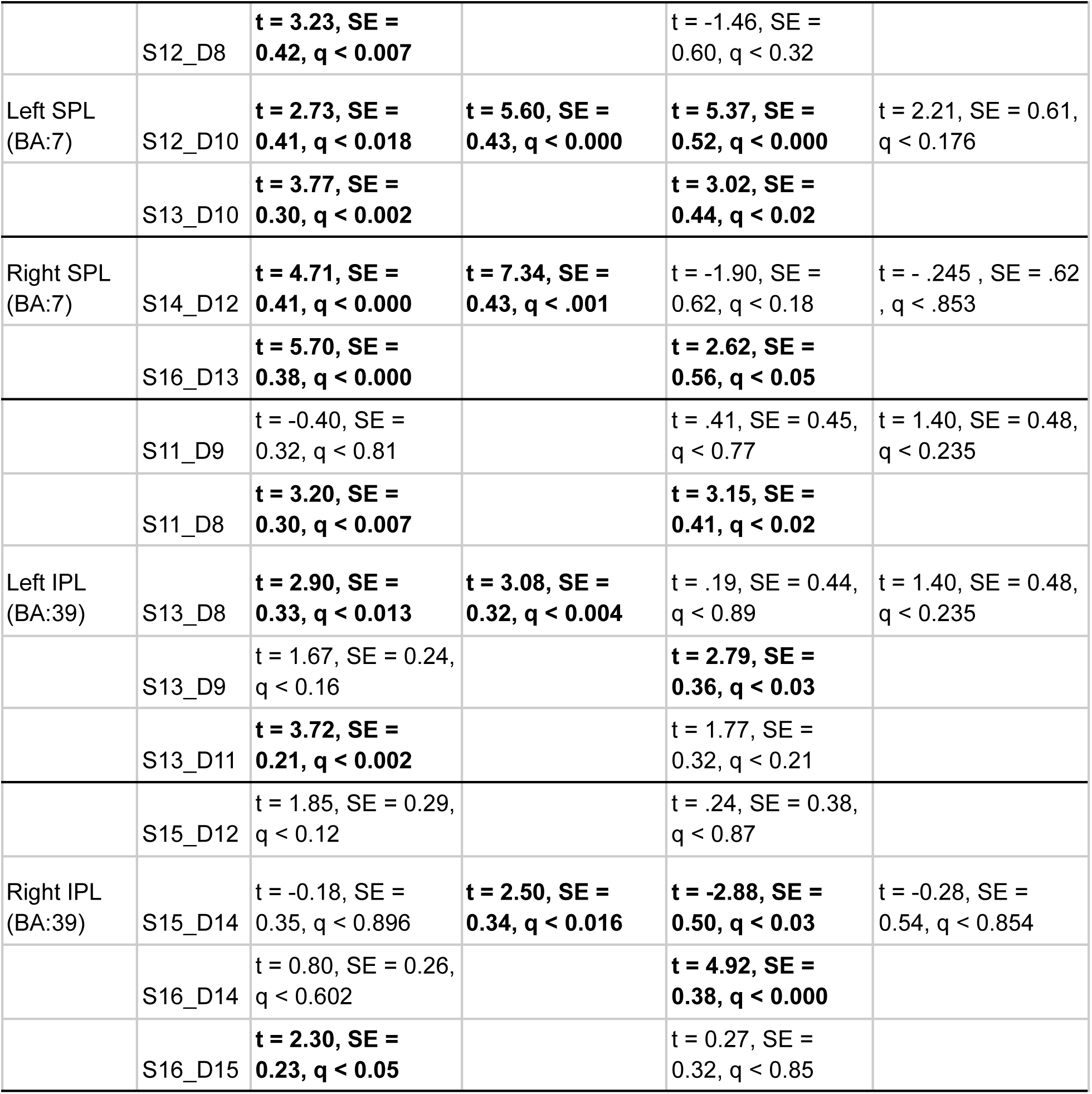
Channel and ROI level task activation statistics for both the Task vs. Baseline and 2nd vs. 1st contrasts. Bolded values reached significance and survived multiple correction (q<.05)

#### 4.2.2 Relational Complexity Manipulation

Task-evoked activation at the channel level was significantly greater for 2nd-order relative to 1st-order reasoning in 12 channels distributed across bilateral prefrontal and parietal channels, correcting for multiple comparisons. Robust effects were observed for several channels in bilateral RLPFC and DLPFC, with additional engagement in right VLPFC. Beyond PFC, several channels in left and right SPL and IPL were also significant (see **Figure 4** and **Table 1**). By contrast with the channel-level analyses, no ROIs demonstrated statistically significant differences between task conditions after correction for multiple comparisons.

**Figure 4.**
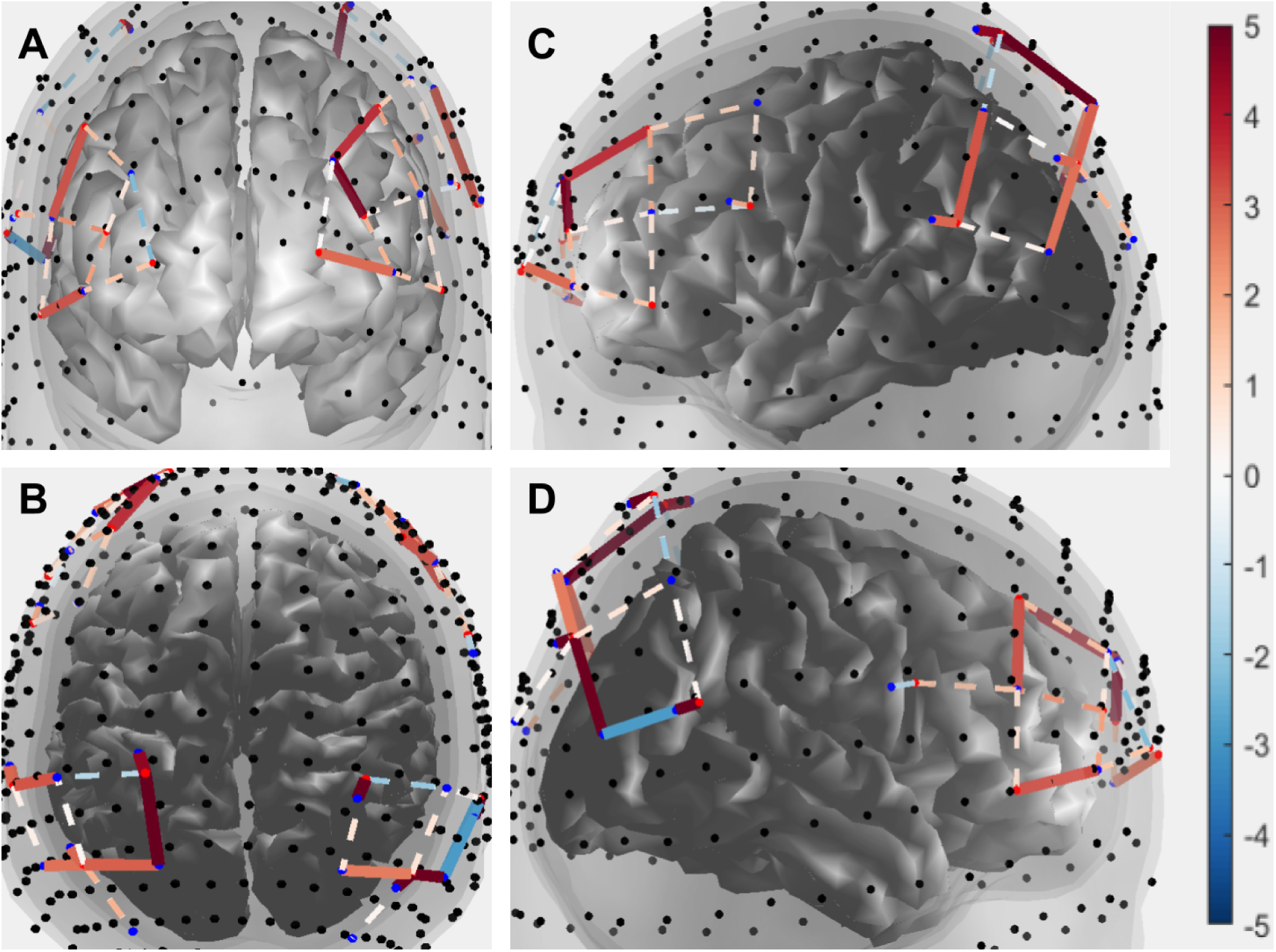
Channel level 2nd vs.1st-order contrast for task activation across participants.. Solids lines indicate channels that were significant after correcting for multiple comparisons across 36 channels; and dashed lines indicate those that did not survive correction. The color bar indicates t-statistic ranging from 5 to -5 . **A.** Anterior view of brain showing significant activation in channels over left RLPFC, left DLPFC, and right VLPFC. **B**. Dorsal view of brain, showing significant activation in channels over bilateral IPL and SPL. **C.** Lateral view of the left hemisphere, showing significant activation in channels over left RLPFC and DLPFC, as well as left IPL and SPL. **D**. Lateral view of the right hemisphere, showing significant activation in channels over right DLPFC, VLPFC, and IPL.

#### 4.2.3 fNIRS and fMRI Effect Size Comparison

As a final step, we sought to get a rough sense of how robust our fNIRS results were relative to fMRI studies using similar relational matching paradigms (Christoff et al., 2003; Smith et al., 2007; Bunge et al., 2009; Dumontheil et al., 2010; Wendelken et al., 2011a; Wendelken et al., 2011b; see **Table 2**). Of these, we selected the three studies that reported a group analysis of task activation based on an adult sample; all three were based on data collected at 3 Tesla (Christoff et al., 2003; Bunge et al., 2009; Wendelken et al., 2011a).

Focusing on effect sizes reported in RLPFC for 2nd > 1st-order trials, we documented the mean and range of reported – or converted – t-values across channels (for fNIRS) or clusters of voxels (for fMRI) that survived correction for multiple comparisons in the manner specified in these studies. Pooling results from these 3 fMRI studies, t-values for the relational complexity manipulation ranged from 2.62 - 6.65 across the 8 significant clusters in RLPFC. By comparison, t-values for this comparison in the present study ranged from 2.88 - 4.82 across the 2 channels in LPFC with significantly stronger activation for 2nd vs. 1st-order blocks, correcting for multiple comparisons across all 8 channels. Thus, while there are numerous differences among the studies in terms of sample sizes (with the present one being the largest), measurement granularity (channel-level vs. voxel-level), statistical thresholding approach, areas of LPFC reported, and details of the study design, reported effect sizes between the present fNIRS study and prior fMRI studies are comparable.

**Table 2.** Comparison of effect sizes in RLPFC across relational matching neuroimaging studies. N refers to the number of participants. The mean and range of t-values are reported for significant RLPFC clusters (or channels, in the present study). In Christoff et al., 2009, the 2nd > 1st-order comparison corresponds to the no-load internally > externally generated contrast; within an anatomically defined ROI of bilateral RLPFC, significant activation is identified for a single cluster in left RLPFC with an average voxel-level p < .05 (Z > 3.28), correcting for multiple comparisons. In Bunge et al., 2009, the 2nd > 1st-order task contrast yields clusters in left and right RLPFC that survive correction for multiple comparisons (p < .05) across voxels located anterior to the MNI coordinate of y = +48. In Wendelken et al., 2011, the 2nd > 1st-order comparison for the visuospatial relational matching task (similar to the present task) identifies a cluster in left RLPFC that survives correction for multiple comparisons (p < .05) across the whole brain. In the present study, two RLPFC channels, both in the left hemisphere, survive correction for multiple comparisons across all 8 RLPFC channels (q < .05).

| Study | Age range | N | RLPFC activations | Mean T-value (min, max) |
| --- | --- | --- | --- | --- |
| Christoff et al., 2003 | 18-25 | 12 | left RLPFC | 2.74 |
| Bunge et al., 2009 | 19-35 | 15 | left & right RLPFC | 2.86 (2.62, 3.33) |
| Wendelken et al., 2011 | Young adult | 22 | left RLPFC | 6.65 |
| Current study | 18-29 | 73 | left RLPFC | 3.85 (2.88, 4.82) |

### 4.3 Effects of Number of Task Blocks

#### 4.3.1 Behavioral Data

First, we examined whether or how task performance changed across the session – looking for hints of participant fatigue (decreased accuracy and/or increased RTs) or learning (the converse pattern) – with a view to aiding the interpretation of effects of block count on fNIRS activation.

To explore changes in task performance across the ∼15-minute experiment, given prior evidence of practice-related improvements in task-switching efficiency over time (Karbach et al., 2009; Schwarze et al., 2025), we examined whether/how performance changed across the 18 individual task blocks (**Supplementary Figure 5**). Accuracy remained consistently high on 2nd-order blocks throughout the session, indicating strong and sustained task engagement. In contrast, accuracy on 1st-order blocks showed an early dip during the first few blocks, which could reflect initial difficulty adapting to the demands of task-switching between color- and shape-based rules, and then gradually improved and stabilized for both conditions around block 9, suggesting successful adjustment and ongoing engagement. RTs for both conditions also leveled off by approximately block 9—rising early for 1st-order trials and declining for 2nd-order trials—further supporting the interpretation that participants adapted to the task demands as the session progressed, but providing no evidence of task disengagement towards the end.

#### 4.3.2 Task Activation

We visualized channel-level activation patterns at the group level in increments of three cumulative blocks – that is, the first 3, 6, 9, 12, or 15 blocks, or all 18 blocks (**Figure 5**). Additionally, we aggregated over channels to provide numerical comparisons of average channel-level beta values and SE across these increments (**Figure 6**). We also examined how the significance of each individual channel changed status (where significance status was defined as activated, deactivated, or not significant, based on our threshold of q < .05) as a function of number blocks for both contrasts (**Supplementary Figures 7 & 8**). In addition to examining the cumulative effect of task blocks included, we conducted complementary analyses comparing the first and second halves of the session (i.e., independent sets of data (**Supplementary Section 9**).

**Figure 5.**
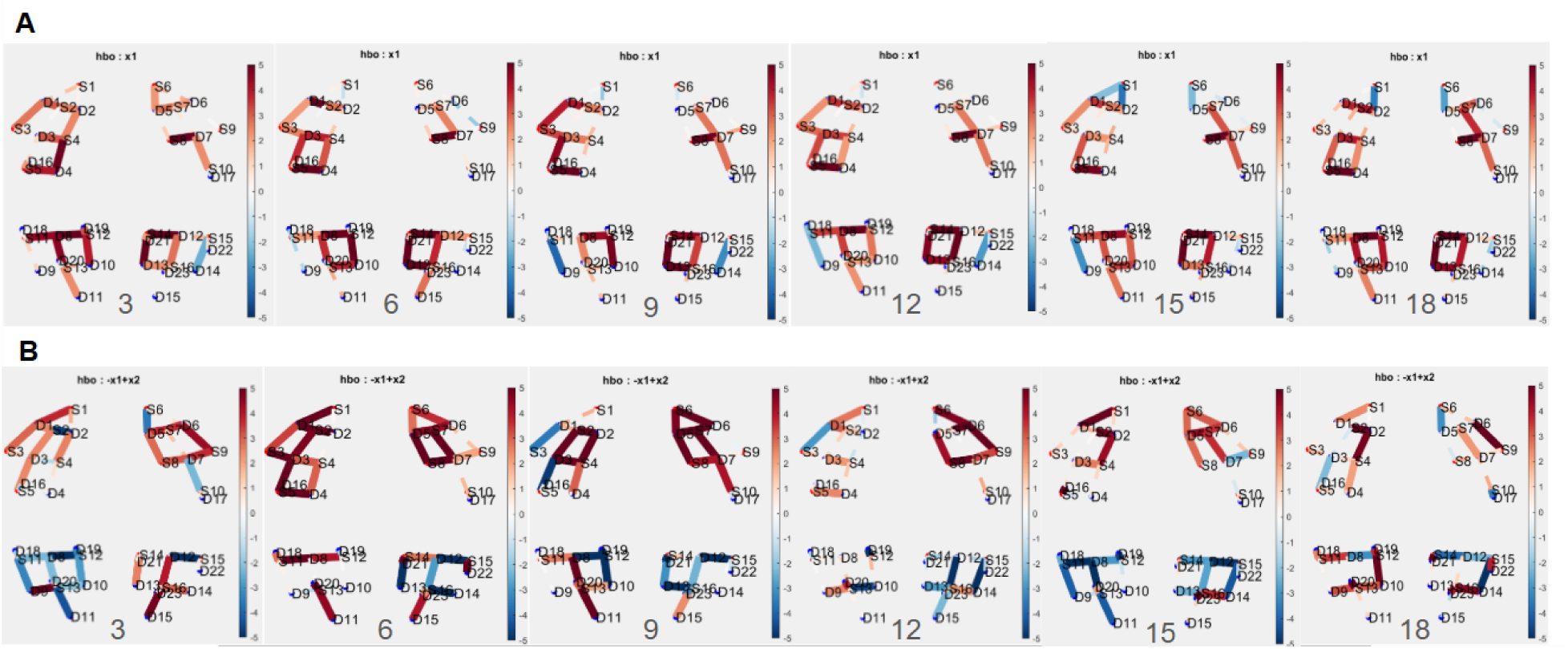
Group level activation plots for cumulative block counts of 3,6,9,12,15,and 18 for **A.** the Task vs. Rest contrast and **B.** 2nd vs.1st-order contrast. Color bars indicate magnitude of t-statistic ranging from 5 to -5.

The Task vs. Baseline activation topographic maps suggest a fair degree of stability in the pattern of activation in the aggregate across the 3-block increments (**Figure 5A**); this qualitative observation is supported by relative stability in channel-level significance status (positive, negative, or no activation at q < .05; **Supplementary Figure 7**). Quantitatively, the mean group-level beta coefficients and SE both trended downwards across block counts, suggesting improved sensitivity to detect overall task activation with more data, but with diminishing returns after 9 blocks (see **Figure 6A**).

**Figure 6.**
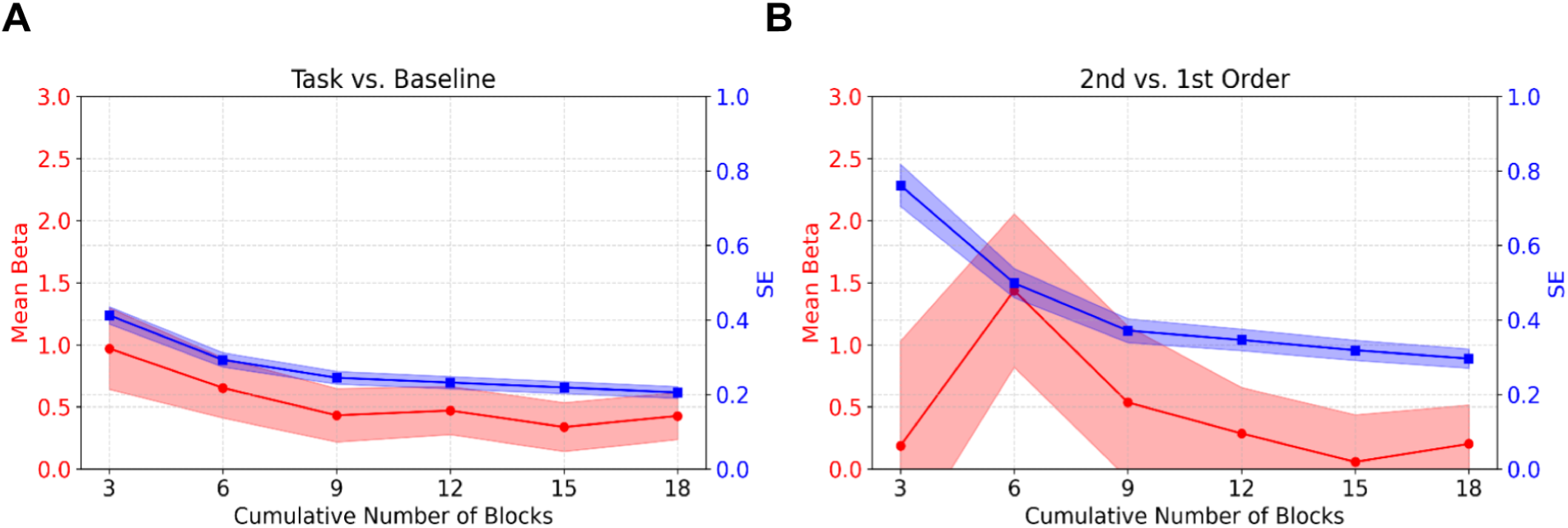
Mean beta values (in red) and SE (in blue) across channels as a function of block count, for the first 3, 6, 9, 12, or 15 blocks or all 18 blocks, for **A.** Task vs.Baseline and **B.** the 2nd vs. 1st-order contrast.

For the 2nd vs. 1st-order contrast, the topographic plots hint at a lower degree of stability in the pattern of activation across the 3-block increments (**Figure 5B**); indeed, many channels changed in significance status across the session for this contrast (**Supplementary Figure 8**). On average, mean beta values peaked around block 6 and then decreased, while SE declined (**Figure 6B**). These aggregated results suggest that statistical estimates for this contrast became more precise as more task data were included, improving precision and a stabilization of condition-specific effect estimates and allowing the model to more accurately distinguish between nuanced differences between conditions. That said, these results must be considered in light of low stability in significance status at the channel level. Moreover, SE values for the specific contrast were higher at the end of the experiment than for the general contrast; reflecting the general instability of condition level contrasts in neuroimaging research (Heilicher et al., 2022; DeYoung et al., 2025), and/or the possibility that additional blocks of each condition could result in higher stability.

Together, the block count analyses for both contrasts suggest that, while smaller numbers of blocks may produce artificially elevated variability and effect sizes, using at least 9 blocks improved the stability of group-level contrast maps. Behaviorally, too, accuracy and RTs stabilized after approximately 9 blocks (**Supplementary Figure 5**). That said, estimates of task activation became increasingly stable with 15–18 blocks, as indicated by the tapering SE and the convergence of mean beta values. Importantly, however, individual channel-level results reveal marked instability in the 2nd > 1st-order contrast, with substantial flux in significance status across blocks.

### 4.4 Functional Connectivity

Functional connectivity analyses were conducted at the channel- and ROI-levels for the subset of 36 participants with the highest channel signal quality, as well as on the ROI-level for both this subset (**Figure 7**) and the full sample of 73 (**Figure 8**). As functional connectivity patterns were nearly identical between 1st- and 2nd-order blocks in the full sample (channel level: *r* = .997 (*p* < 1.00 × 10^-30^); ROI level: *r* = .995 (*p* < 1.00 × 10^-30^), all subsequent connectivity analyses were conducted collapsing across conditions.

At the channel level (**Figure 7**), task-evoked functional connectivity revealed extensive synchronization across bilateral LPFC and PPC; particularly among channels within an ROI, but also between ROIs. At the ROI level (**Figure 8**), the strongest pairwise connectivity for the full sample was found among PFC channels – in particular between left RLPFC and left VLPFC, and between left and right RLPFC – and to a lesser extent within PPC. Both the channel- and ROI-level analyses converged in identifying a coherent frontoparietal network characterized by bilateral and long-range connections. ROI-level connectivity patterns observed in the 36 participants with the highest quality data closely mirrored that of the full sample of 73 (**Supplementary Figure 6**), highlighting the potential to exclude substantially fewer participants when aggregating across channels by dropping bad channels from the ROI estimates. Thus, while channel-level analysis provided finer spatial resolution and revealed specific node-level interactions, aggregation of channels into ROIs enabled robust connectivity estimation across the full sample while excluding poor-quality channels.

**Figure 7.**
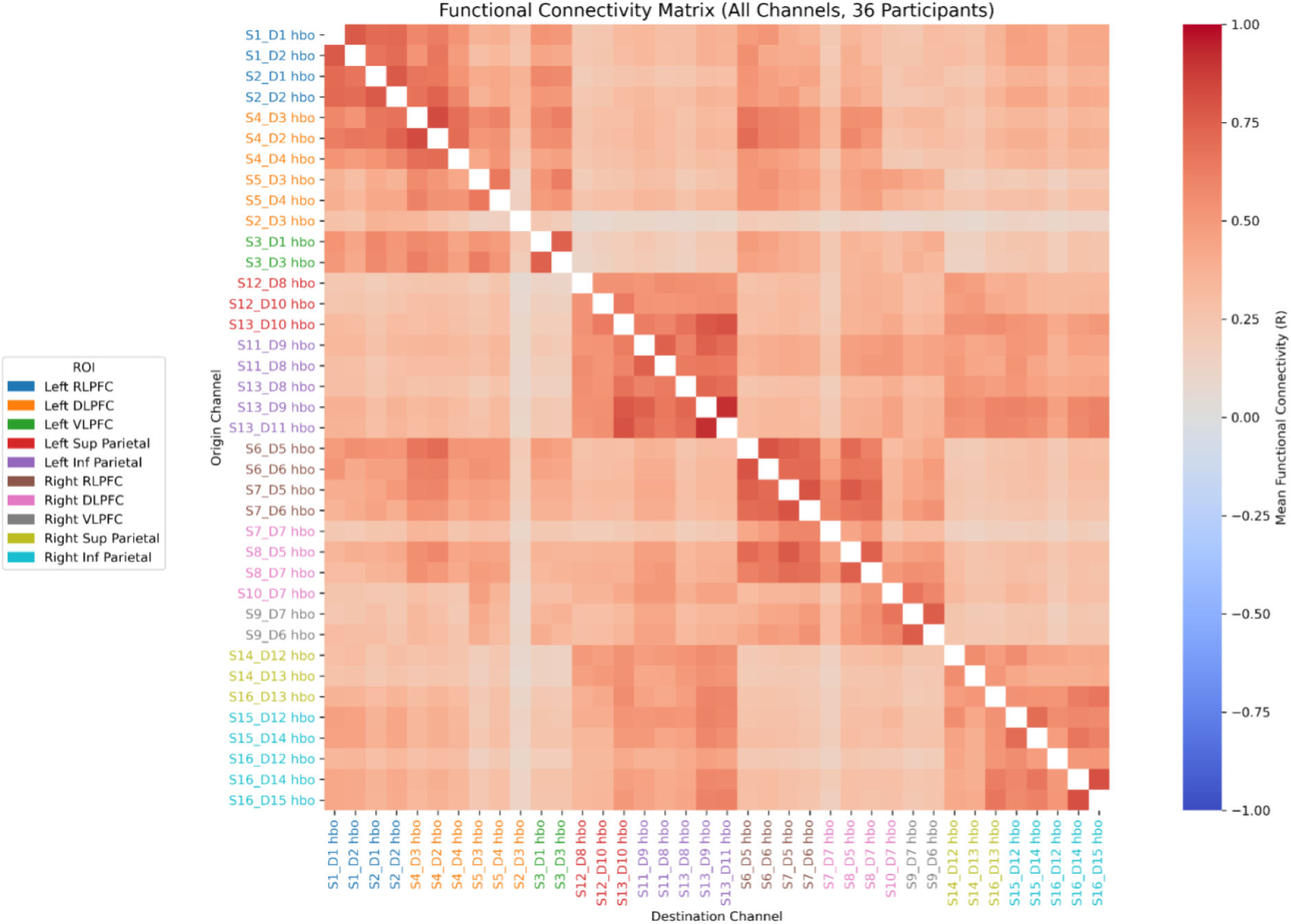
Channel x Channel functional connectivity matrix on the subset of participants with highest quality data (N =36).

**Figure 8.**
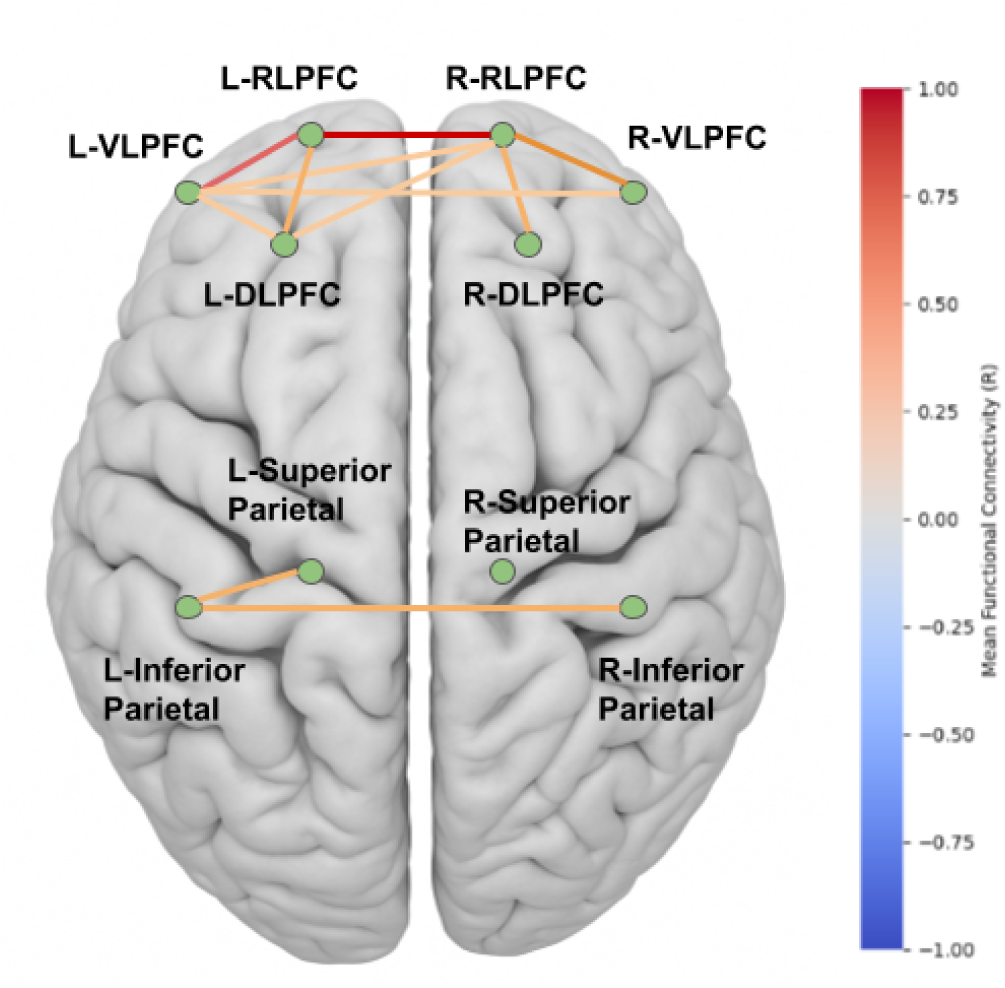
Dorsal view schematic illustrating the strongest patterns of pairwise connectivity (r >.3) in ROI x ROI functional connectivity on the full data set (N = 73).

### 4.5 Within-Session Stability

#### 4.5.1 Task Activation

We assessed within-session internal consistency reliability of task-evoked activation within individuals with beta values across interleaved (odd vs. even) blocks, both at the channel and ROI levels. At the channel level, the Task vs. Baseline contrast showed moderate internal consistency reliability on average (ICC = .54), with additional appreciable variance attributable to channel (ICC = .07) and the interaction between channel and subject (ICC = .12). Indeed, there was substantial variation in ICC values across channels, ranging from ICC = .00 to .79 (**Figure 9A**). On average, individual channels in LPFC (M_ICC_ = .63) had significantly higher ICC values than channels in PPC (M_ICC_ = .49; t(34) = 2.84, *p* = .008). At the ROI level, the Task vs. Baseline contrast showed moderate internal consistency reliability and substantial variance across ROIs in reliability estimates (see **Supplementary Section 10.2**).

By comparison with Task vs. Baseline, the 2nd vs. 1st-order contrast showed lower reliability (ICC = .44; **Figure 9B**), with little-to-no variance attributable to channel (ICC = .00) or the interaction between channel and subject (ICC = .00). When calculating reliability per-channel, there was a wide range of ICC values (see **Supplementary Figure 11**; range = .00 to .54). On average, individual channels in LPFC (M_ICC_ = .18) had similar ICC values to channels in PPC (M_ICC_ = .24; (t(34) = -1.16, *p* = .26). At the ROI level, the 2nd vs.1st-order contrast again showed lower reliability, with substantial variance across ROIs in reliability estimates (see **Supplementary Section 10.2**). Overall, the Task vs. Baseline contrast was more reliable than the 2nd vs. 1st-order contrast, and demonstrated moderate within-session reliability across individuals with channel-level metrics consistently outperforming ROI-level estimates; however, these internal consistency estimates varied substantially across channels.

**Figure 9.**
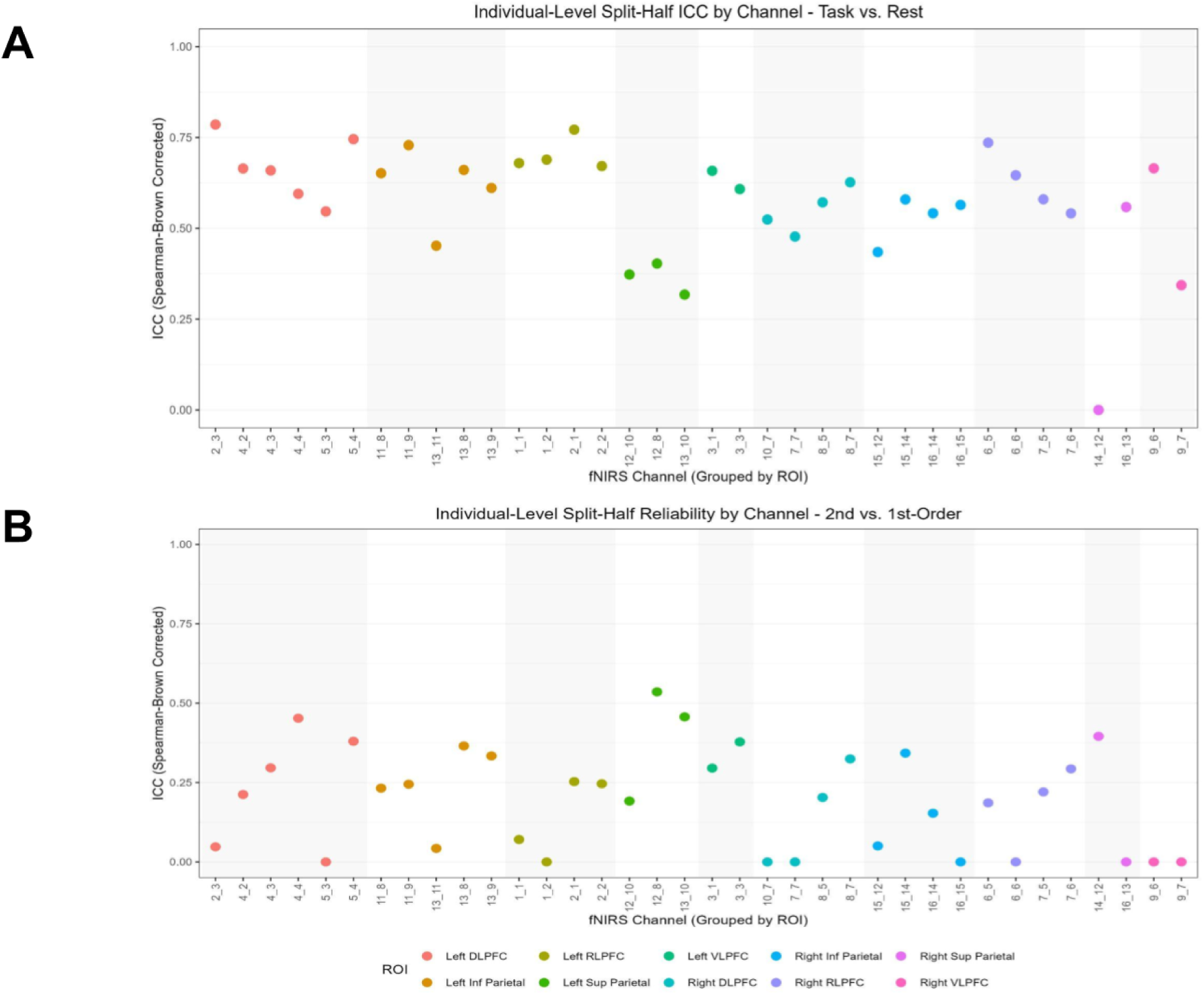
Within-session participant level split half reliability using interleaved blocks for **A**. Task vs.Baseline **B**. 2nd vs. 1st-order contrast. Spearman-Brown corrected ICC values are plotted for each channel, grouped and color-coded by brain region.

We also assessed internal consistency reliability at the group level, for both the channel and ROI spatial scales. Group-level results at the channel level for the Task vs. Baseline contrast showed high within-session spatial stability for interleaved blocks (r = .60, ρ = .75, variance = .97 p < .001). However, the 2nd vs. 1st-order contrast for interleaved blocks showed low spatial stability (r = .06, ρ =.12, variance = .44, p = .71) (see **Supplementary Section 10.1)**. At the ROI level, Task vs. Baseline demonstrated near perfect internal spatial consistency (r = .89, ρ = .94, variance = .87, *p* < .001), while the 2nd vs.1st-order yielded zero reliability (r = –.31, ρ = –.91, variance = .18, *p* = .38; **Supplementary Section 10.2**). Thus, aggregating data across channels in an ROI and across individuals resulted in very high stability for the general contrast but the specific task contrast was still unstable.

#### 4.5.2 Functional Connectivity

To assess the internal consistency of task-based functional connectivity within a session at the individual and group levels, we computed Pearson correlations between connectivity matrices derived from odd and even task blocks, separately for channel-level and ROI-level analyses.

Individual-level within-session stability was extremely high across both spatial scales. For the 36-participant subset, channel-level connectivity correlations between odd and even blocks ranged from r = 0.944 to 0.986 (mean = 0.97, median = 0.97, SD = 0.01), with all participants exceeding r = 0.94. This consistency reflects robust preservation of fine-grained network patterns within individuals over the course of the session (see **Supplementary Section 10.3**).

Group-level analyses also revealed exceptionally high reliability at both spatial scales for both the channel and ROI-levels (see **Supplementary Section 10.3**). At the channel level, the correlation between connectivity matrices across all 630 unique channel pairs was r = 0.998 (p = 2.36 × 10^-4^, permutation test), indicating near-identical network structures between odd and even blocks. Thus, functional connectivity showed very high stability within a session.

### 4.6 Between-Session Stability

#### 4.6.1 Behavioral Data

To confirm participant engagement and assess task performance across conditions, we examined accuracy and RTs for 1st-order and 2nd-order trials across sessions 1 and 2. All participants exceeded the pre-established performance threshold of 70% accuracy in both sessions, and were therefore included in the final analysis. As in session 1, accuracy in session 2 was high (1st-order = 91.27%, 2nd-order = 95.36%), and there was a trend towards improvement from session 1 to 2. Due to limited inter-individual variability in accuracy, we did not conduct a correlation analysis to assess stability. However, RTs showed strong within-subject stability across sessions, with strong correlations for both 1st-order (r = .75, p < .001) and 2nd-order (r = .86, p < .001) conditions. (see **Figure 10**). For full statistics see Supplementary Section 11.1.

**Figure 10.**
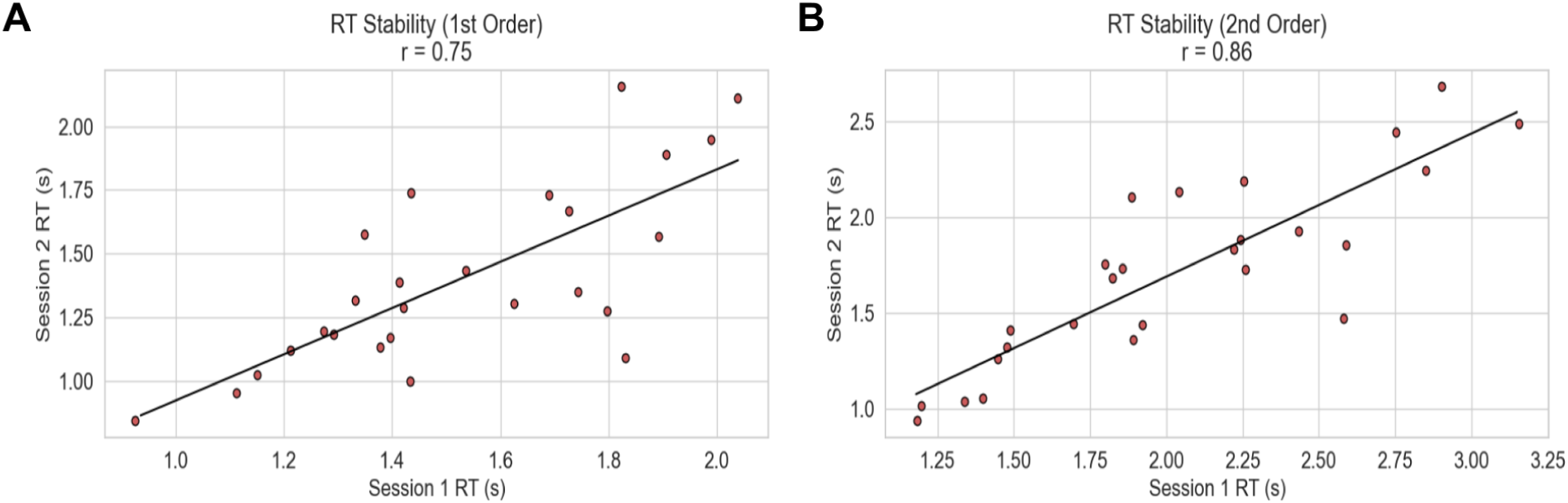
Between session RT stability for **A.** 1st-order and **B.** 2nd-order conditions. Individual data points correspond to average RTs for each individual at session 2 plotted against session 1.

#### 4.6.2 Task Activation

##### Individual-Level

To assess the test-retest reliability of task-evoked neural responses across sessions, we examined the stability of beta values across sessions at the channel and ROI levels within participants. At the channel level, the Task vs. Baseline contrast showed relatively low test-retest reliability within participants (ICC = .18), with appreciable variance attributable to channel (ICC = .07) and to the interaction between channel and subject (ICC = .05). Despite the low overall average test-retest reliability, a wide range was observed, ICCs = .00 - .65, and there were several channels with good test-retest reliability estimates (see **Figure 11A**). On average, individual channels in LPFC (M_ICC_ = .35) showed significantly higher test-retest reliability than channels in PPC (M_ICC_ = .21) (t(34) = 2.23, *p* = .03). Like the channel-level results for Task vs. Baseline, the 2nd vs. 1st-order contrast showed low test-retest reliability (ICC = .13), with very little-to-no variance attributable to channel (ICC = .00) or the channel by subject interaction (ICC = .00). When examining differences across channels in reliability, over half of the channels showed no reliability, but there were some exceptions, ICCs = .00 - .45 (see **Figure 11B**). Aggregating at the ROI level yielded similar results: both contrasts showed relatively low test-retest reliability, but there were notable differences across ROIs in these reliability estimates (see **Supplementary Section 11.2.1**).

**Figure 11.**
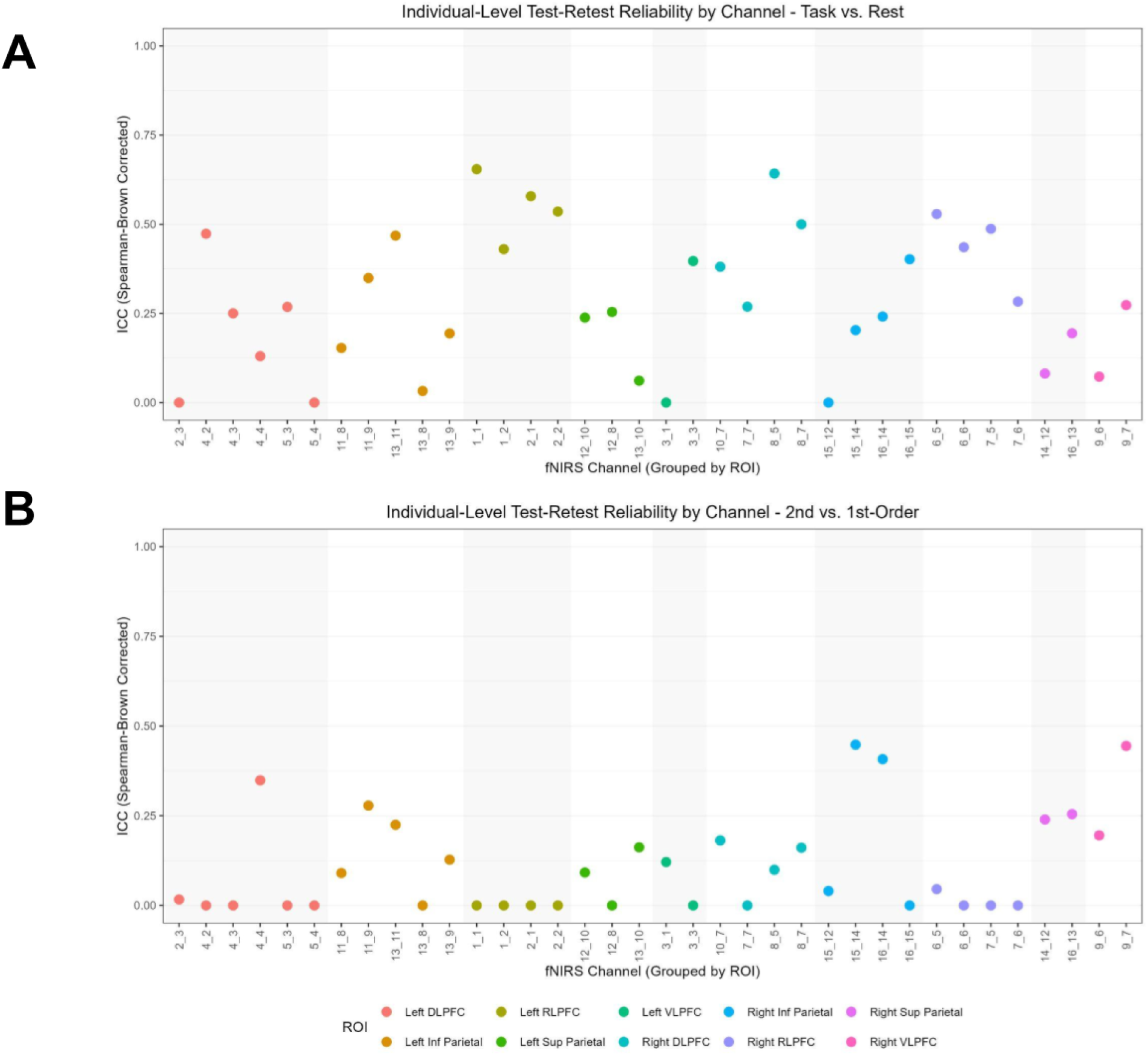
Between-session participant level stability for **A**. Task vs. Baseline contrast indicating moderate reliability for channels in bilateral RLPFC and right DLPFC, and **B.** 2nd vs. 1st-order contrast, indicating low reliability for most channels

Overall, both contrasts demonstrated low test-retest reliability across both spatial scales at the participant level, with channel-level metrics consistently outperforming ROI-level estimates. Task vs. Baseline contrasts showed higher internal consistency reliability than the 2nd vs. 1st-order contrast at the ROI level, but the opposite was true at the channel level. While ROI aggregation improves anatomical interpretability, it may reduce sensitivity to stable individual differences—especially for condition-specific contrasts—underscoring the utility of fine-grained channel-wise analysis in reliability-focused fNIRS studies.

##### Group-Level

Although test-retest reliability was low at the individual participant level, we sought to assess test–retest reliability of the group-level activation pattern. To this end, we examined both the spatial consistency of t-values and the stability of activation variance (t²) across sessions and the difference in amplitude from session to session, both at the channel and ROI levels. At the channel level, spatial activation patterns were moderately stable for Task vs. Baseline, but not for the 2nd vs. 1st-order contrast (**Figure 12**). Variance stability had the opposite pattern, with nonsignificant correlations for Task vs. Baseline and high 2nd vs. 1st-order contrast variance stability. At the ROI level (**Supplementary Section 11.2.1**), the Task vs. Baseline contrast showed even greater test–retest reliability (r = .87, variance = .93, p < .001), while the 2nd vs. 1st-order contrast remained unstable (r = .28, variance = .31, p = .44) (**Supplementary Figure 12**).

**Figure 12.**
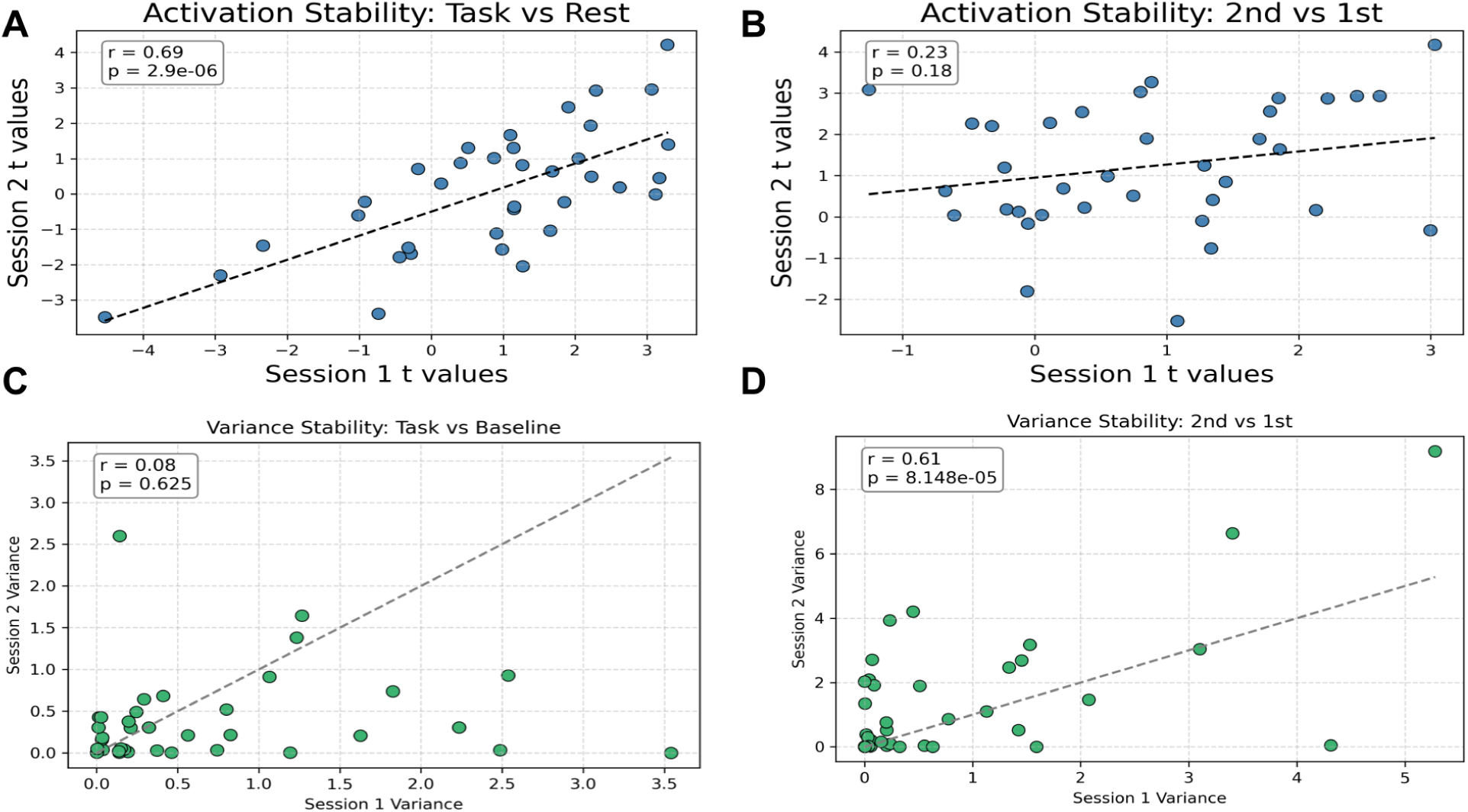
Group-level activation and variance stability data at the channel level. **A.** Task vs.Baseline correlation across sessions indicating strong correlations between sessions (r = .69) **B.** 2nd vs.1st correlation across sessions, indicating very low correlation between sessions (r = .23) **C.** Task vs. Baseline variance correlation across sessions, indicating low correlation between sessions (r = .08) **D.** 2nd vs.1st variance correlation across sessions, indicating moderate correlation between sessions (r = .61)

With regards to overall signal amplitude, we observed higher channel-level amplitude in session 1 than session 2 for the Task vs, Baseline contrast (t = 4.03, p < .001). The 2nd vs.1st-order contrast showed the opposite pattern; however, these results were not significant (t = -1.28, p = .21). At the ROI level, the pattern of signal amplitude differed across sessions for both the Task vs. Baseline contrast (t = 3.96, p = .004) and the 2nd vs. 1st-order contrast (t = 4.90, p = .001), indicating higher ROI level activity in session 1 than session 2. In sum, both group-level spatial and variance-based metrics indicate that ROI-level activation estimates demonstrate greater test–retest reliability than channel-level data for overall task activation; both were poor for the specific task contrast. While channel-level analyses may offer higher spatial resolution and sensitivity to localized effects, they appear to be less stable across sessions at the group level, especially for fine-grained condition contrasts.

#### 4.6.3 Functional Connectivity Stability

To assess the test–retest reliability of individual functional connectivity, we computed Pearson correlations between each participant’s session 1 and session 2 connectivity matrices separately at the channel and ROI levels. At the channel level, subject-wise correlations across the 26 participants showed high consistency across all conditions. For 1st-order, correlations ranged from r = .17 to .95 (mean+/- SD: .73 +/- .12); for 2nd-order, from r = 0.18 to 0.94 (.74 +/- .11); and for Task vs. Baseline, from r = 0.23 to 0.95 (.70 +/- .12). All correlations were statistically significant (p < 0.001, permutation test), indicating robust individual-level stability of channel-wise functional connectivity (**Figure 13B**). At the ROI level, functional connectivity demonstrated even greater stability (**Supplementary Section 11.3.2**). To summarize, ROI-level connectivity exhibited slightly higher test–retest reliability than channel-level data, with tighter correlation ranges and greater central tendency. This pattern confirms the robustness of ROI-level connectivity as a stable and interpretable marker of between-session functional architecture. Both approaches, however, confirmed strong within-subject stability, supporting their utility for longitudinal designs and individualized network characterization.

Group-level test-retest reliability for functional connectivity at the channel level was also extremely strong (r = 0.97, p < 1 × 10^-66^) (**Figure 13A; Supplementary Section 11.3.1**). At the ROI level, similarly high stability was observed (**Supplementary Section 11.3.2** and **Supplementary Figure 14**).

**Figure 13.**
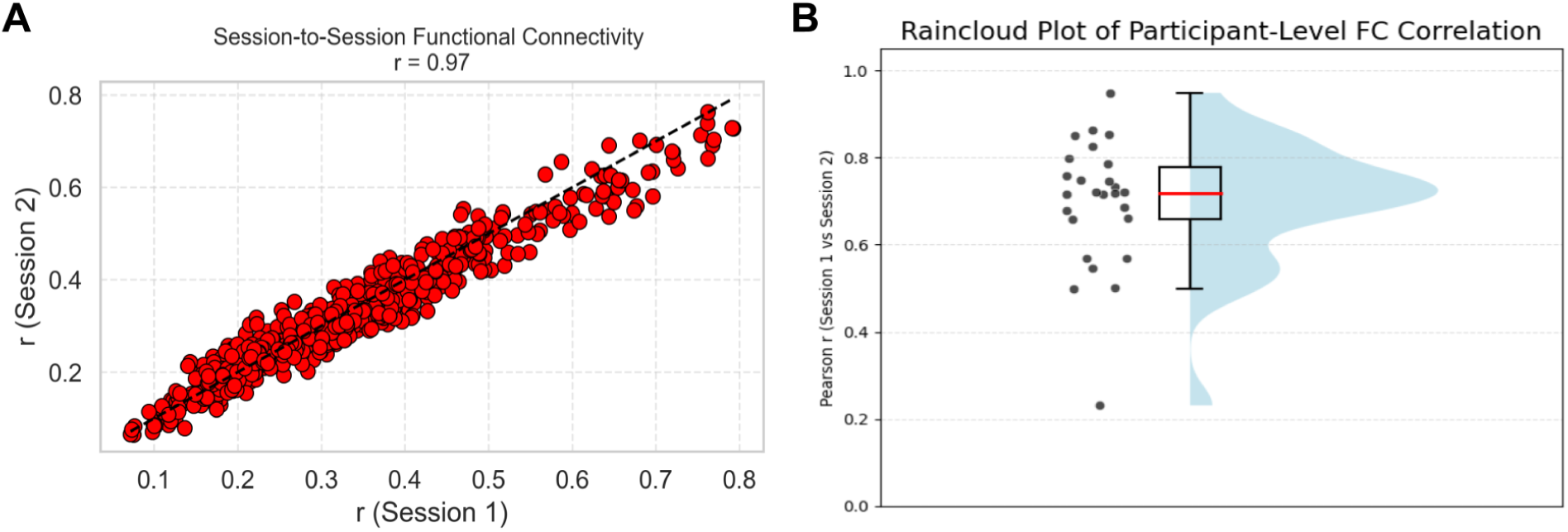
Channelwise functional connectivity correlations between sessions at **A.** Group-level (r = .97, p = 2.28 x 10^-66^) and **B.** Participant level functional connectivity correlations between sessions, ranging from .23 to .95 (mean =.7)

### 4.7 Exploratory Analysis: Effects of Hair and Skin Types

An exploratory analysis was conducted to assess whether phenotypic characteristics—specifically skin tone, hair color, hair type, and hair texture—predicted fNIRS signal quality, as documented in prior studies (Holmes et al., 2024; Simmons et al., 2025; Yücel et al., 2024). For full methods see **Supplementary Section 12**.

Skin tone significantly predicted bad channel count, F(3, 50) = 4.07, p = .012. Post hoc Tukey’s HSD tests revealed that participants with brown skin had more bad channels than those with fair skin (mean difference = 10.92, p = .082), and participants with medium skin showed a similar trend (mean difference = 6.01, p = .084), though both effects were marginally significant. Hair color was also a significant predictor of signal loss, F(2, 50) = 6.71, p = .003. Participants with red or black hair had more bad channels than those with blonde hair (mean difference = 10.56, p = .007) and brown hair (mean difference = 5.31, p = .027), while the difference between brown and blonde hair was not significant (p = .297). In contrast, neither hair type (F = 0.93, p = .43) nor hair texture (F = 0.045, p = .64) significantly predicted the number of bad channels (**Supplementary Figure 15**).

Together, these findings suggest that certain phenotypic features—particularly darker skin tones and non-blonde hair colors—are associated with greater fNIRS signal loss. Although limited by small and uneven subgroup sizes, this pattern supports prior research on melanin- and hair-related light attenuation and emphasizes the need for inclusive methodological practices and better-powered future studies examining phenotypic bias in optical neuroimaging.

## 5. Discussion

In the current study, we sought to evaluate the reliability of fNIRS data on a sample of healthy adults during a reasoning task, with a view toward extending it to pediatric populations. While fNIRS technology has great potential for advancing developmental cognitive neuroscience due to its unique advantages of robustness to motion artifacts and environmental noise, making it particularly suitable for applications beyond traditional laboratory settings (Huppert et al., 2006; Pinti et al., 2018) and for studying hard-to-test populations (Lloyd-Fox et al., 2010), the signal quality and neurometric properties of the signal need to be comprehensively evaluated. After using fNIRS to corroborate results from similar fMRI paradigms, we found that signal quality and reliability (both within-session and across-session) varied across different quantification techniques and spatial scales.

The instability observed in condition level contrasts (2nd vs.1st-order), especially at the individual level, is largely attributed to their computation as “difference scores”. This analytical approach can introduce a high degree of measurement error and consequently result in lower reliability estimates. These same analytical and methodological concerns have been shown in previous fMRI research, resulting in similarly low reliability for contrast-based ROI work. Previous research has suggested that using condition specific beta (β) coefficients, which directly characterize percent signal change, yields greater reliability values, suggesting that previous reports of low fMRI test-retest reliability may stem from methodological considerations in data analysis rather than an intrinsic limitation of the modality itself (Heilicher et al., 2022; DeYoung et al., 2025).

Given known anatomical and physiological challenges in optode-scalp coupling over posterior regions (Beauchamp et al., 2011; Tachtsidis et al., 2016; Skau et al., 2022; Klein, J.H., 2024), particular attention was given to data quality in PPC. Four channels located over right SPL (S14_D13, S14_D21, S16_D12, S16_D23) were excluded due to their poor signal quality and three channels located over left SPL consistently yielded signal quality values near or at the exclusion threshold across multiple participants. Additionally, PPC showed lower reliability in both within and between session metrics compared to LPFC. This reflects a broader issue in fNIRS research: PPC is notoriously difficult to image reliably, due to factors such as greater hair density, increased curvature of the posterior scalp, and a thicker section of skull, all of which attenuates near-infrared light transmission. These factors contribute to higher rates of signal dropout and greater variability in signal-to-noise ratio in parietal areas compared to more anterior regions like PFC. As a result, although these superior parietal channels were retained in the final montage to preserve anatomical coverage and network completeness, their inclusion warrants caution when interpreting activation or connectivity findings involving this region. The challenges underscore the need for region-specific quality checks in fNIRS preprocessing pipelines and reinforce the growing consensus that some cortical areas remain technically constrained for optical neuroimaging.

### 5.1 Task-Based Activation

Task performance reliably engaged widespread frontoparietal regions, with the Task vs. Baseline contrast revealing significant activation in 20 of 36 channels spanning bilateral PFC and PPC: most notably bilateral RLPFC and DLFC. When aggregating across channels, significant activation was observed in all ROIs: bilateral RLPFC, DLPFC, VLPFC, and parietal cortices, underscoring the value of ROI-level analysis in detecting broad patterns of engagement by increasing statistical power and reducing noise via spatial averaging.

The more selective 2nd vs. 1st-order contrast elicited activation in a subset of 12 channels, predominantly in the left hemisphere. These activations were qualitatively similar to those of prior fMRI studies of relational reasoning, particularly the robust activation observed in left RLPFC (Smith et al., 2007; Bunge et al., 2009; Dumontheil et al., 2010; Wendelken et al. 2011; Wendelken et al., 2011; Christoff et al., 2020; see Vendetti & Bunge, 2014). Moreover, the t-values observed in left RLPFC in our fNIRS data (2.88–4.82) were similar in magnitude to those reported in comparable fMRI studies of relational matching, suggesting that fNIRS provides sufficient sensitivity to detect meaningful differences in relational complexity at the level of individual channels.

In contrast, ROI-level analysis for this contrast did not reveal any statistically significant effects after correction. This highlights a key limitation of spatial averaging: subtle differences between closely matched task conditions may be obscured, particularly when both involve high cognitive demands—as in the current paradigm, where 1st-order shape and color trials were intermixed within blocks, increasing task-switching requirements.

Together, the results for the general and specific task contrasts illustrate a critical trade-off between spatial specificity and statistical robustness across the two approaches to fNIRS analysis. Channel-level analysis excels at detecting localized, condition-sensitive activation, making it especially valuable for fine-grained contrasts. ROI-level analysis, by contrast, enhances sensitivity to broad activation patterns, as shown in the Task vs. Baseline contrast, but may fail to detect more subtle condition differences when spatial averaging reduces functional resolution. The choice between these approaches should therefore be guided by the contrast of interest and the expected spatial distribution of effects.

### 5.2 Effects of Number of Blocks on Task Activation

Here, we sought to determine how many task blocks are needed to obtain reliable behavioral and neural activation estimates in our paradigm, starting with an adult sample. Our goal was to balance statistical power with participant experience and experimental validity. To this end, we conducted blockwise convergence analysis across the full 18-block session, examining average beta values, variability, and standard error for the general and specific contrasts.

For the Task vs. Baseline contrast, beta values and standard error declined steadily with each additional set of the three task blocks, while the proportion of significant channels increased. These trends suggest improved signal-to-noise ratio with more data, although gains diminished after approximately 9 blocks. While extending the session beyond this point yielded marginal benefits, the most substantial improvements in model precision and group-level sensitivity for the general contrast were achieved within the first half of the experiment.

For 2nd vs. 1st-order trials, early blocks yielded inflated beta values and a high proportion of significant channels. Both overall activation strength and variability stabilized by around block 9, suggesting convergence on a more accurate representation of the neural processes underlying relational reasoning. The gradual decline in standard error across blocks further supports this improvement in estimation precision. Trends in behavioral performance across the session lends credence to the idea that this early peak reflects heightened cognitive demands during initial task-switching, followed by stabilization. Supplementary analyses comparing activation for the 1st and 2nd halves of the session provided complementary insights to the cumulative block number analyses, revealing substantial reorganization in the spatial distribution of significant channels—resulting in low correlations between 1st and 2nd halves—that may reflect cognitive shifts as a function of extensive practice. These observations underscore the point that more data isn’t always better: rather, it is important to identify a window during which brain activity is most representative of the cognitive processes of interest, after initial adaptation but before disengagement or overlearning. It may also be the case that the increased cognitive demands of the 1st-order trials in this paradigm may contribute to the difficulty in detecting stable neural differences between conditions. Consequently, extending the number of task blocks beyond 18 could provide sufficient data to improve the stability of the 2nd vs. 1st-order contrast. This interpretation is supported by the observed steady decline in standard error up to block 18, which likely continues with additional data, facilitating more precise and reliable estimates. However, these conclusions remain speculative, and further research is necessary to determine the optimal block count for this and similar paradigms.

This initial exploration aims to identify a methodological “sweet spot” for this paradigm in young adults, capturing stable behavioral performance, well-separated condition-specific neural signals, and improved group-level sensitivity without incurring the potential costs of longer sessions. Our approach also reinforces the value of combining amplitude, pattern- and variance-based reliability metrics, as well as the importance of visualizing spatial patterns, to fully capture the temporal dynamics of fNIRS signals in cognitive tasks. However, these analyses will need to be repeated in future with developmental and clinical samples, where sustained attention is limited and/or long sessions are not feasible. Our results provide a starting point for homing in on task duration for future study designs: across both contrasts, converging evidence from neural and behavioral data suggest diminishing returns after ∼9 blocks, marking a potential convergence point for reliable and interpretable contrast estimation. However, for finer-grained contrasts for which there may be more pronounced shifts in activation over time, we recommend piloting to empirically identify the optimal temporal window of initial practice on the task and subsequent data collection, assessing both signal reliability and behavioral stability within a session. Optimizing this window is likely crucial for interpretation of developmental change and differences between populations.

### 5.3 Functional Connectivity

To evaluate large-scale cortical integration during task performance, we compared functional connectivity analyses at both the channel and ROI levels. No significant differences in connectivity emerged between the 1st and 2nd-order conditions (consistent with unpublished results from our laboratory), suggesting that while relational complexity modulates localized activation patterns, the broader functional integration of the frontoparietal network remained stable across conditions. Accordingly, we collapsed across conditions for subsequent connectivity analyses. Significant connectivity was observed across multiple channels, but the strongest and most reliable pairwise connections were concentrated between LPFC channels and between PPC regions. Notably, left RLPFC channels exhibited the strongest pairwise connectivity, both with left VLPFC and with right RLPFC, consistent with its proposed role as a hub for executive control and relational reasoning.

The ROI-level analysis revealed a functionally coherent and anatomically interpretable frontoparietal network, with significant connectivity both within and across hemispheres. Robust frontal lobe coordination was observed between left RLPFC, DLPFC, and VLPFC, as well as between homologous regions in the right hemisphere. Bilateral frontoparietal integration was also prominent, including strong cross-hemispheric connections between RLPFC and inferior parietal cortex. Notably, left RLPFC again emerged as the most strongly connected ROI, consistent with findings from prior fMRI research involving this paradigm in a pediatric sample (Wendelken et al., J Neuro, 2017).

Crucially, ROI-level analyses produced consistent results across both the full sample (N = 73) and a particularly high-quality subsample (n = 36), indicating that this approach enables inclusion of more participants in functional connectivity analyses by averaging over variable channel-level quality within each region. This methodological strength preserved statistical power and enhanced generalizability—an important consideration for population-level research or clinical studies in which optode coverage may vary.

Together, these findings underscore the complementary strengths of each analytic approach. Channel-level analysis is highly sensitive to focal, optode-specific interactions and can reveal nuanced connectivity patterns that may be masked by data aggregation. Nevertheless, this approach is also more susceptible to local measurement noise and variability in optode positioning across participants, potentially leading to higher rates of data exclusion and reduced generalizability. Future research may benefit from combining both analytic levels, thereby leveraging their respective advantages while mitigating individual limitations.

### 5.4 Within Session Reliability of Task Activation

To evaluate the internal reliability of fNIRS task-based measures within a session, we compared activation and connectivity estimates across interleaved blocks, at both the channel and ROI levels. Interleaved designs offered the most direct assessment of measurement stability while minimizing confounds such as learning or fatigue, serving as a complementary validation of temporal consistency to the comparison of 1st and 2nd halves of the session (**Supplementary Section 9**).

The current findings provide a nuanced view of within-session stability in fNIRS-derived task activation, underscoring the importance of contrast type, spatial resolution, and region of interest. Task vs. Baseline activation demonstrated the highest within-session reliability across all analyses.

At the individual level, the Task vs. Baseline contrast showed moderate internal consistency at the channel level, with notable variability across individual channels and channel-by-subject interactions. LPFC channels exhibited significantly greater reliability than PPC channels, indicating that prefrontal regions may provide more robust signal estimates for broader task activation. This pattern was mirrored at the ROI level, which also showed moderate reliability with clear variability across regions. In contrast, the 2nd vs. 1st-order contrast again showed slightly lower internal consistency at both levels, with minimal variance attributable to specific channels or channel-by-subject interactions. LPFC and PPC regions did not differ significantly in this case, and channel-level ICCs were generally lower. These findings reinforce the conclusion that task complexity and cognitive contrast type play a critical role in determining the stability of measured activation.

At the group level, channel-level spatial patterns were highly consistent across interleaved comparisons, with strong correlation and variance preservation. In contrast, the 2nd vs. 1st-order contrast yielded low spatial stability at the group level. This suggests that this particular condition-specific contrast may not be suitable for investigating the spatial distribution of activation at either the channel or ROI level.

While our activation based internal consistency analyses yield similar results to previous research for both group and participant data at channel- and ROI-levels, our functional connectivity measures showed exceptional within-session reliability, with near perfect split-half correlations at the group level and across individuals. These findings far exceed the reliability levels reported in previous fNIRS studies using activation-based ICCs or overlap metrics. Our results suggest that connectivity-based approaches offer superior reproducibility and may be less affected by spatial noise or individual differences in activation topography.

Together, these findings highlight the critical role of spatial resolution and split-half methodology in assessing reliability. While ROI aggregation provides interpretable group-level summaries for broad contrasts like Task vs. Baseline, it tends to mask finer-grained variability that is captured more effectively at the channel level. Channel-wise analysis proved more sensitive to individual and condition-level effects, particularly for detecting subtle differences in relational reasoning contrasts.

Overall, these results support the value of multi-scale approaches to within-session reliability assessment in fNIRS. Our specific task contrast was highly unstable, whereas the general contrast of task activation versus baseline was more robust. Functional connectivity measures emerged as the most robust indicator of stable brain network organization, demonstrating exceptional consistency across both spatial scales and participant-level analyses. This suggests that connectivity-based metrics may offer a more reproducible alternative to activation-based approaches, especially for longitudinal designs and individual differences research.

### 5.5 Between-Session Stability of Task Activation and Functional Connectivity

To evaluate the suitability of fNIRS-derived measures for longitudinal and individual differences research, we assessed the inter-session stability of task-evoked activation and functional connectivity at both the channel and region-of-interest (ROI) levels. Sessions were spaced 14–33 days apart, offering a stringent test of short-term stability that provides a ceiling estimate for the reliability expected in developmental or intervention-based designs. By comparing these spatial scales across group- and subject-level analyses, we provide a comprehensive characterization of their test–retest reliability, including not only the consistency of activation patterns but also the stability of signal variance—an often-overlooked dimension critical to interpreting inter-individual variability and signal robustness.

#### 5.5.1 Task Activation Stability

These findings offer a detailed characterization of test–retest reliability in task-evoked fNIRS activation across sessions, highlighting key differences in stability as a function of spatial scale, contrast type, and anatomical region. At the participant level, between-session test–retest reliability was generally low, regardless of spatial scale or contrast. For the Task vs. Baseline contrast, reliability estimates were modest overall, though some individual channels—particularly in lateral prefrontal regions—showed acceptable-to-high levels of stability. ROI-level aggregation, while beneficial for group-level interpretability, did not enhance individual-level reliability and in some cases reduced sensitivity to consistent individual differences. For the 2nd vs. 1st-order contrast, reliability was again low across the board, with few channels demonstrating meaningful stability and parietal regions slightly outperforming frontal regions. These findings underscore the challenge of capturing stable individual-level activation patterns across sessions, particularly for complex contrasts.

At the group level, neural signal amplitude was consistently higher in session 1 than in session 2 across both contrasts and spatial scales, suggesting a potential decline in engagement or signal magnitude over time. Spatial activation patterns for the Task vs. Baseline contrast were reliably reproduced across sessions, particularly at the ROI level, highlighting the benefits of spatial averaging for enhancing reproducibility. In contrast, the 2nd vs. 1st-order contrast exhibited poor spatial reliability at both scales, indicating greater vulnerability to session-to-session variability. Variance-based analyses revealed a complementary pattern: signal variance was more stable for the 2nd vs. 1st-order contrast, whereas Task vs. Baseline variance was less consistent across sessions. This dissociation suggests that while spatial patterns of broad task engagement are stable, response magnitude may be more sensitive to task complexity and individual differences.

These results underscore the importance of aligning spatial scale with analytic goals when designing fNIRS studies. At the group level, ROI-based analyses demonstrated clear advantages for detecting stable spatial activation patterns, particularly for broad task contrasts like Task vs. Baseline, due to the statistical benefits of spatial averaging. ROI aggregation mitigates the influence of noisy or missing channels and enhances reproducibility, making it well-suited for studies prioritizing anatomical interpretability and robust detection of task-related activation across sessions. However, this benefit did not extend to condition-specific contrasts such as the 2nd vs. 1st-order comparison, which remained unstable across sessions even at the ROI level. Moreover, variance-based measures revealed that signal variability over time was better preserved for complex contrasts at the ROI level, despite poor spatial reproducibility.

In contrast, channel-level analyses, while offering finer spatial resolution, showed lower between-session stability on average, particularly at the group level and for condition-specific contrasts. Nevertheless, some individual channels, especially in prefrontal regions, showed acceptable or good reliability, and channel-wise metrics preserved variability in signal amplitude that may be obscured by regional averaging. This suggests that channel-level data may be more sensitive for capturing participant-specific features and should be considered in studies focused on individual differences, longitudinal change, or personalized applications such as neurofeedback or intervention tracking.

Together, these findings support a complementary analytic strategy: using ROI-level aggregation to enhance group-level robustness and interpretability, while retaining channel-level analyses to preserve sensitivity to individual variability and spatially localized effects.

#### 5.5.2 Functional Connectivity Stability

In contrast to activation analyses, functional connectivity measures—both at the channel and ROI level—demonstrated exceptionally high test–retest reliability, especially at the group level. Channel-level connectivity matrices yielded near-perfect correlations across sessions for all conditions, indicating the global architecture of functional networks is highly preserved over time, even at a fine-grained spatial resolution. ROI-based matrices showed slightly higher correlations, suggesting a marginal advantage in signal stability, likely due to enhanced signal-to-noise ratio via anatomical aggregation. Importantly, these patterns held across task conditions and in the Task vs.Baseline comparison, indicating that core network topology remains stable across both cognitive state and time.

At the individual level, both spatial scales again produced strong test–retest reliability. Channel-wise connectivity correlations were high for all participants across conditions, with all participants exhibiting significant stability. ROI-level correlations were even stronger with 94% of participants showing correlations exceeding r = .80. This pattern suggests that while channel-level functional connectivity captures individualized neural signatures, ROI-level connectivity provides a more robust and generalizable index of inter-session reliability, potentially making it the preferred approach for clinical, developmental, or biomarker research where individual tracking is critical.

Importantly, because connectivity was estimated using pairwise correlations across full trial epochs, variance measures were not computed separately. However, the consistently strong test–retest correlations at both scales implicitly reflect stability in both connectivity patterns and underlying signal structure, supporting their use as reliable longitudinal markers.

The current study demonstrates higher between-session reliability than previously reported in fNIRS test–retest studies. For example, Yeung et al. (2023) reported only moderate ICCs (0.37–0.62) for task-evoked frontal activation during a working memory task, with limited individual-level reliability for differential contrasts. Similarly, Nittel et al. (2025) found replicable group-level patterns in working memory and inhibitory control tasks but low individual ICCs (< 0.50) and inconsistent overlap metrics. de Rond et al. (2023) showed that activation reliability in motor cortex declined sharply when the fNIRS cap was removed and refitted, underscoring the impact of optode placement. Xu et al. (2023) demonstrated strong resting-state fNIRS test–retest reliability for graph metrics in stroke patients, particularly when scans were sufficiently long and focused on low-frequency bands.

Our findings extend this literature by incorporating both t-value and signal variance correlations at the group level, as well as ICC and variance attributed to both channel and ROI at the participant level. While the participant level data showed similar results to previous work (ICCs ranging from .00 - .65 for Task vs. Baseline and .00 - .45 for 2nd vs. 1st-order), the group-level ROI-based metrics in our study yielded stronger reliability than those reported previously (e.g., r = .87 and variance = .93 for Task vs. Baseline), while functional connectivity far outperformed typical activation-based estimates (channel-level r > .97, ROI-level r > .98). It is very likely that participant level ICCs for both spatial scales were significantly impacted by the overall poorer data quality in parietal channels compared to PFC.

These results provide a robust foundation for future longitudinal research, highlighting the strengths of combining both ROI and channel-level approaches to capture reproducible, interpretable, and spatially sensitive neural measures. It is important to note that these results reflect only short-term stability (2–4 weeks), which is appropriate in the context of developmental populations where neural and behavioral measures may fluctuate over time. This short-term window provides a practical upper bound, or “ceiling,” on the maximum stability that can be expected.

### 5.6 Within- and Between Session Comparison

A direct comparison of within- and between-session reliability highlights several important trends regarding the robustness of fNIRS-derived activation measures. Across both temporal scales, the Task vs. Baseline contrast demonstrated consistently higher reliability than the 2nd vs. 1st-order contrast, supporting prior findings that condition-specific contrasts—particularly those calculated as difference scores (e.g., Condition 1 - Condition 2)—result in a high degree of error making them inherently less stable and more vulnerable to cognitive variability and noise (Heilicher et al., 2022). This instability may be further exacerbated by practice or strategy-related effects across sessions, which can alter the strength of differential contrasts such as 2nd vs. 1st-order comparisons.

At the channel level, both within- and between-session results revealed high variability in reliability across channels, yet a subset of channels—particularly those located in lateral prefrontal cortex (LPFC)—showed moderate to high reliability in both contexts. These findings suggest that while channel-level data can be noisy and sensitive to signal quality fluctuations, certain individual channels can nonetheless provide robust and reproducible indices of activation, particularly in well-covered and low-noise regions such as the PFC. In contrast, parietal channels consistently showed lower reliability across both within- and between-session assessments, likely due to poorer signal quality and higher rates of noisy or borderline-usable channels.

At the ROI level, overall reliability was lower than at the channel level for individual subjects, especially for the 2nd vs. 1st-order contrast. However, some regions, namely bilateral rostrolateral prefrontal cortex (RLPFC), demonstrated relatively greater stability across sessions. Importantly, the direction and magnitude of reliability estimates for specific ROIs sometimes differed between within- and between-session comparisons, suggesting that aggregation may obscure meaningful subject-level variance or amplify inconsistencies across time. This highlights the need to interpret ROI-based results with caution, particularly when assessing individual-level effects in longitudinal designs.

Functional connectivity analyses further revealed exceptional stability across both within- and between-session assessments. At the individual level, channel-wise connectivity showed robust reproducibility across participants, and ROI-level measures yielded even higher consistency, with the majority of participants exceeding reliability thresholds commonly required for clinical or biomarker research. At the group level, split-half correlations for connectivity matrices were near perfect across all conditions and both spatial scales, far surpassing traditional activation-based reliability metrics. These findings indicate that the global structure of functional brain networks is highly preserved across time and cognitive states. These results suggest that functional connectivity, especially when derived from ROI-level aggregation, provides a highly stable and generalizable neural signature that is well-suited for longitudinal, developmental, and individualized research applications.

Together, these results emphasize the complementary value of multi-scale analysis in assessing the stability of task-evoked activation. Channel-level analyses may be more vulnerable to variability, but they still capture reliable subject-specific patterns, especially in the PFC. As a result, they are particularly useful for research focused on individual differences. ROI-level analyses, though more robust at the group level, may underperform in capturing reliable participant-level effects, especially for condition-specific contrasts. These findings underscore the importance of aligning spatial scale, contrast type, and analytic goals when interpreting fNIRS data in both cross-sectional and longitudinal research contexts.

These findings offer a cautious but optimistic assessment of fNIRS as a tool for longitudinal and developmental research. At the group level, ROI-based activation analyses of broad contrasts such as Task vs. Baseline yielded high spatial stability (r > .85), while functional connectivity measures demonstrated near-ceiling reproducibility (r > .95). At the individual level, task activation ICCs were modest at best, often falling below the thresholds required for individual-level inference. However, select prefrontal channels showed acceptable reliability, and individual functional connectivity profiles remained stable across sessions at both the channel and ROI levels. Overall, the most stable data emerged from ROI-level aggregation at the group level and from selective high-quality channels or connectivity matrices at the individual level. A hybrid approach—leveraging ROI-level metrics for group robustness and interpretability, and channel-level or connectivity-based metrics for individual variability—appears best suited for fNIRS applications in clinical, developmental, and longitudinal research contexts.

### 5.7 Impact of Hair and Skin Characteristics

Our findings provide further evidence that individual differences in hair color and skin tone meaningfully impact fNIRS signal quality, underscoring the critical need to incorporate phenotypic characteristics into methodological reporting and study design (cf. Bradford et al., 2024). Specifically, darker skin tones (more melanin) and darker hair colors, particularly black and red (more eumelanin or pheomelanin, respectively), were associated with a significantly greater number of bad channels identified during preprocessing. These effects were observed independently, without significant moderation by hair texture or type, suggesting that melanin-related optical properties and hair pigmentation remain primary sources of signal degradation in near-infrared spectroscopy.

Although not all pairwise comparisons reached conventional significance thresholds, the overall pattern aligns with previous studies demonstrating reduced optode-scalp coupling and increased signal attenuation in participants with darker pigmentation or denser hair coverage (Holmes et al., 2024; Simmons et al., 2025; Yücel et al., 2024). The present findings reinforce the argument that hair and skin phenotypes should not be treated as nuisance variables or omitted from consideration, but rather as legitimate sources of biological variance that can systematically bias data quality.

These results call for standardized inclusion of skin tone and hair phenotype metrics in all future fNIRS reports, particularly in the context of sample description, quality control procedures, and exclusion criteria. Without such reporting, the generalizability of neuroimaging findings remains limited, particularly in diverse populations.

Moving forward, it is essential for the field to develop and test targeted mitigation strategies to address these phenotypic impedances. This includes engineering improvements in optode design, better cap configurations for dense or curly hair, and novel preprocessing techniques that explicitly model or correct for signal loss due to pigmentation. Crucially, these solutions must be validated across a broad spectrum of hair and skin types to ensure equity in data acquisition and interpretation. By treating phenotypic signal barriers not as artifacts but as solvable design and analytic challenges, the field of fNIRS can move toward more inclusive and scientifically rigorous practices.

## 6. Conclusion

This study demonstrates that task-based fNIRS reliably engages frontoparietal networks, producing effect sizes comparable to prior fMRI studies of reasoning. ROI-level analyses enhanced sensitivity to broad activation, while channel-level analyses captured finer spatial detail, particularly for condition-specific contrasts. Approximately nine blocks of data optimized the balance between signal stability and participant burden, though complex contrasts may require additional data. Functional connectivity emerged as a highly reliable metric across sessions and spatial scales, outperforming traditional activation measures and underscoring its utility for longitudinal and individualized research.

Group-level analyses revealed greater between-session stability for broad ROI-level contrasts. However, reliability at the individual level was more limited, with only select prefrontal channels and connectivity metrics showing potential. These findings highlight the value of combining ROI-and channel-level approaches and incorporating multiple reliability metrics, including amplitude-, pattern-, and variance-based measures, to capture the full temporal and spatial dynamics of fNIRS signals.

Importantly, signal quality was lower in parietal regions, contributing to reduced reliability and emphasizing the methodological challenge of maintaining consistent coverage across the scalp. The observed low reliability in parietal channels likely reflects these signal quality issues, which should be considered when interpreting results. Additionally, while the task was effective in eliciting reasoning-related activation, it may have lacked the contrast strength necessary to produce highly stable condition-specific effects. Future work should extend this framework to pediatric populations, where signal quality and participant variability pose greater challenges.

Finally, we emphasize the impact of hair and skin phenotypes on fNIRS signal quality. Darker pigmentation and hair were associated with greater signal loss, reinforcing the need for standardized phenotypic reporting and methodological innovations to improve data equity. Together, these findings support fNIRS as a promising tool for cognitive neuroscience, while identifying critical areas for refinement to enhance its reliability, inclusivity, and generalizability across diverse populations and study designs.

### Limitations and Future Directions

Several limitations should be acknowledged when interpreting the current findings. As anticipated, signal quality was lower in parietal regions, which likely contributed to reduced reliability in these areas and underscores persistent challenges in achieving consistent coverage across the scalp. Additionally, the task manipulation employed, while well-validated for eliciting relational reasoning, may have been too subtle to generate highly reliable activation patterns, particularly for complex contrasts such as 2nd vs. 1st-order conditions. These results are based on a single experimental paradigm and sample of healthy young adults; future work should extend this reliability framework to pediatric populations, where greater inter-subject variability and lower signal quality may be expected. To ensure preproducible results in such contexts, it is especially important to focus on the most robust analytic metrics, such as ROI-level functional connectivity. A critical avenue for future research involves addressing the impact of phenotypic variation on signal quality, including the well-documented impacts of darker or more textured hair and higher melanin content in skin. Although recent advances in hardware design have aimed to mitigate these disparities, systematic evaluation of their effectiveness is still urgently needed to ensure fNIRS can serve as an equitable and reliable tool in diverse populations.

## Supplementary Materials

### S1. Participant Demographics

**Supplementary Table 1.**
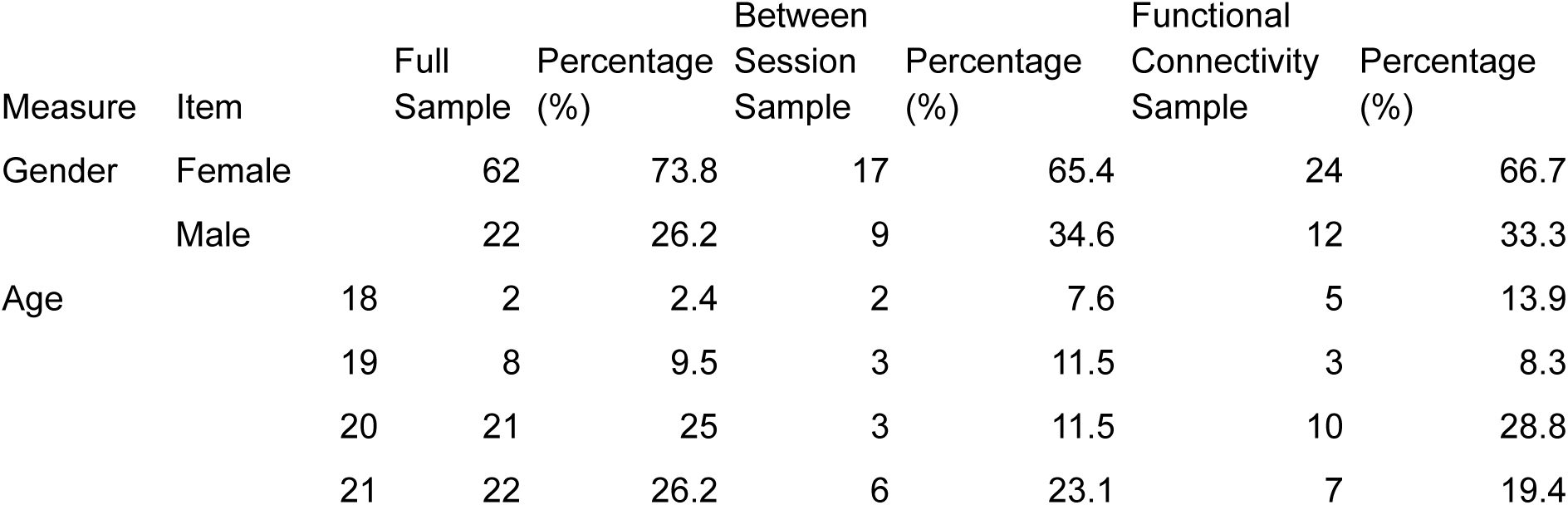
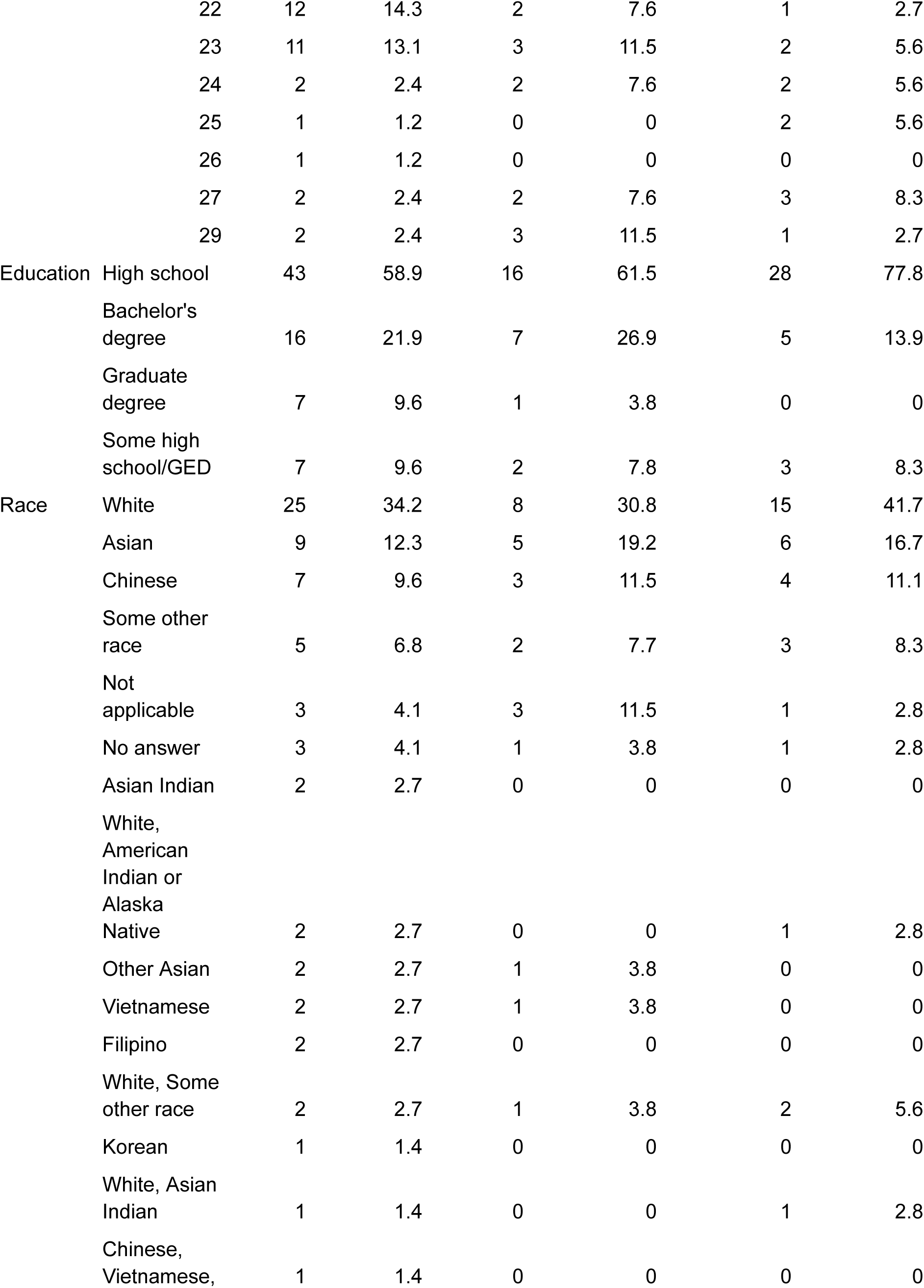
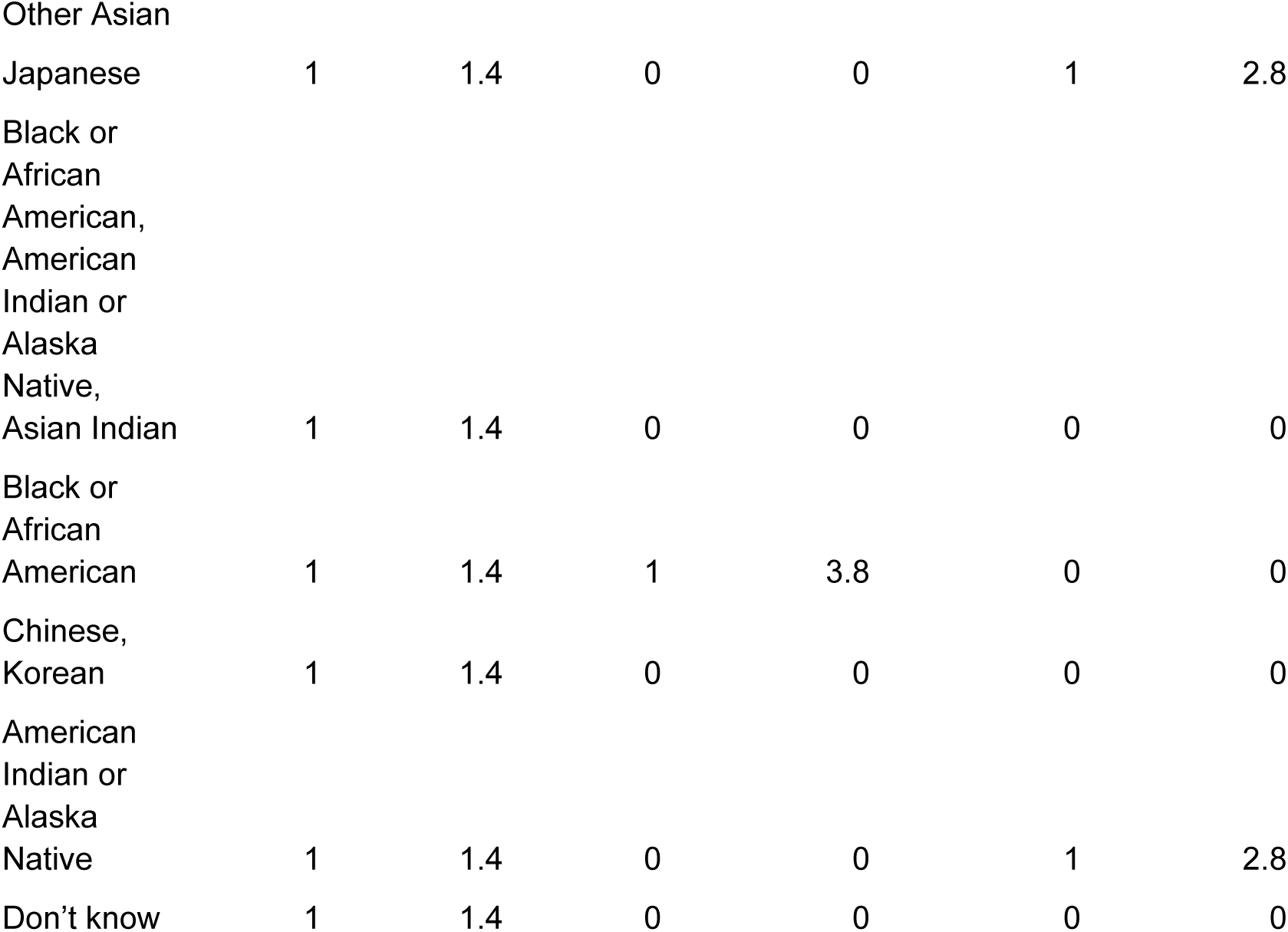
Demographic information for all samples.

### S2. Channel Information

**Supplementary Table 2.**
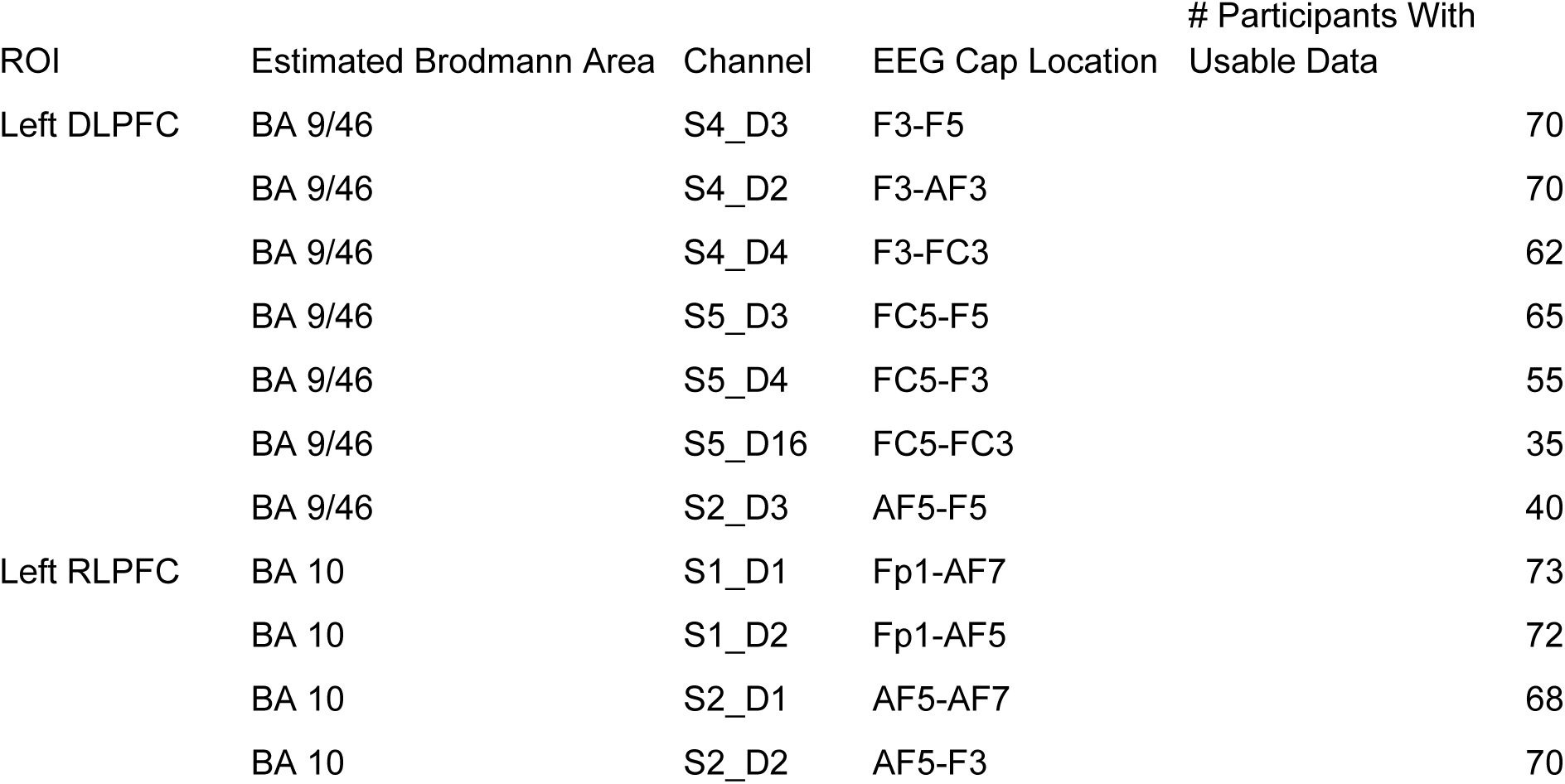
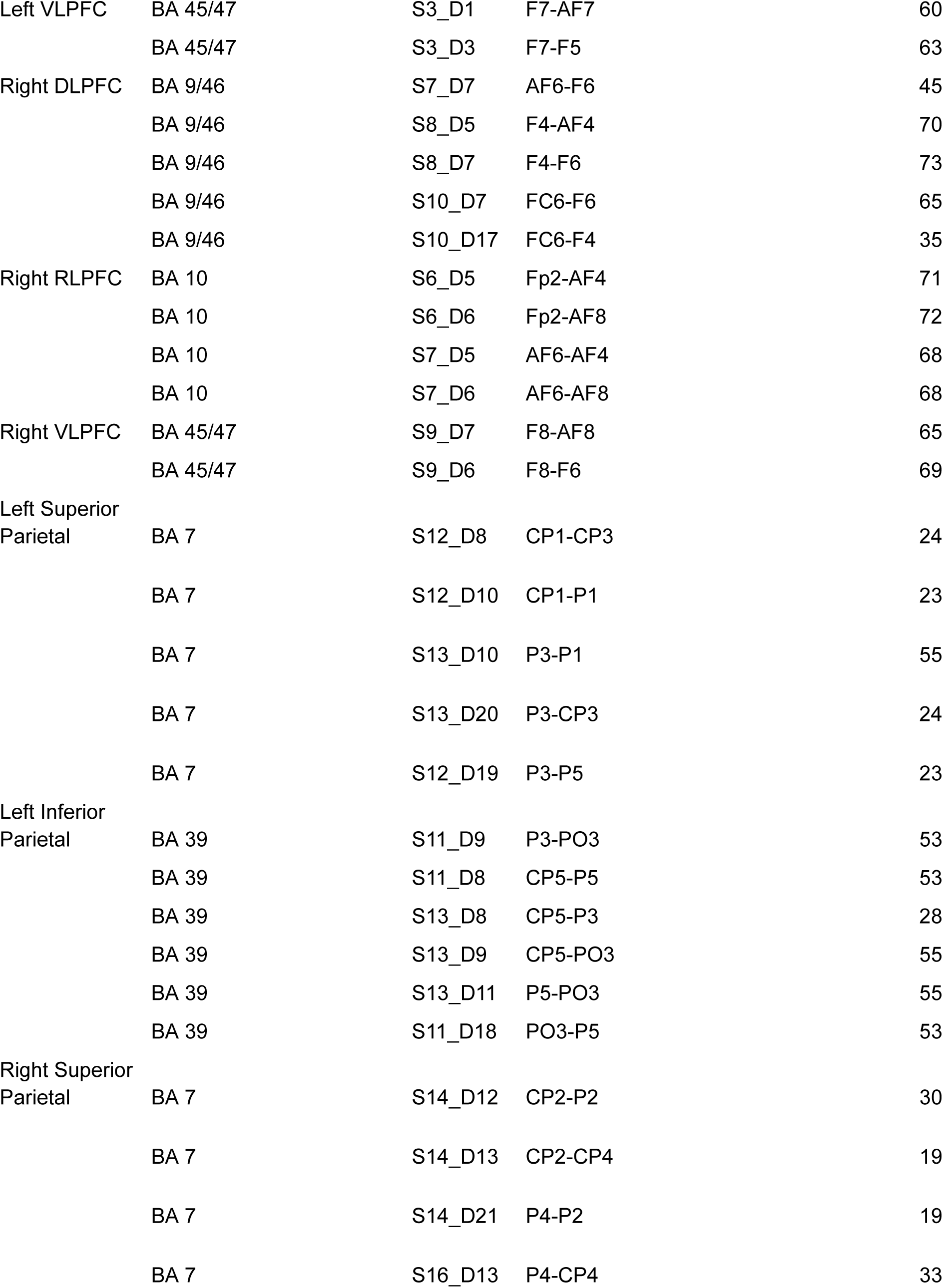
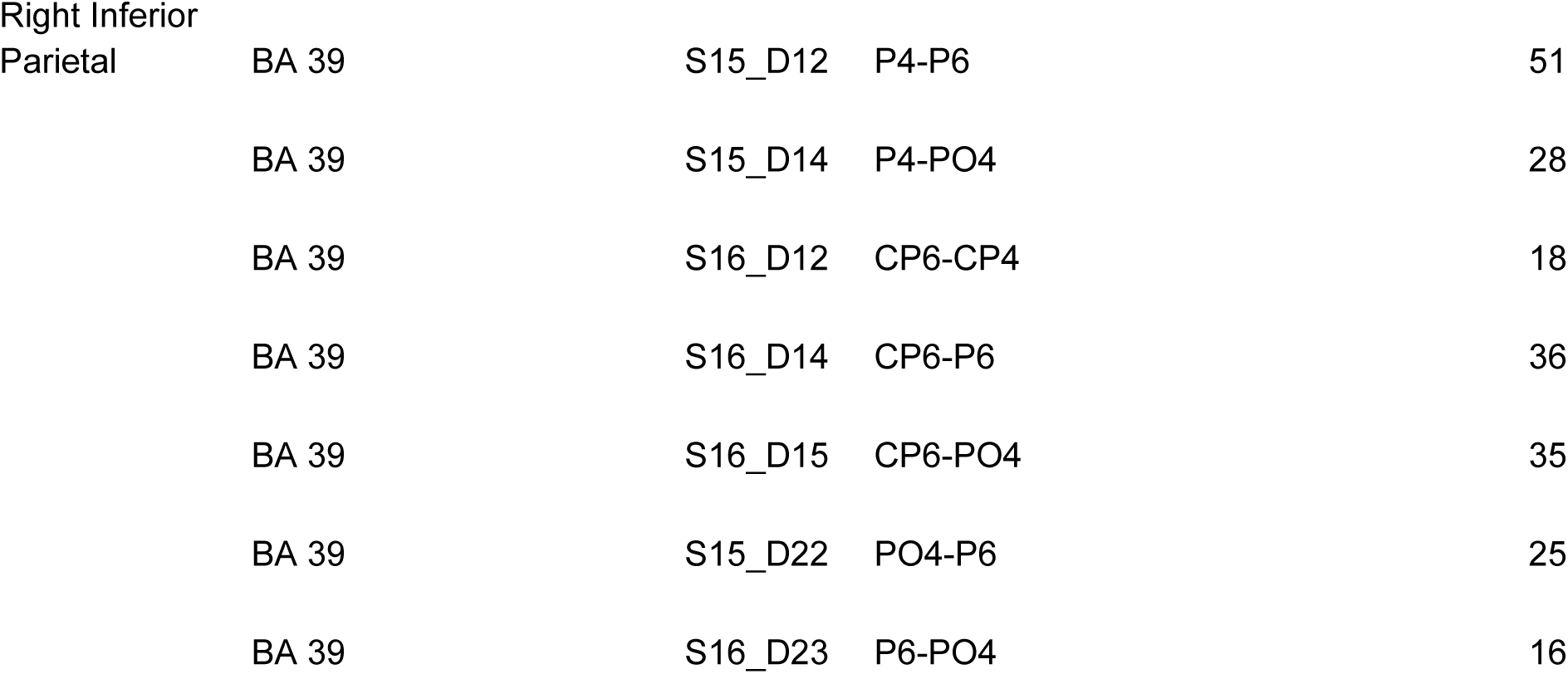
Channels with corresponding EEG cap locations, approximate Broadman area locations, and number of good participants for each channel.

### S3. Channel Signal Quality

**Supplementary Figure 1.**
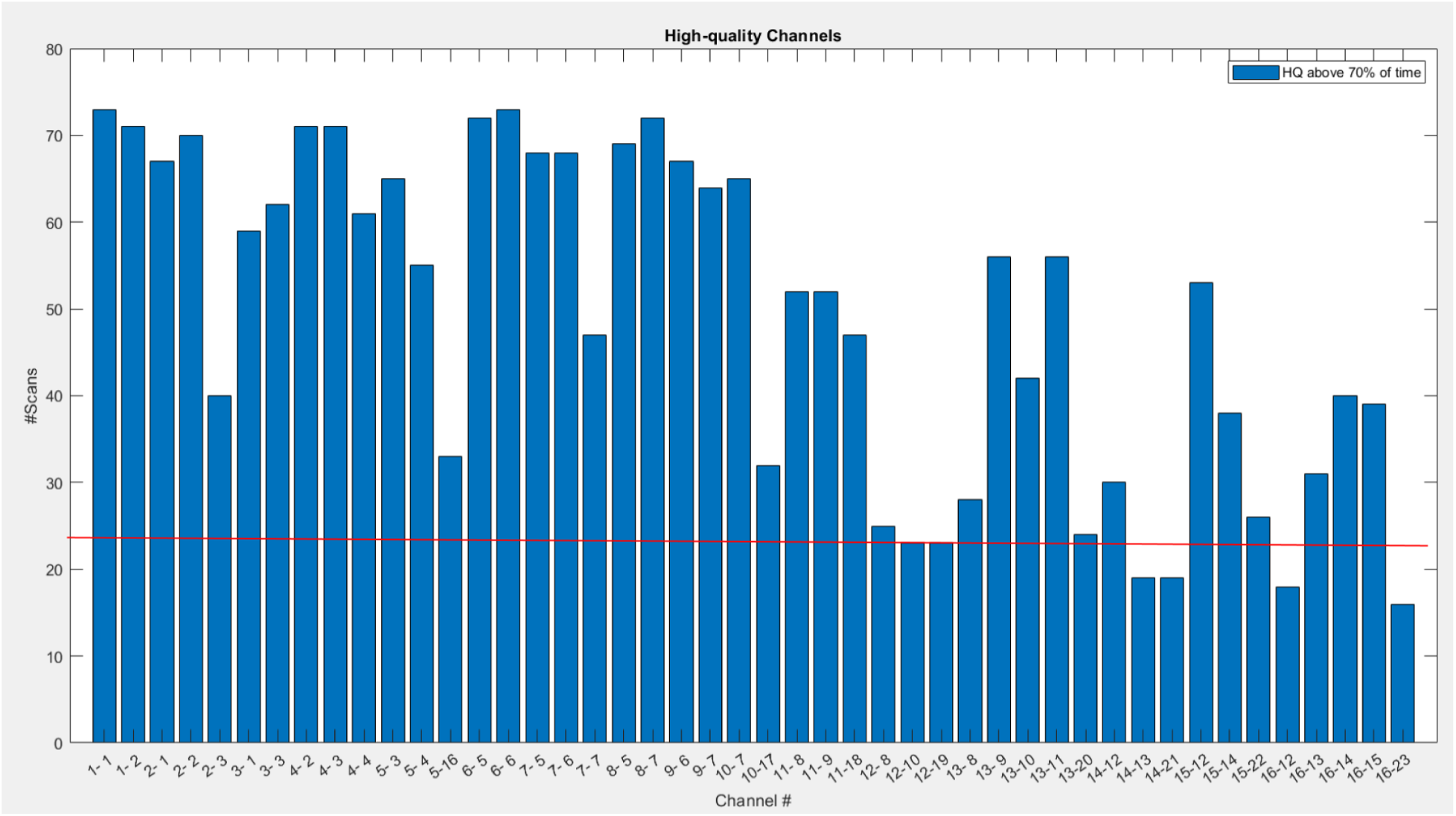
Channel level plot of the number of participants with good quality data per channel. The red line represents our exclusion criteria of a minimum of 23 good channels to be included in the following analyses. The 4 channels that did not meet the minimum criteria were excluded.

**Supplementary Figure 2.**
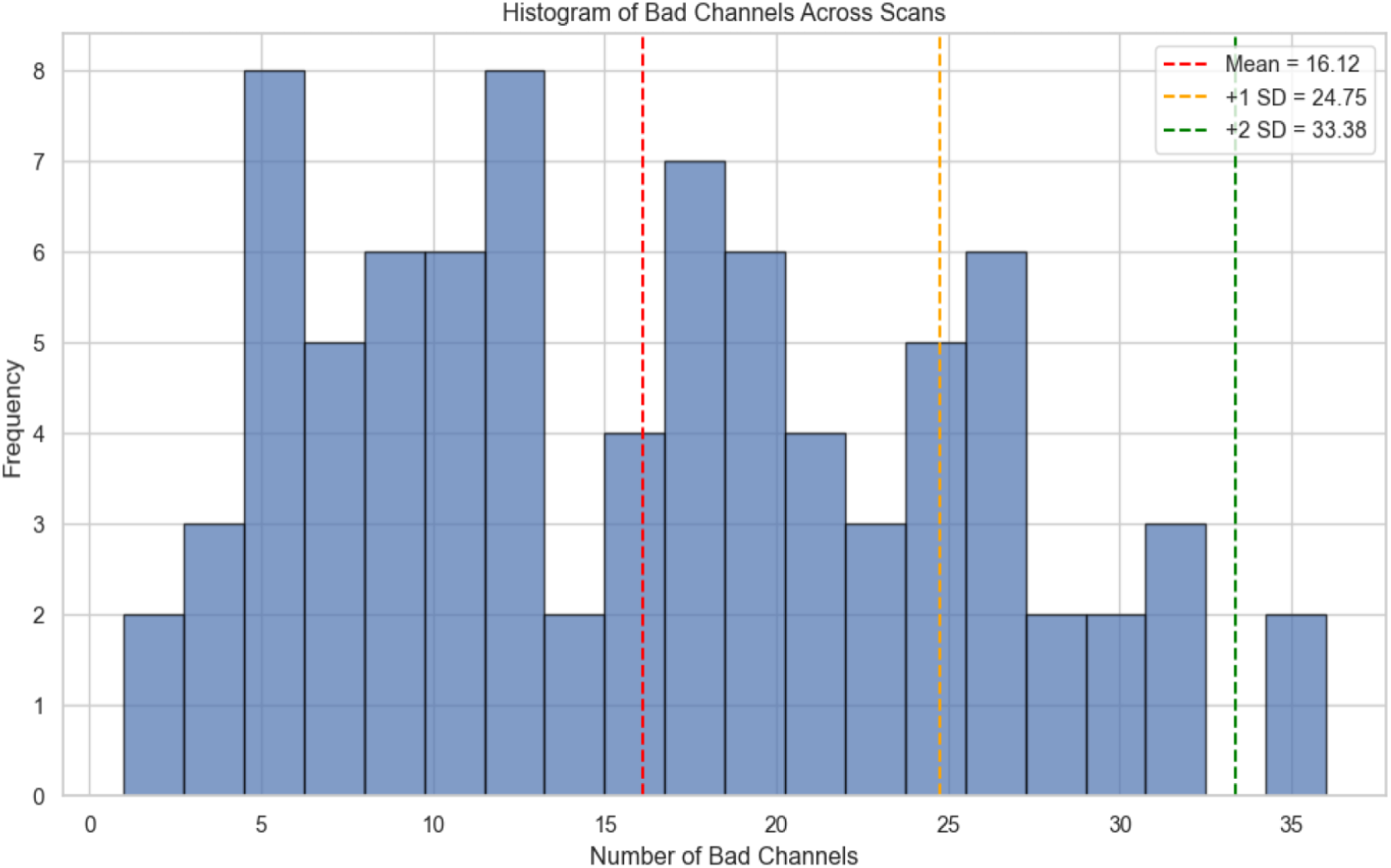
Distribution of number of bad channels for full sample prior to exclusion (N = 84). Red dashed line is the mean of all bad channels across participants, gold line is 1 standard deviation above mean. Exclusion of participants based on the rounded value of 1 SD above the mean of bad channels (>27). If participants exceeded this threshold of 27 bad channels they were removed entirely from the analysis.

### S4. Behavioral Data

**Supplementary Figure 3.**
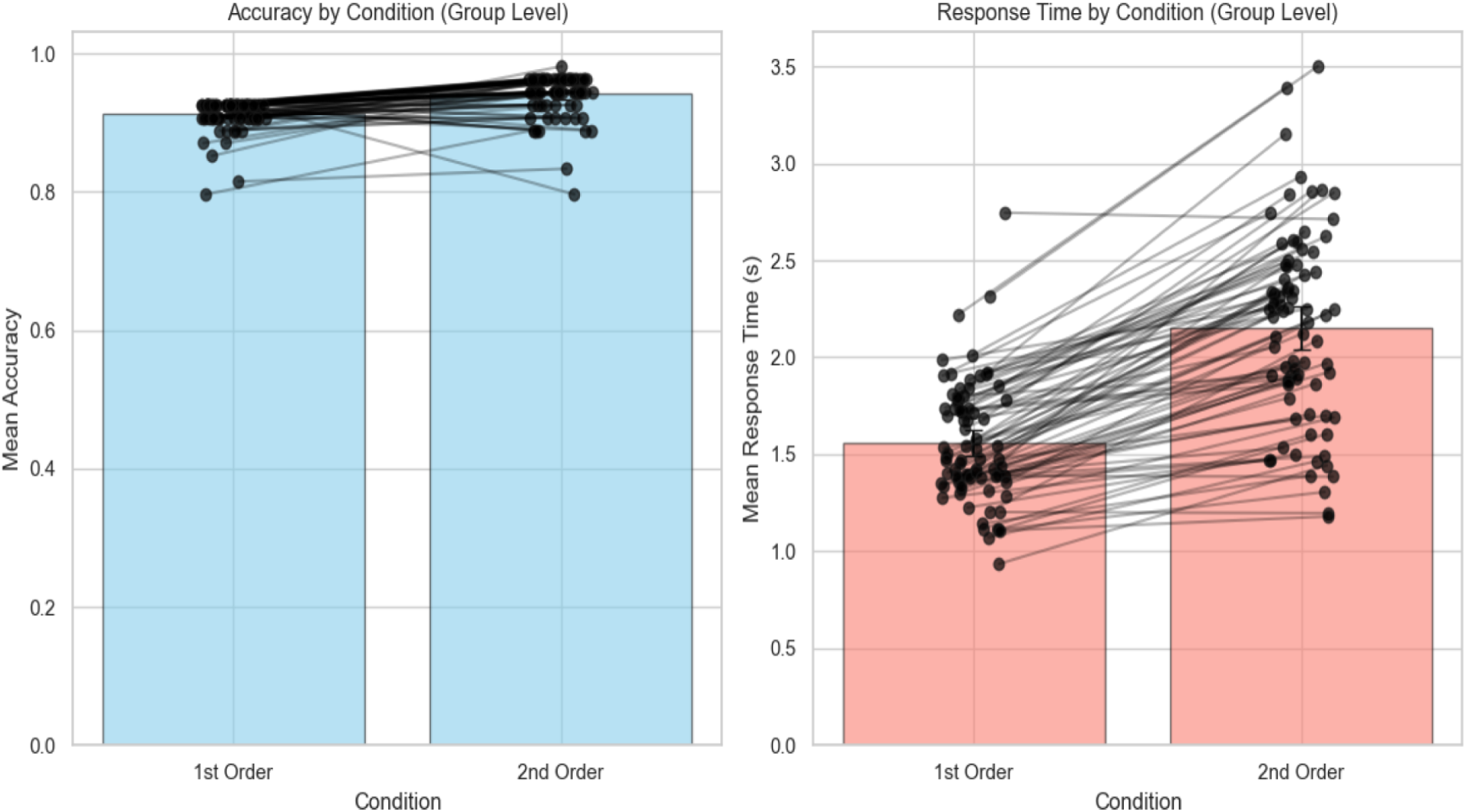
Group level accuracy and response time by condition.

### S5. Task Activation for Individual Conditions

#### 1st-order vs. Baseline contrast

During 1st-order trials (Shape and Color conditions) relative to baseline, significant group-level activation (q < 0.05) was observed in 15 channels across bilateral frontal and parietal regions. These included channels positioned in such a way as to target left and right dorsolateral prefrontal cortex (DLPFC), left rostrolateral prefrontal cortex (RLPFC), bilateral inferior parietal cortex, and bilateral superior parietal cortex. The activation pattern indicates engagement of frontoparietal regions even during lower-order relational judgments.

#### 2nd-order vs. Baseline contrast

For the 2nd-order (Match) condition, which required integration across multiple dimensions, we identified 14 significantly active channels (q < 0.05). Similar to Condition 1, activation was found bilaterally in DLPFC, RLPFC, VLPFC and Parietal Cortex, with additional engagement of more anterior prefrontal regions, compared to first order.

**Supplementary Figure 4.**
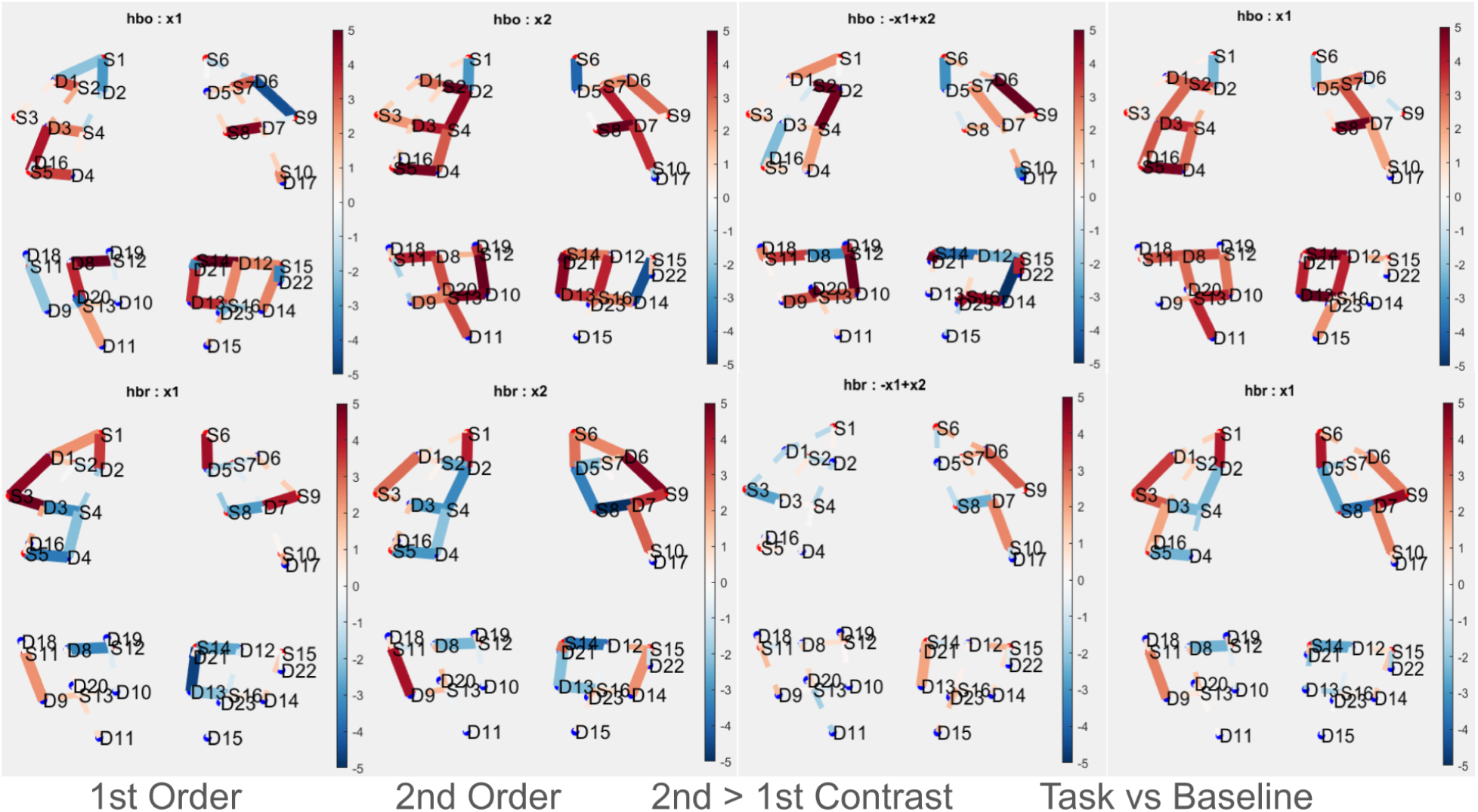
Condition level HbO (top row) and HbR (bottom row) results. Indicates the relationship between HbO and HbR are not perfectly negatively correlated and there is some discrepancy between the two signals.

### S6. Changes in Performance Across Blocks

**Supplementary Figure 5.**
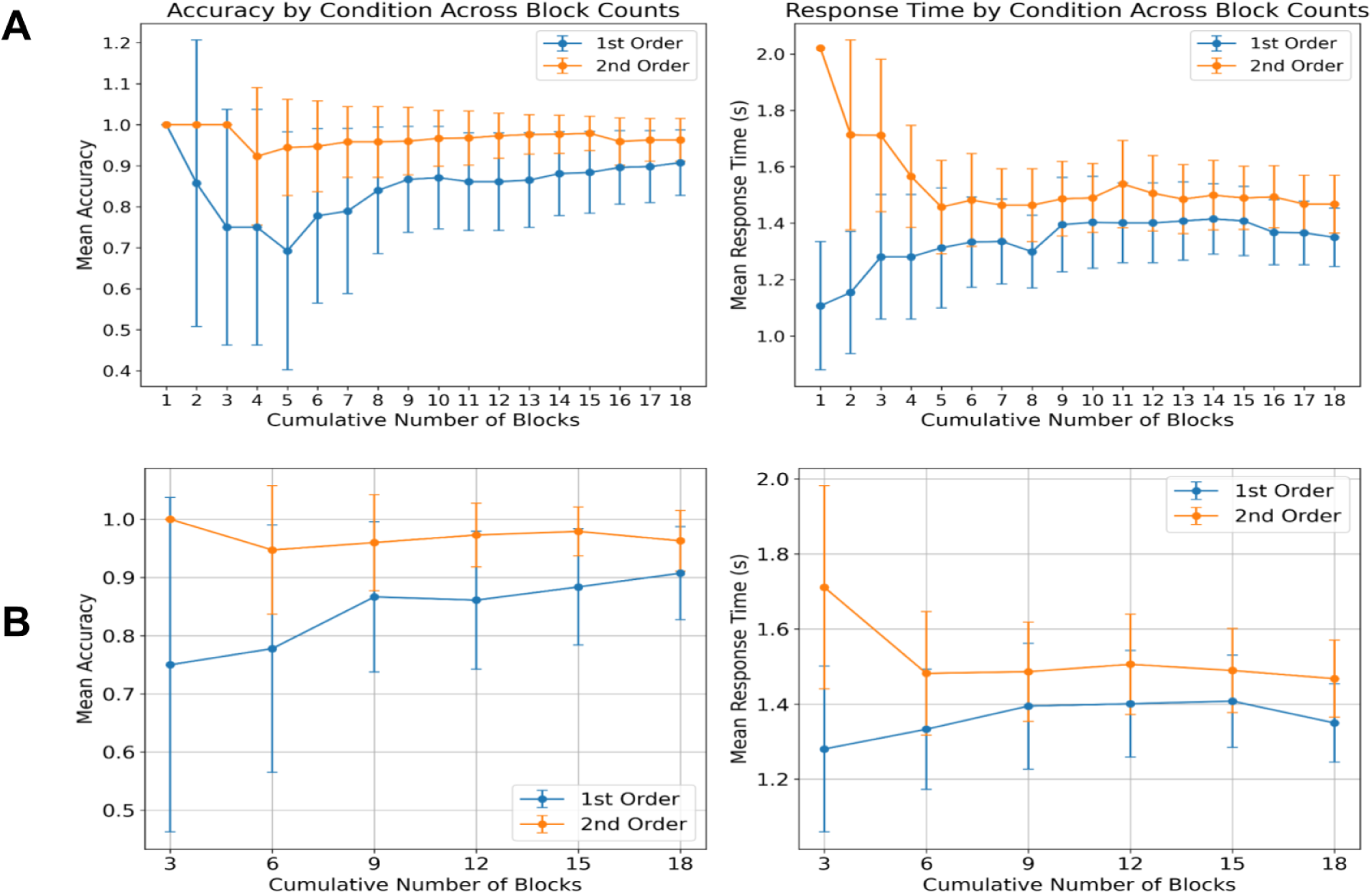
Mean conditional level accuracy and response time as a function of number of blocks. Block order was completely random for each participant. **A**. All 18 blocks **B**. Block intervals of 3

### S7. Functional Connectivity at the ROI level

**Supplementary Figure 6.**
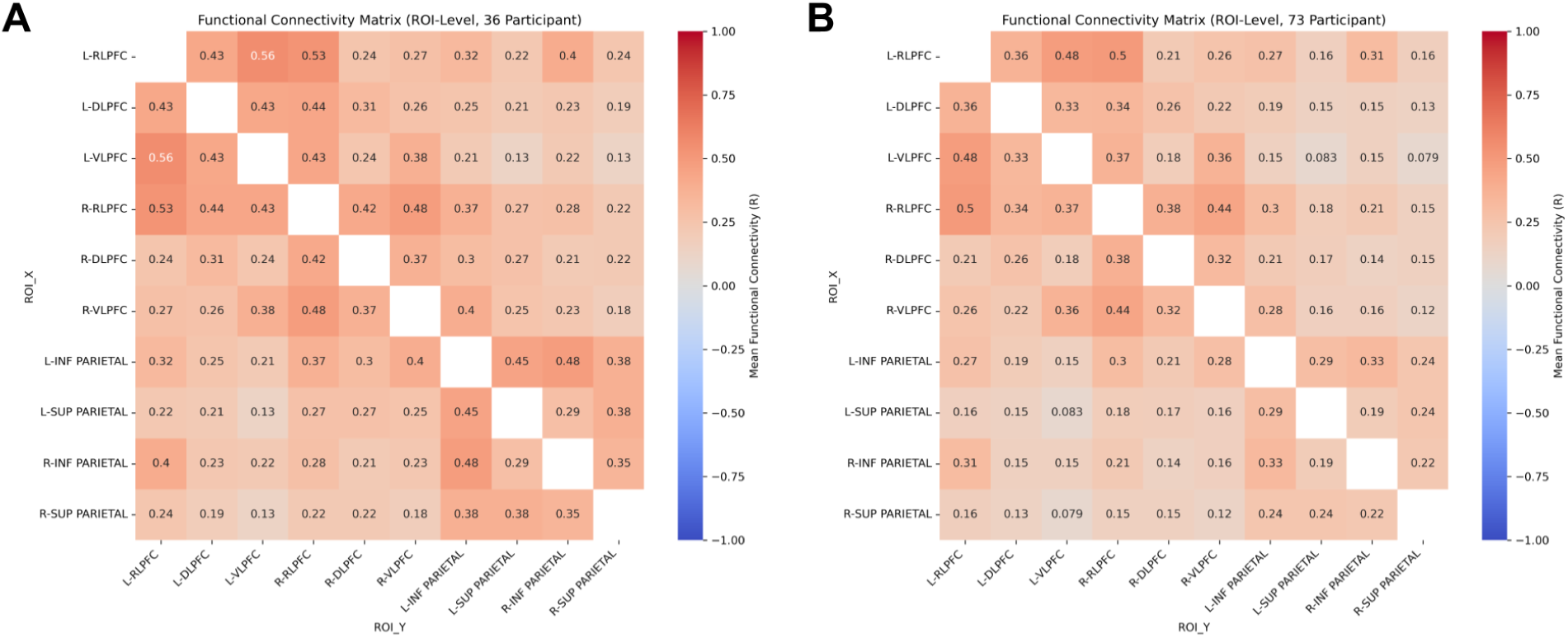
Functional Connectivity correlation matrix at the ROI level. **A**. Subsample of 36 participants. **B.** Full sample of 73 participants.

### S8. Channel Significance Status Across Blocks

**Supplementary Figure 7.**
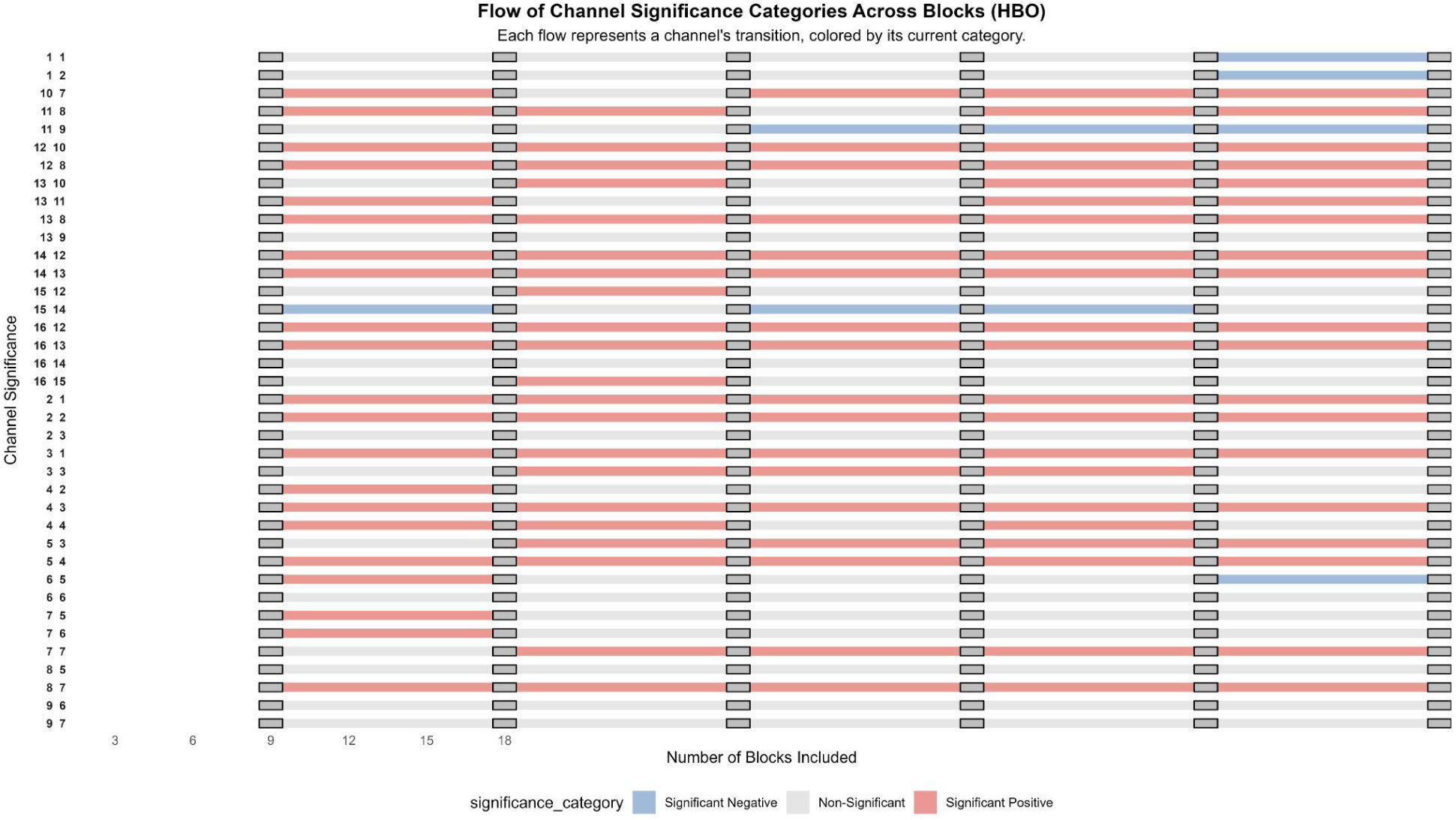
Channel level data on the significance and sign of activation as a function of the number of blocks included for the Task vs. Baseline condition.

**Supplementary Figure 8.**
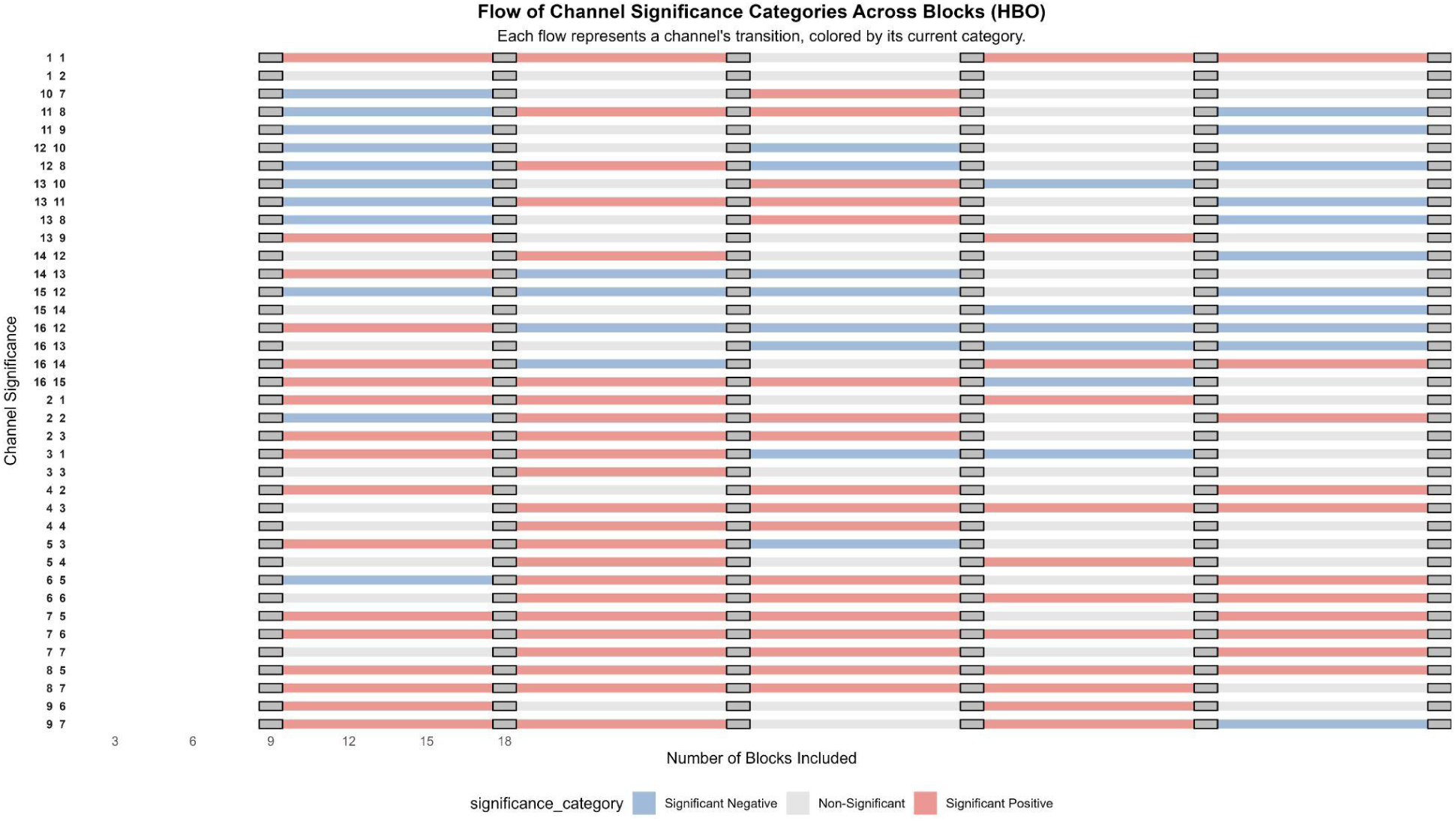
Channel level data on the significance and sign of activation as a function of the number of blocks included for the 2nd vs. 1st-order condition.

### S9. Comparison of Task Activation Across 1st and 2nd Halves of Session

In addition to conducting split-half analyses across interleaved blocks to assess internal consistency reliability, we conducted parallel analyses comparing the 1st and 2nd halves of the dataset for all 73 participants who completed one testing session. In this way, we assess the degree to which activation changed over time, complementing the analyses examining the effect of cumulative block number.

#### Group-Level Analyses

At the channel level, Task vs.Baseline showed high stability across the 1st and 2nd halves (r = .65, ρ = .79, variance = .90, p < .001 across 36 channels), though this dropped substantially when limited to significant channels (r = .20, ρ = .33, variance = .27, p = .71). By contrast, the 2nd vs. 1st-order contrast showed low stability (r = –.25, ρ = –.66, variance = .35, p = .14), but with higher variance stability, indicating more consistency in signal magnitude than in activation patterns. Filtering to significant channels in this comparison also showed low stability (r = –.04, ρ = –.09, variance = .32, p = .90). At the ROI level, the Task vs. Baseline contrast again demonstrated strong internal consistency for the 1st vs. 2nd half comparison (r = .60, ρ = .74, *p* = .07, variance = .43). In contrast, the 2nd vs. 1st-order contrast yielded low and nonsignificant reliability for 1st vs. 2nd half blocks (r = –.46, ρ = .63, *p* = .22, variance = .47). Overall, both channel- and ROI-level stability for Task vs.Baseline was consistently high while the stability for both Task vs.Baseline and 2nd vs.1st-order contrasts remained low across spatial scales.

In sum, topographic visualizations of group-level beta maps revealed that the set of channels showing significant activation in the first half of the experiment often differed from those in the second half. While the task continued to elicit strong hemodynamic responses, the cortical regions most engaged changed over time, resulting in low split-half correlations. This pattern is likely to be explained by practice effects. As participants progressed through the session, their cognitive strategies may have shifted, enabling them to perform the task with greater automaticity or efficiency. Given that accuracy did not decline in the second session, this reorganization of the neural signal may reflect a change in strategy and/or neural adaptation across trials.

**Supplementary Figure 9.**
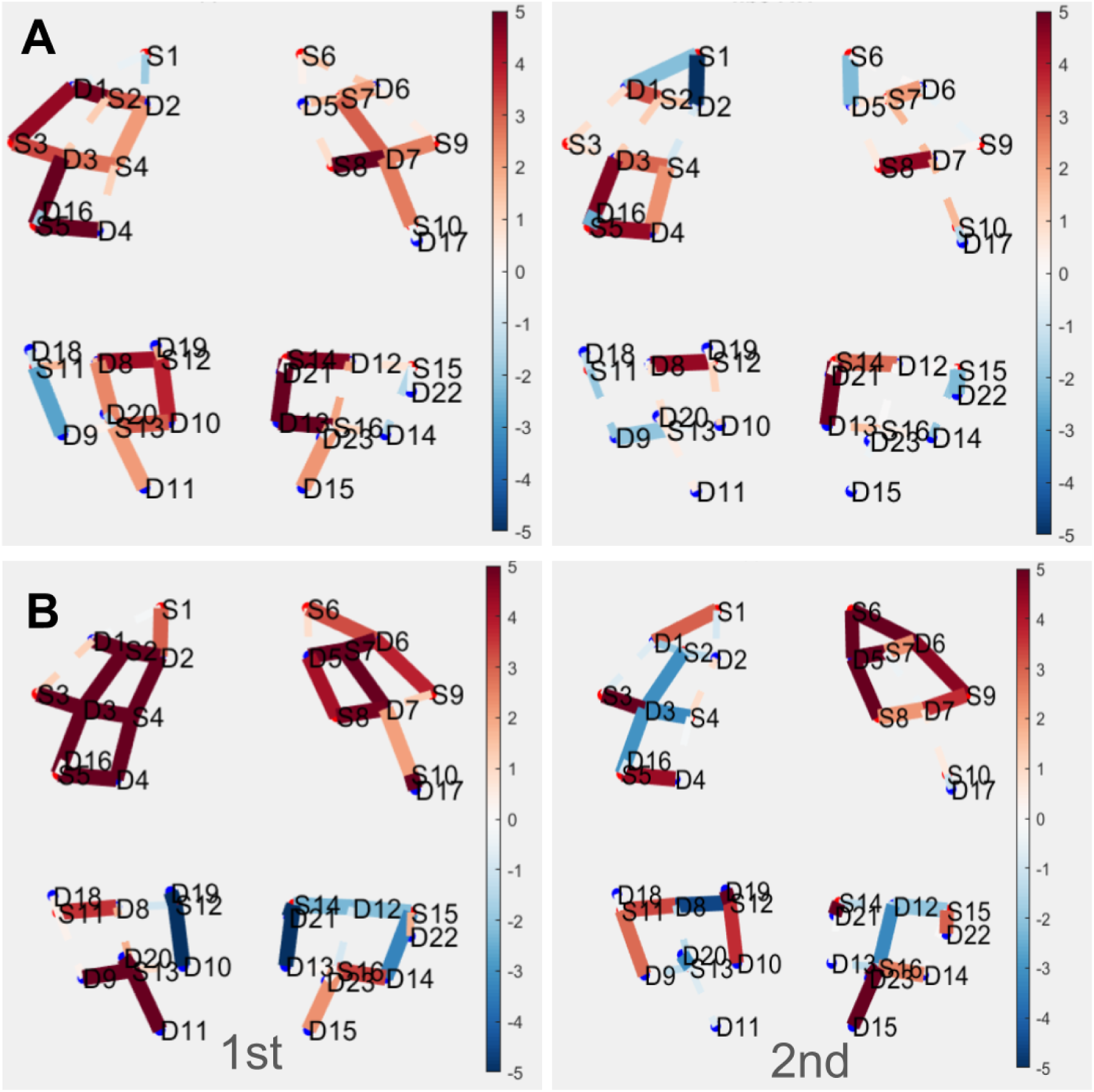
Comparisons of 1st and 2nd half. **A**.1st half (left) 2nd half (right) Group level activation plots for the channel level Task vs.Baseline contrast. **B.** 1st half (left) 2nd half (right) Group level activation plots for the channel level 2nd vs.1st contrast.

#### Participant-Level Analyses

### S10. Within-Session Stability

#### S10.1 Group Channel-level Task Activation Reliability for Interleaved Split-half Analyses

At the channel level, the Task vs. Baseline group contrast showed high within-session spatial stability for interleaved blocks (r = .60, ρ = .75, variance = .97 p < .001). However, when filtering for significant channels, stability became non-significant (r = .49 ρ = .66, variance = .08, p = .40). For the 2nd vs. 1st-order contrast, interleaved blocks showed low spatial stability (r = .06, ρ =.12, variance = .44, p = .71); this was also true when including only channels with significant activation in both halves (r = .04, ρ = .09, variance = .63, p = .89). At the ROI level, Task vs.Baseline demonstrated near perfect internal consistency while the 2nd vs.1st-order yielded zero reliability. Overall, both channel- and ROI-level analyses revealed strong within-session spatial stability for Task vs. Baseline contrasts. However, the 2nd vs.1st-order relational complexity contrast showed no spatial stability, suggesting that contrast is likely not suitable for studying spatial distribution of activation, at either the channel nor ROI aggregation levels — or that a much larger amount of data would be needed to achieve stability.

#### S10.2 ROI-level Task Activation Reliability for Interleaved Split-half Analyses

In addition to the channel-level results, we computed within-session reliability across ROIs, both at the group and participant levels. At the participant level, aggregating at the ROI level yielded similar results. The Task vs. Baseline contrast showed moderate internal consistency reliability (ICC = .56) with additional variance attributable to ROI (ICC = .09), but not the interaction between ROI and participant (ICC = .00). There was substantial variation in ICC values across channels (Supplemental Figure 12), ranging from ICC = .28 - .69. On average, individual ROIs in frontal locations (M_ICC_ = .61) had significantly higher ICC values than those in parietal locations (M_ICC_ = .40; t(8) = 3.27, *p* = .01). The 2nd vs. 1st-order contrast showed comparatively lower reliability (ICC = .47), with little-to-no variance attributable to channel (ICC = .00) or the interaction between channel and subject (ICC = .00). When calculating reliability per-ROI, there was a wide range (see Supplemental Figure 12; range = .00 - .56). On average, individual ROIs in frontal locations (M_ICC_ = .16) had significantly lower ICC values than ROIs in parietal (M_ICC_ = .38) locations (t(8) = -2.47, *p* = .04).

At the group level, the Task vs. Baseline contrast demonstrated strong internal consistency (r = .89, ρ = .94, *p* < .001, variance = .87). In contrast, the 2nd vs. 1st-order contrast yielded zero reliability (r = –.31, ρ = –.91, *p* = .38, variance = .18).

**Supplementary Figure 10.**
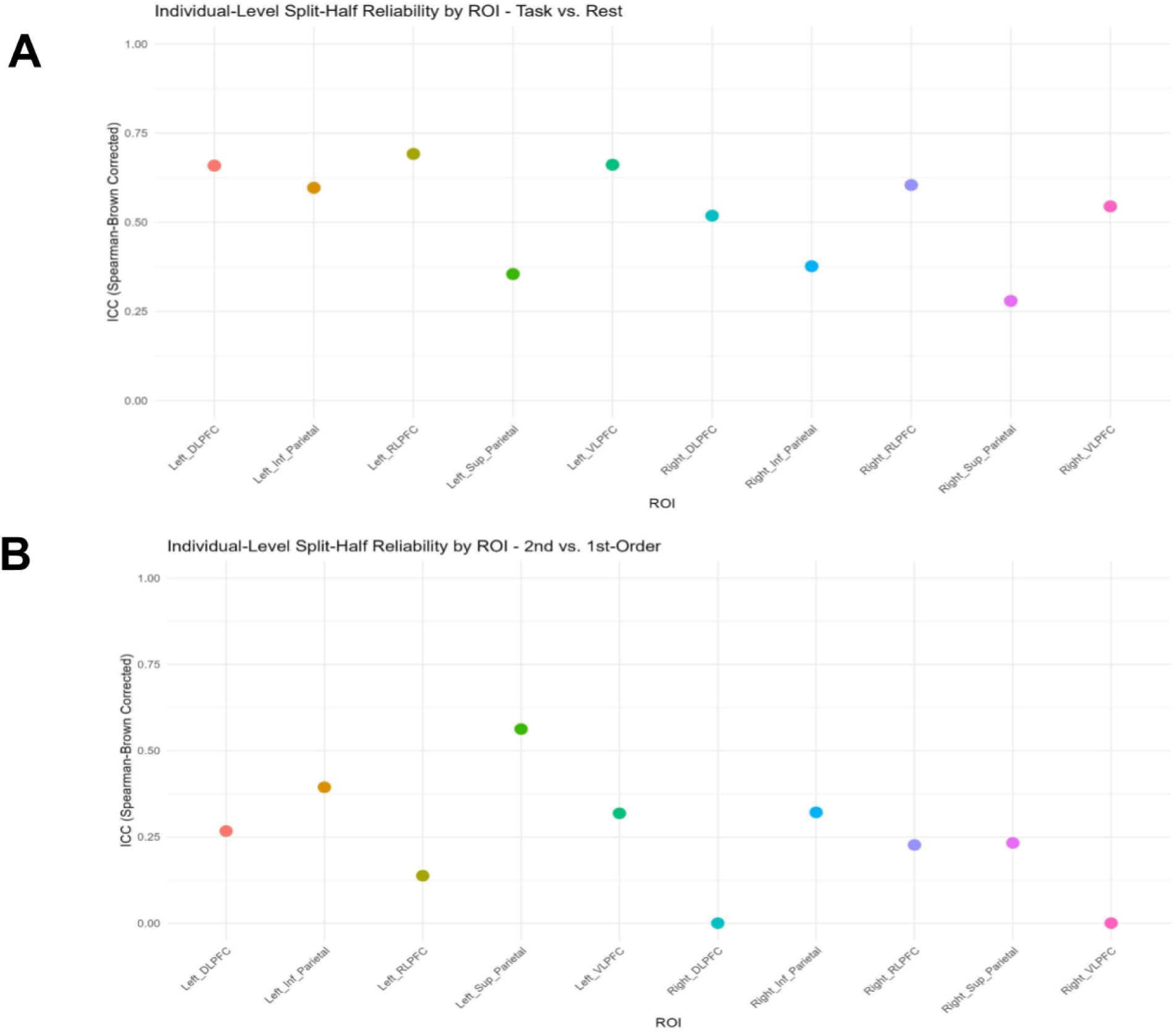
Individual level ROI ICC’s for **A.** Task vs. Baseline **B**. 2nd vs.1st-order

#### S10.3 ROI-Level Functional Connectivity Reliability for Interleaved Split-half Analyses

At the participant level, ROI-based correlations ranged from r = 0.95 to 0.99 (mean = 0.98, median = 0.99, SD = 0.01), again demonstrating strong reproducibility of individual connectivity profiles when signals are spatially aggregated. These results were consistent in the full 73-participant sample, where ROI-level correlations ranged from r = 0.96 to 0.99 (mean = 0.98, median = 0.99, SD = 0.01), with all participants exceeding r = 0.96. This pattern confirms the robustness of ROI-level connectivity as a stable and interpretable marker of within-session functional architecture.

At the group level for the subsample of 36 participants, ROI aggregation showed similarly high reliability (r = 0.998, p = 1.55 × 10^-104^) to channel-level results, confirming that broader region-based estimates preserve within-session functional network topology. These findings were further corroborated in the full sample of 73 participants, where ROI-level connectivity correlations remained highly robust (r = 0.998, p = 7.03 × 10^-109^), emphasizing the reproducibility of global network coordination within a single session.

### S11. Between-Session Reliability

#### S11.1 Behavioral Data for Session 2

Participants performed with high overall accuracy in both sessions. In Session 1, accuracy was 87.27% (SE = 0.0026) for 1st-order trials and 93.36% (SE = 0.0035) for 2nd-order trials. In Session 2, accuracy improved to 91.27% (SE = 0.0026) for 1st-order and 95.36% (SE = 0.0035) for 2nd-order trials. Reaction times consistently favored the 1st-order condition, with faster responses in both Session 1 (1.53 s vs. 2.02 s; SEs = 0.0345, 0.0559) and Session 2 (1.40 s vs. 1.71 s; SEs = 0.0345, 0.0559). Paired-sample t-tests revealed significantly faster response times in Session 2 than Session 1 (t(n–1) = 5.16, p < .001), suggesting a practice or familiarity effect. Accuracy differences were marginal (t(n–1) = –2.02, p = .054), with a trend toward improvement in Session 2.

#### S11.2 Task Activation in Sessions 1 and 2

**Supplementary Figure 11.**
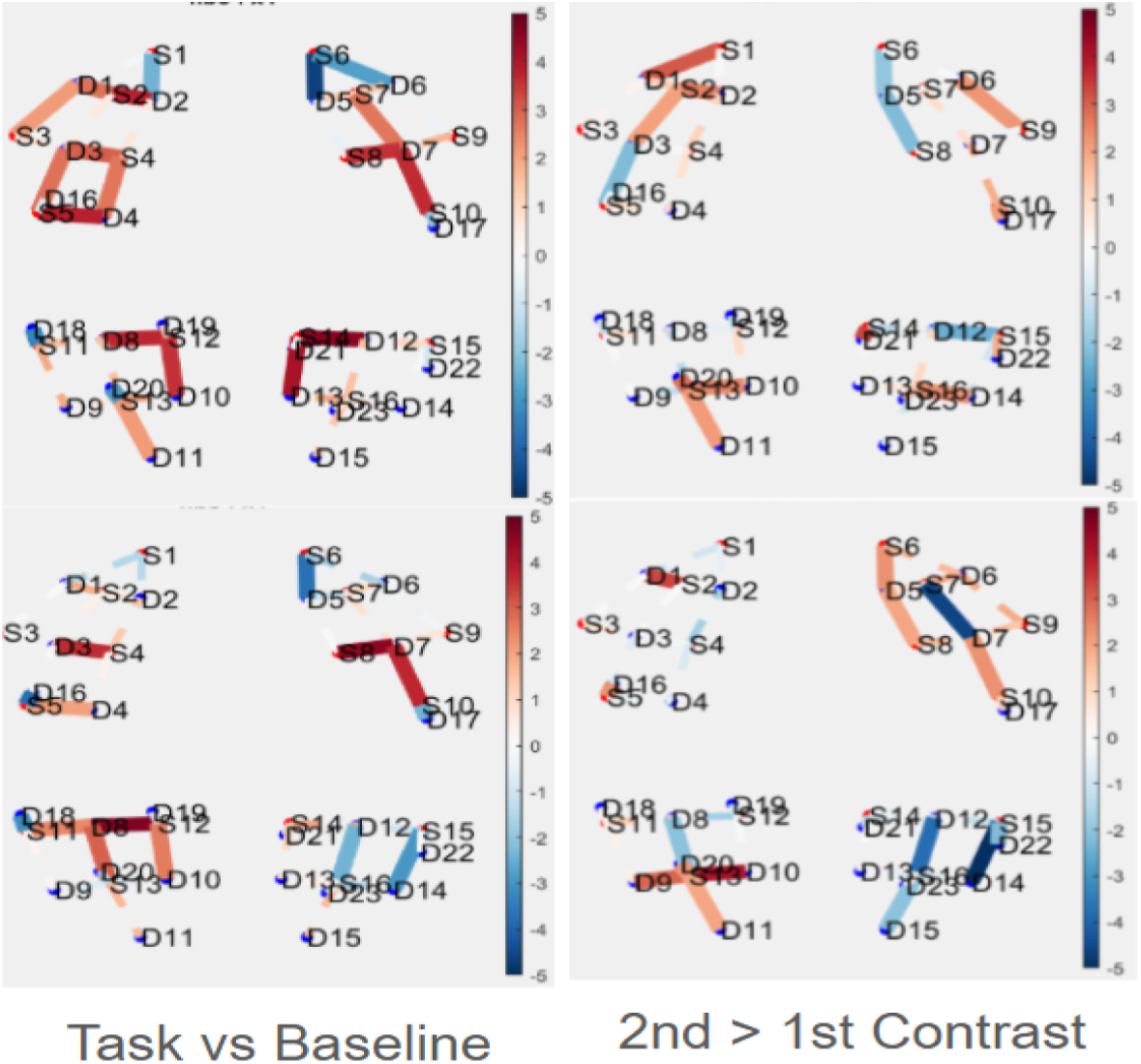
Channel x Channel group level activation plots for session 1 (top row) and session 2 (bottom row) for the task vs. baseline and 2nd vs. 1st-order contrasts.

##### S11.2.1 Task Activation Reliability at the ROI Level

At the participant level, aggregating into ROIs yielded similar results to the channel-level results. The Task vs. Baseline contrast showed relatively low test-retest reliability (ICC = .24) with little variance attributable to ROI (ICC = .01). Condition-specific contrasts showed a different pattern of test-retest reliability: 1st-order activation (ICC = .20), with no variance attributable to channel (ICC = .00), whereas 2nd-order activation (ICC = .09), with little variance attributable to channel (ICC = .03), showed noticeably lower test-retest reliability. The 2nd vs. 1st-order contrast showed similar test-retest reliability (ICC = .23) with little variance attributable to channel (ICC = .02).

At the group level, spatial consistency of activation for ROIs was higher than at the channel level for Task vs. Baseline, but remained poor for the 2nd vs. 1st-order contrast. The Task vs. Baseline contrast showed strong test–retest reliability (r = .87, p < .001), with similarly robust correlations for 1st-order (r = .72, p < .002) and 2nd-order (r = .73, p < .002). The 2nd vs. 1st-order contrast remained unreliable (r = .28, p = .44). Variance correlations were strongest for Task vs. Baseline (r = .87, p < .001) and remained high for both 1st-order (r = .76, p < .002) and 2nd-order (r = .74, p < .002), while the contrast again showed poor reliability (r = .31, p = .381). The pattern of signal amplitude differed across sessions for the Task vs.Baseline contrast (t = 3.96, p = .004) and the 2nd vs.1st-order contrast (t = 4.90, p = .001) indicating higher ROI level activity in session 1 than session 2.

**Supplementary Figure 12.**
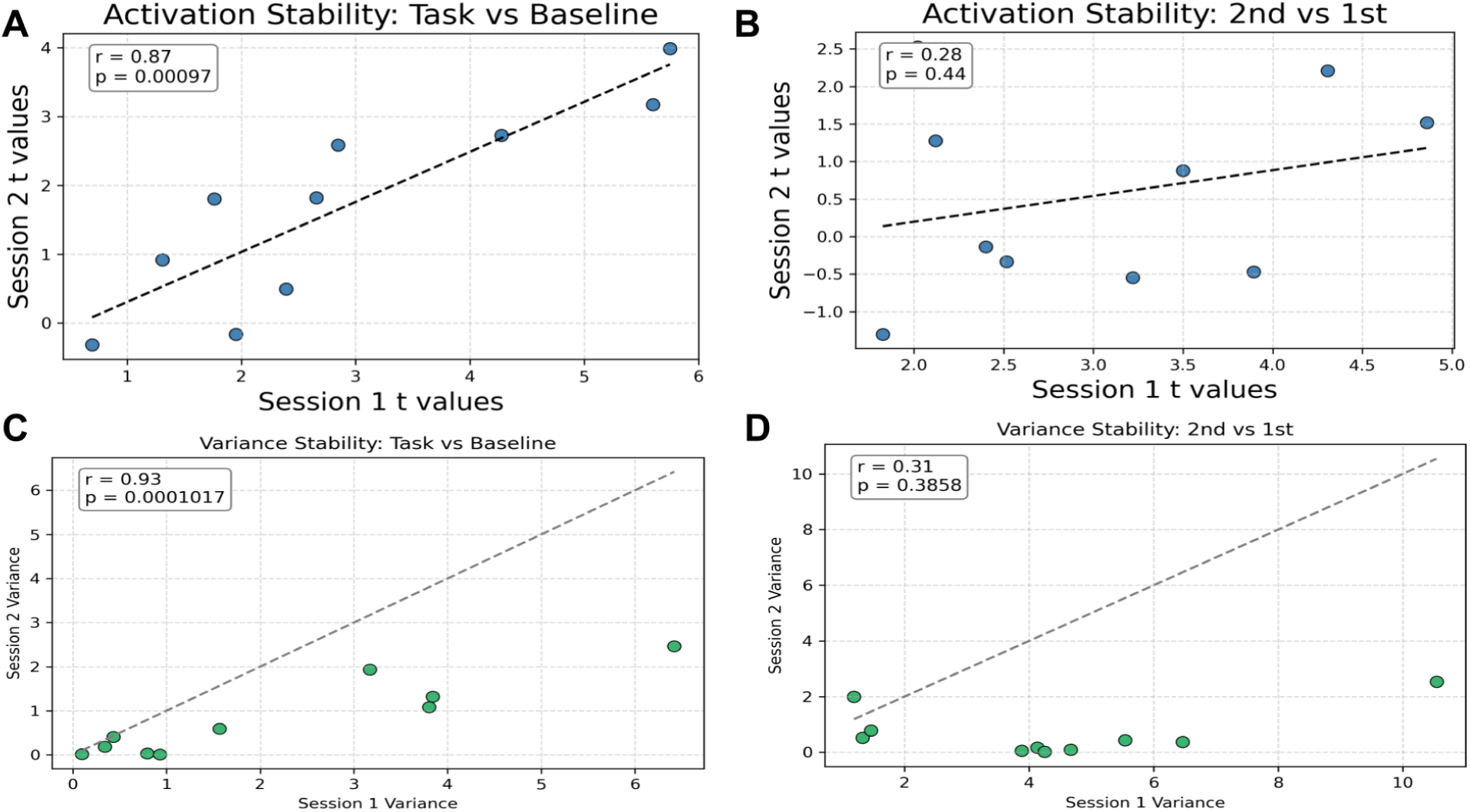
Group-level between session activation and variance stability for ROI data. **A.** Task vs. Baseline activation stability **B.** 2nd vs. 1st-order activation stability. **C.** Task vs. Baseline variance stability **D**. ROI level 2nd vs. 1st-order variance stability

#### S11.3 Functional Connectivity

To assess the test–retest reliability of group-level functional connectivity, we compared full connectivity matrices across sessions at both the channel and ROI levels on the entire between session subsample (N = 26). This analysis was also performed on the subset of participants from both sessions who had the highest quality data (<8 bad channels) and the results were almost identical. All p-values were derived using permutation testing, confirming the statistical significance of the observed stability.

##### S11.3.1 Functional Connectivity Reliability by Channel at the Group Level

At the channel level, functional connectivity patterns were highly stable across all 630 unique channel pairs. Group-level correlations between sessions were exceptionally strong for the 1st-order condition (r = 0.96, p < 1 × 10^-66^), 2nd-order condition (r = 0.95, p < 1 × 10^-66^), and Task vs. Baseline (r = 0.97, p < 1 × 10^-66^). To ensure these findings were not influenced by global signal (average correlation between all channels), we removed each participant’s unique global signal from each session and recalculated the group level functional connectivity for each session. The correlation for centered r values was also extremely strong (r = .99, p < 1 x 10^-63^), indicating this effect was not driven by global signal correlation across sessions. These findings indicate near-perfect reproducibility of global and condition-specific network structures (**Figure 13A**).

##### S11.3.2 Functional Connectivity Reliability at the ROI-Level

For participant level data at the ROI spatial scale, functional connectivity demonstrated even greater stability for individual participants than at the channel level. For 1st-order, correlations ranged from r = 0.49 to 0.98 (.89 +/- .11); for 2nd-order, from r = 0.46 to 0.97 ( .90 +/- .10); and for Task vs. Baseline, from r = 0.49 to 0.98 (.90 +/- .11). Ninety-four percent of participants showed correlations above 0.80 in all conditions, with all p-values < 0.001 (permutation test).

For group level data at the ROI spatial scale, similarly high stability was observed relative to the channel-level results. Group-level correlations of ROI-to-ROI connectivity between sessions were r = 0.98 for both the 1st-order (p < 4.50 × 10^-66^) and 2nd-order (p < 1.47 × 10^-66^) conditions, as well as for Task vs. Baseline (r = 0.98, p < 1 × 10^-66^), confirming highly reproducible patterns of large-scale network coordination across repeated measurements.

**Supplementary Figure 15.**
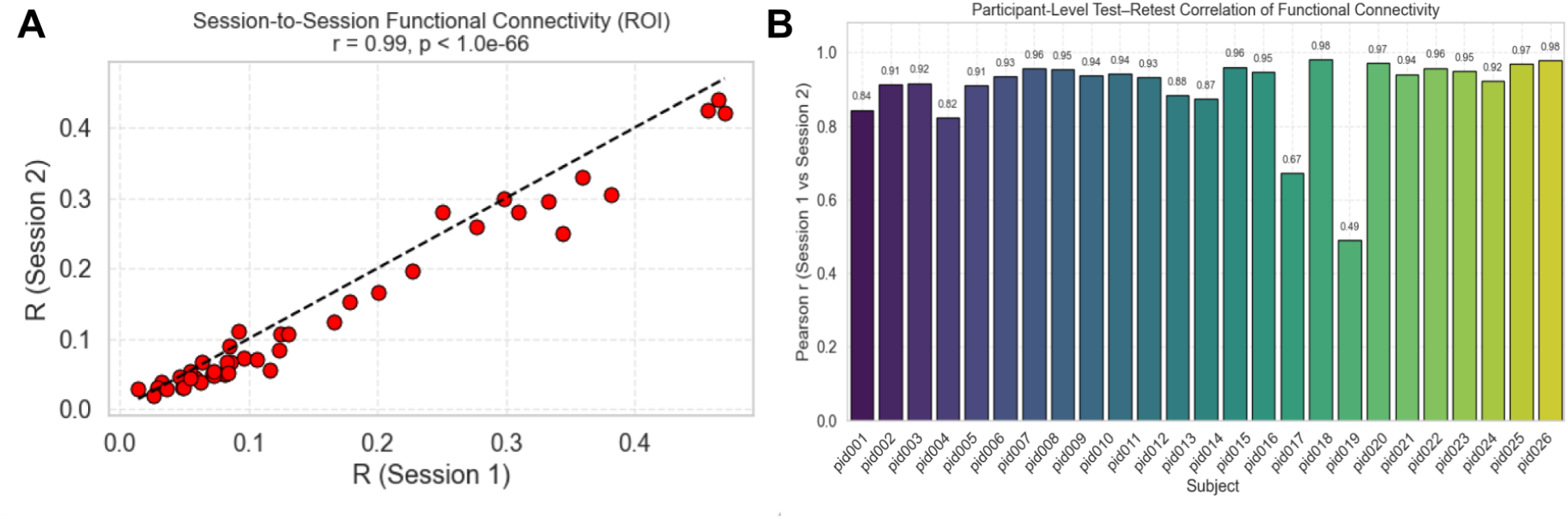
**A**. ROI group level functional connectivity correlations from session 1 to session 2 (r=.99) plotted against an identity line. **B.** ROI subject level functional connectivity correlations from session 1 to session 2 with R values ranging from .49 - .98.

### S12. Channel Quality as a Function of Hair and Skin Types

An exploratory analysis was conducted to assess whether phenotypic characteristics—specifically skin tone, hair color, hair type, and hair texture—predicted fNIRS signal quality, as in prior studies (Holmes et al., 2024; Yücel et al., 2024). Here, we operationalized signal quality as the number of unusable (bad) channels identified during preprocessing. The results of this analysis should be considered with caution, as it was not fully powered to detect main effects or interactions; a larger and even more diverse sample would be needed to measure the combined effects of skin color, hair color, and hair texture.

Participant-level metadata were compiled for 77 individuals, including manually coded phenotypic attributes and the number of bad channels calculated using the QT-NIRS module in the NIRS Toolbox. Skin tone was rated on a five-point ordinal scale based on visual inspection and video recordings, and recoded into four categories: Fair, Medium, Light Brown, and Brown. Hair color was categorized as Blonde, Brown, or Red/Black based on self-report and visual inspection. Hair type was classified by curl pattern (Straight, Wavy, Curly, Kinky), and hair texture was categorized as Fine, Medium, or Coarse. Participants with missing data for any variable were excluded.

A factorial linear model was constructed with the number of bad channels as the dependent variable and skin tone, hair color, hair type, and hair texture as fixed-effect predictors.

**Supplementary Figure 16.**
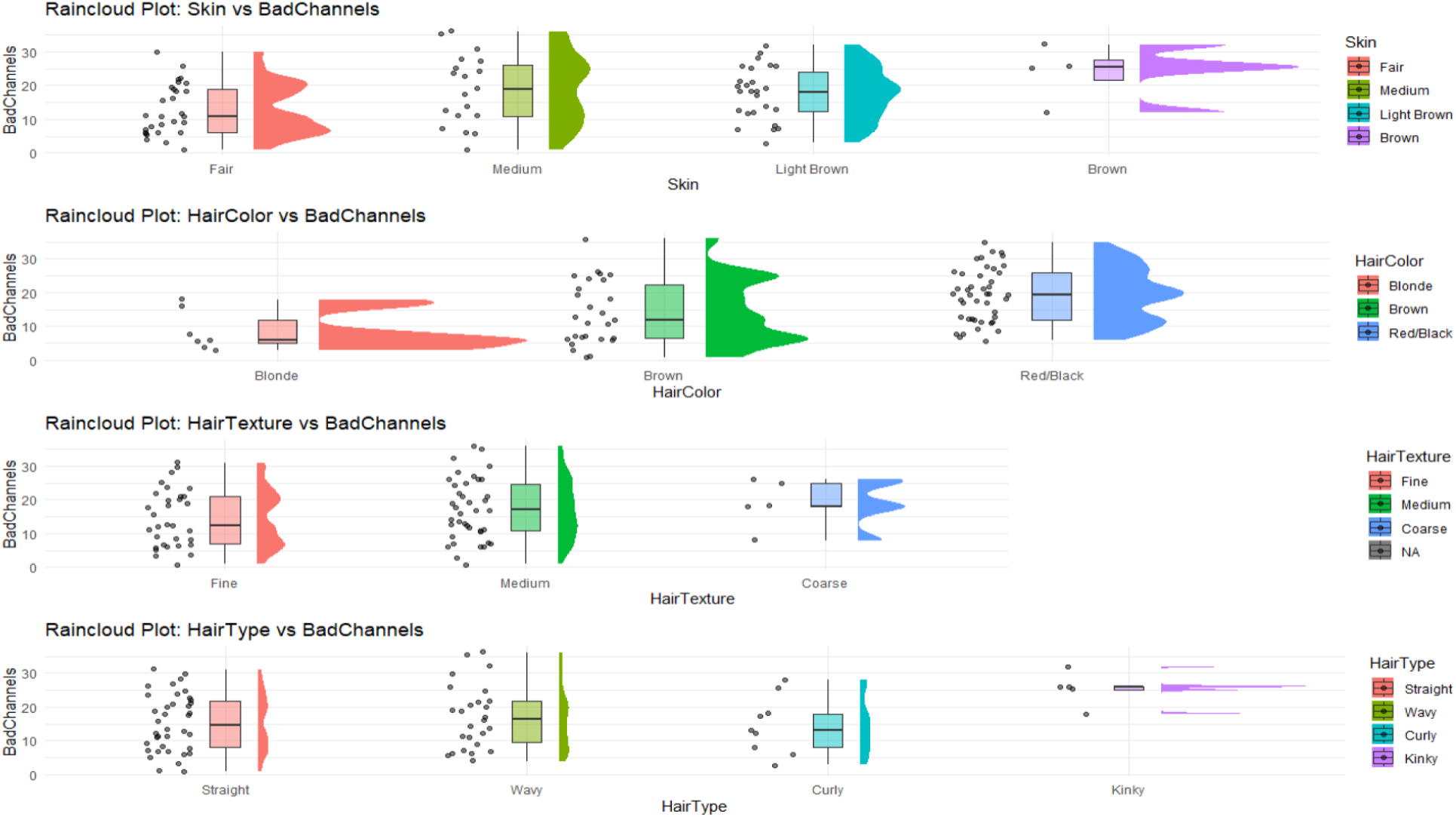
Raincloud plots of the number of bad channels as a function of skin color and hair color, texture, and type.

## Acknowledgements

This project was funded by a Jacobs Foundation Creating Impact Science Program (CRISP) fellowship (S.A.B.), an equipment award from the Berkeley Faculty Research Fund for the Biological Sciences (Professor Fei Xu and S.A.B., and University of California Berkeley internal research funds (K.J.). Research assistants Nick Zhang, Angela Ling, Nataly Lopez, Qi Guo, Kate Dellet, and Allison Chen assisted with data collection, and Julia Dil assisted with both data collection and a meta-analysis.

